# Generation of a novel *Slc7a9*^G105R^ mutant mouse identifies new biomarkers for cystinuria

**DOI:** 10.64898/2026.02.10.705194

**Authors:** Nirmal P. Bhatt, Gabriel R. Rodriguez, Giulia Iacono, Theresa T. H. Nguyen, Connor R. B. Anderson, Andrew Perry, Christopher K. Barlow, Gaetan Burgio, Simon H. Jiang, B. Med, Benjamin J. Marsland, Aniruddh V. Deshpande, Malcolm R. Starkey

## Abstract

Cystinuria is a rare inherited disease characterized by increased urinary cystine levels resulting in the formation of cystine stones in the urinary tract. Mutations in the genes encoding the cystine transporter complex, *SLC3A1* and *SLC7A9,* are the primary drivers of the disease. Current mouse models used to study cystinuria rely on gene deficiency or spontaneous mutations in mice that do not accurately reflect the pathogenic mutations found in humans. To overcome this limitation, we generated novel *Slc7a9*^G105R^ mice carrying the most common pathogenic single-point mutation in the *SLC7A9* gene. Both male and female *Slc7a9*^G105R^ mice developed a cystinuria phenotype by 9 weeks of age, characterized by substantial cystine stone formation and increased urinary cystine, lysine, arginine, and ornithine. *Slc7a9*^G105R^ mice displayed distinct serum and urinary metabolite profiles mapped to dibasic amino acid pathways. Fecal metagenomics revealed that *Slc7a9*^G105R^ mice had a heterogeneous microbiota with altered functional pathways, including increased L-cysteine biosynthesis. Depletion of the microbiota with antibiotics did not impact cystine stone burden but reduced urinary tract inflammation. Prophylactic or therapeutic dietary supplementation with alpha-lipoic acid reduced stone burden and inflammation, but it also caused damage to the urothelium. Untargeted metabolomics analysis following alpha-lipoic acid supplementation identified metabolites that can increase cystine solubility, reduce inflammation, and damage epithelial cells. Correlation analysis revealed novel serum biomarkers of stone burden, including blood urea nitrogen, 2-hydroxybutyric acid, 2-amino-2-thiazoline-4-carboxylic acid, and indole-3-acetylglycine. Collectively, the *Slc7a9*^G105R^ mutant mouse model offers a precise, rapid-onset, and translational platform for investigating cystinuria pathogenesis and evaluating potential therapeutic strategies.

**Translational statement:** The development of a novel knock-in mouse model carrying the most common pathogenic point mutation in the human *SLC7A9* gene provides a clinically relevant and translationally valuable platform for investigating cystinuria pathogenesis and testing emerging therapies. This model represents the closest possible approximation of human *SLC7A9*-mediated cystinuria, enabling rigorous preclinical evaluation of small molecules and gene therapies. It has also facilitated the identification of candidate biomarkers for cystine stone burden and treatment response, which are urgently needed to improve disease monitoring and clinical decision-making. The next critical step is to validate these biomarkers in human cystinuria cohorts to support their clinical translation.

## Introduction

Cystinuria is a rare autosomal recessive disorder characterized by defective reabsorption of cystine in the renal tubules, leading to elevated urinary cystine levels and recurrent cystine stone formation.^1–3^ This condition can result in serious complications, including urinary obstruction,^4, 5^ infections,^6^ and progressive kidney dysfunction.^7–9^ Cystinuria is primarily caused by variants in the *SLC3A1* and *SLC7A9* genes, which encode the rBAT and b^0,+^AT subunits of the heteromeric amino acid transporter responsible for the reabsorption of cystine and dibasic amino acids.^1, 2, 10^ The most common pathogenic variants include *SLC3A1* p.Met467Thr and *SLC7A9* p.Gly105Arg,^1, 2, 11^ both of which are loss-of-function alleles known to impair amino acid transport.

Despite a well-defined genetic basis, therapeutic and diagnostic options for cystinuria remain limited. Current management strategies, such as high fluid intake, urinary alkalinization, and thiol-based drugs, offer only modest benefits and are often poorly tolerated.^12, 13^ Diagnosis typically relies on stone analysis, urinary cystine quantification, genetic screening, and computed tomography (CT), the latter being the gold standard for stone detection.^14^ However, repeated CT imaging poses radiation risks, particularly in pediatric patients, underscoring the need for non-invasive biomarkers to monitor disease progression.

In mouse models, cystine stones are predominantly found in the urinary bladder and are associated with histopathological changes in both the bladder and kidney, including tubular dilation, necrosis, inflammation, and fibrosis.^15–20^ *Slc3a1* and *Slc7a9* knockout mice also exhibit impaired renal function, reflected by elevated blood urea nitrogen (BUN), serum creatinine, or reduced glomerular filtration rate.^15, 21^ These features mirror clinical observations in cystinuria patients.^10, 22^ However, existing models do not replicate the loss of function missense variants commonly seen in patients and are limited by delayed disease onset, low stone penetrance, and sex-specific differences in stone formation.^4, 17, 23^ Moreover, the lack of reliable biomarkers for disease monitoring and treatment response remains a major challenge.

Recent advances in metabolomics have revealed metabolic alterations in nephrolithiasis, particularly involving fatty acids and lipid-related pathways.^24^ Concurrently, growing evidence suggests that the gut microbiota plays a role in stone formation, particularly calcium oxalate stones, by influencing oxalate metabolism and the composition of microbial communities.^25–27^ These findings suggest that metabolomic and microbiome-based approaches may uncover novel mechanisms and biomarkers relevant to cystine stone formation and progression.

Among the emerging therapeutic candidates for cystinuria, alpha-lipoic acid (ALA), a naturally occurring antioxidant, has shown efficacy in reducing cystine stone burden in *Slc3a1*-deficient mouse models,^20^ supported by clinical case reports.^28^ A Phase II clinical trial is currently evaluating ALA in adult patients with cystinuria (NCT02910531).^29^ However, its efficacy in *Slc7a9*-based models has not been investigated, representing a critical gap in understanding genotype-specific treatment responses.

To address the limitations in the current cystinuria models, we developed a novel *Slc7a9*^G105R^ knock-in mouse model carrying a clinically relevant, patient-derived variant commonly associated with cystinuria. This is the first mouse model to incorporate a patient-specific *SLC7A9* mutation, enabling genotype-specific investigation of cystinuria. Using this model, we performed a comprehensive analysis of disease progression, including sex-specific differences, inflammatory responses, and serum and urine metabolite profiling. We also examined the role of the gut microbiota in cystine stone formation and evaluated the therapeutic potential of ALA. Notably, BUN and three metabolites, 2-hydroxybutyric acid, 2-amino-2-thiazoline-4-carboxylic acid, and indole-3-acetylglycine, emerged as promising candidate biomarkers.

Collectively, these findings establish the *Slc7a9*^G105R^ model as a clinically relevant platform for advancing our understanding of cystinuria and accelerating the development of targeted diagnostics and therapies.

## Methods

*Detailed methods, including animal conditions, mutant mouse generation, genotyping, diets, tissue collection, kidney function assessment, spironolactone administration, gene expression analysis, proteomics, metabolomics, antibiotic administration, metagenomics, cohousing, and detailed statistical analyses, are provided in the online supplementary material*.

### Micro-computed tomography

µ-CT imaging of mouse urinary bladder cystine stones was performed *ex vivo* using a NanoScan PET/CT machine (Mediso, Budapest, Hungary) with the following parameters: tube voltage, 70 kVp; tube current, 680 μA; exposure time, 300 ms; 1440 projections, 1 rotation; volume size: 22 x 22 x 12 μm; voxel size (reconstructed), 11 μm^3^.

### Histology

Transverse sections of paraffin-embedded mouse urinary bladder (4 μm) were stained with hematoxylin and eosin (H&E). Images were taken on the 4x objective, and magnified structures were further imaged on the 40x objective.

### Amino acid quantification

Arginine, cystine, lysine, and ornithine were quantified in mouse serum and urine, and compared against external calibration curves by Liquid Chromatography-Mass Spectrometry (LC-MS) using a Dionex RSLC3000 UHPLC coupled to a Q-Exactive Plus Orbitrap MS (Thermo-Fisher Scientific, North Ryde, VIC, Australia). The peak area corresponding to the extracted ion chromatograms of the protonated amino acids was integrated using TraceFinder 4.1 (Thermo-Fisher Scientific).

### Untargeted metabolomics

Mouse serum and urine metabolites were extracted using a 4:1 ratio of methanol containing 1 μM CCTP internal standards, 5 μM BHT and 13C,15N-amino acid mixture. Urine volume was adjusted to creatinine levels. Samples were acquired using a Q-Exactive Orbitrap mass spectrometer (Thermo-Fisher). LC-MS data was processed using the metabolome-lipidome-MSDIAL pipeline (https://zenodo.org/records/15701972). Feature peaks were detected using MS-DIAL (version 5.3) and matched against the MassBank database (September 2024). R (version 4.4.2) was used with the pmp R package (version 1.18.0). Features were further annotated against the Human Metabolome Database (HMDB, version 5). Limma (version 3.62.2) was used to detect differentially abundant metabolites between groups (False Discovery Rate (FDR) < 0.05, |LogFC| > 0.5). Hmisc (version 5.2.3) was used for Pearson correlations (*P* < 0.05). Features were mapped against metabolic pathways within the KEGG database.

### Fecal metagenomics

Mouse fecal DNA samples were quantified and quality-checked using the Invitrogen Qubit system (Invitrogen, USA). Sample integrity was verified with the Agilent Bioanalyzer 2100 microfluidics system (Agilent Technologies, Germany) before constructing Illumina NexteraXT sequencing libraries (Illumina Inc., USA). Libraries were sequenced on an MGI DNBSEQ G400 (MGI Tech Co., Ltd, China). Trimmed, quality-filtered, host-depleted paired-end reads were used to estimate taxonomic abundances for each sample with the MetaPhlAn4 software.^30^ Differentially abundant species were determined using ANCOM-BC2.^31^ Enriched functional pathways in the MetaCyc database^32^ were quantified using HUMAnN3,^33^ and differential pathway abundance was estimated using MaAsLin2.^34^

### Study approval

The Australian National University Animal Ethics Committee (A2017/44) approved the generation of the *Slc7a9*^G105R^ mice. Experimental studies were approved by the Alfred Research Alliance Animal Ethics Committee (E/2074/2021/M) in accordance with the National Health and Medical Research Council’s Australian Code of Practice. All experiments are in accordance with the ARRIVE guidelines (Animal Research: Reporting of *in vivo* experiments).

### Statistical analyses

Statistical analyses were performed using GraphPad Prism (version 9.0.2). Grubb’s test was applied to all data sets. A maximum of one statistical outlier per experimental group was identified using Grubb’s test with a significance level (alpha) 0.05. The unpaired non-parametric Mann-Whitney test was used to compare two groups. A two-way ANOVA followed by Tukey’s multiple comparisons test was used to analyze differences across multiple time points. Statistical significance was set at a 95% confidence interval or *P* value < 0.05. Statistical models for metabolomics and metagenomics are described in the respective figure legends.

## Results

### Homozygote *Slc7a9*^G105R^ mice develop substantial cystine stones by 9 weeks

To characterize our novel mouse model of cystinuria that mimics the most common pathogenic single nucleotide variant in human *SLC7A9* (**Supplementary Figure S1**, **Supplementary Table S1**), we placed homozygote *Slc7a9*^G105R^ mice on a cysteine-controlled diet (**Supplementary Table S2**) at 3 weeks of age, and body weight, kidney function, cystine stone formation, and histopathology were assessed over time (**Figure 1a**).

**Figure 1.**
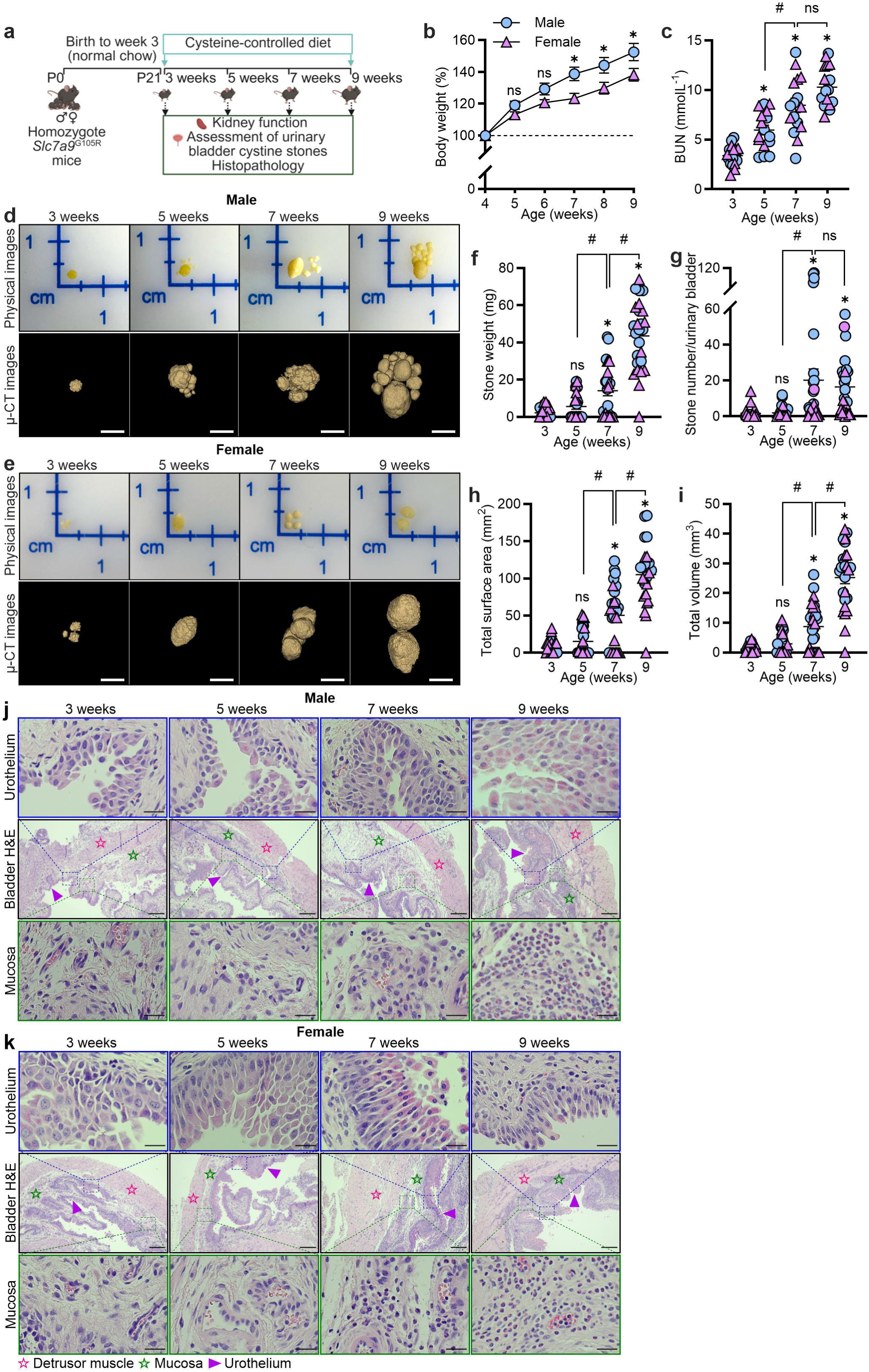
Homozygote *Slc7a9*^G105R^ mice develop substantial cystine stones by 9 weeks. Kidney function, urinary bladder cystine stone growth and histopathology were assessed at 3, 5, 7, and 9 weeks. (**a**) Schematic representation of the experimental model. (**b**) Percentage body weight change in male (blue; *n* = 13) and female (pink; *n* = 13) homozygote *Slc7a9*^G105R^ mice over time. (**c**) Serum blood urea nitrogen (BUN) levels at 3, 5, 7, and 9 weeks in males (*n* = 8-10) and females (*n* = 8). Representative matched physical and µ-CT urinary bladder cystine stone images from a single mouse at 3, 5, 7, and 9 weeks for (**d**) males and (**e**) females. Urinary bladder cystine stone (**f**) weight, (**g**) number, (**h**) total surface area, and (**i**) total volume in males (*n* = 11-13) and females (*n* = 11-14) at 3, 5, 7, and 9 weeks, obtained from *ex vivo* µ-CT imaging. Representative images of hematoxylin and eosin (H&E)-stained transverse sections (4 µm) of the urinary bladder at 3, 5, 7, and 9 weeks in (**j**) males and (**k**) females. The scale bar indicates 2.5 mm for the representative urinary bladder cystine stone µ-CT images. Ill, Ill, and ► denote the detrusor muscle, mucosa, and urothelium, respectively. The scale bar indicates 2.5 mm for urinary cystine stones, 200 µm for the bladder H&E section, and 25 µm for the magnified sections of urothelium and mucosa, respectively. The data presented were obtained from two independent experiments. Data are presented as the mean ± the standard error of the mean (SEM). Statistical differences were calculated using a two-way ANOVA with a Tukey multiple comparison test (**b**) and a non-parametric unpaired Mann-Whitney t-test (**c**, **f**-**i**). *P* value < 0.05. * denotes a statistical difference compared to females (**b**). * also denotes a statistical difference compared to 3 weeks (**c**, **f**-**i**). ^#^ denotes statistical differences between two consecutive time points. ns denotes not significant.

Male homozygote *Slc7a9*^G105R^ mice had significantly greater body weight than females at weeks 7, 8, and 9 (**Figure 1b**). BUN, a surrogate marker of kidney function, significantly increased with stone burden over time, peaking at 9 weeks (**Figure 1c**). Representative matched physical and µ-CT images visually demonstrate that urinary bladder cystine stone growth progressed markedly from weeks 3 to 9 in both males and females (**Figure 1d**-**e**). Notably, µ-CT imaging revealed no cystine stones in the kidneys of homozygote *Slc7a9*^G105R^ mice at 9 weeks of age (**Supplementary Figure S2**).

Cystine stone weight, number, surface area, and volume significantly increased at weeks 7 and 9 compared to week 3 (**Figure 1f-i**). Cystine stone weight, number, surface area, and volume significantly increased from week 5 to 7, and all but stone number increased from week 7 to 9 (**Figure 1f**-**i**).

BUN was positively correlated with cystine stone weight, surface area, and volume in both male and female homozygote *Slc7a9*^G105R^ mice (**Supplementary Figure S3a**, **c-e**, **g-h**). However, BUN only showed a positive correlation with cystine stone number in males, but not in females (**Supplementary Figure S3b**, **f).**

In male homozygote *Slc7a9*^G105R^ mice, physical cystine stone weight positively correlated with cystine stone number, area, and volume (**Supplementary Figure S4a-c**). In female homozygote *Slc7a9*^G105R^ mice, physical cystine stone weight positively correlated with cystine stone area and volume, but not number (**Supplementary Figure S4d-f**). Collectively, this indicates that the µ-CT imaging accurately captures actual cystine stones.

The urinary bladder from homozygote *Slc7a9*^G105R^ mice displayed profound histopathology, characterized by substantial mucosal inflammation and hyperplasia/metaplasia of the urothelium in both male and female mice (**Figure 1j**-**k**, **Supplementary Figure S5**). There was also notable inflammation present within the detrusor muscle of the urinary bladder.

BUN, cystine stone weight, surface area, and volume, as well as urinary bladder histopathology throughout the time course, were similar between male and female homozygote *Slc7a9*^G105R^ mice. The exception was cystine stone number, which was significantly increased in male *Slc7a9*^G105R^ mice at weeks 7 and 9 compared to age-matched females (**Supplementary Figure S6**). Suppression of testosterone with spironolactone significantly reduced serum testosterone levels and cystine stone number but did not alter BUN, cystine stone weight, total surface area, total volume, or urinary bladder histopathology in male homozygote *Slc7a9*^G105R^ mice (**Supplementary Figure S7**).

Notably, there was minimal progression of cystine stone burden in homozygote *Slc7a9*^G105R^ mice from 9 to 12 weeks, with no change in cystine stone weight or volume but a modest increase in stone number and surface area (**Supplementary Figure S8**). Therefore, week 9 was chosen as the ideal time for further characterization.

### Homozygote *Slc7a9*^G105R^ mice have increased blood urea nitrogen, bladder histopathology, and urinary levels of arginine, cysteine, lysine, and ornithine

Hallmark features of cystinuria in homozygote *Slc7a9*^G105R^ mice were compared to wild-type and heterozygote controls at week 9 (**Figure 2a**). Increased body weight in males was not genotype-dependent (**Figure 2b**). Homozygote *Slc7a9*^G105R^ mice had significantly increased BUN compared to wild-type or heterozygote controls (**Figure 2c**). BUN levels were significantly decreased in heterozygote *Slc7a9*^G105R^ mice compared to wild-type controls (**Figure 2c**). *In vivo* assessment of kidney function using real-time transdermal glomerular filtration rate (*t*GFR) showed no change in homozygote *Slc7a9*^G105R^ mice compared to wild-type or heterozygote controls (**Supplementary Figure S9**). Therefore, *t*GFR was not assessed in subsequent experiments.

**Figure 2.**
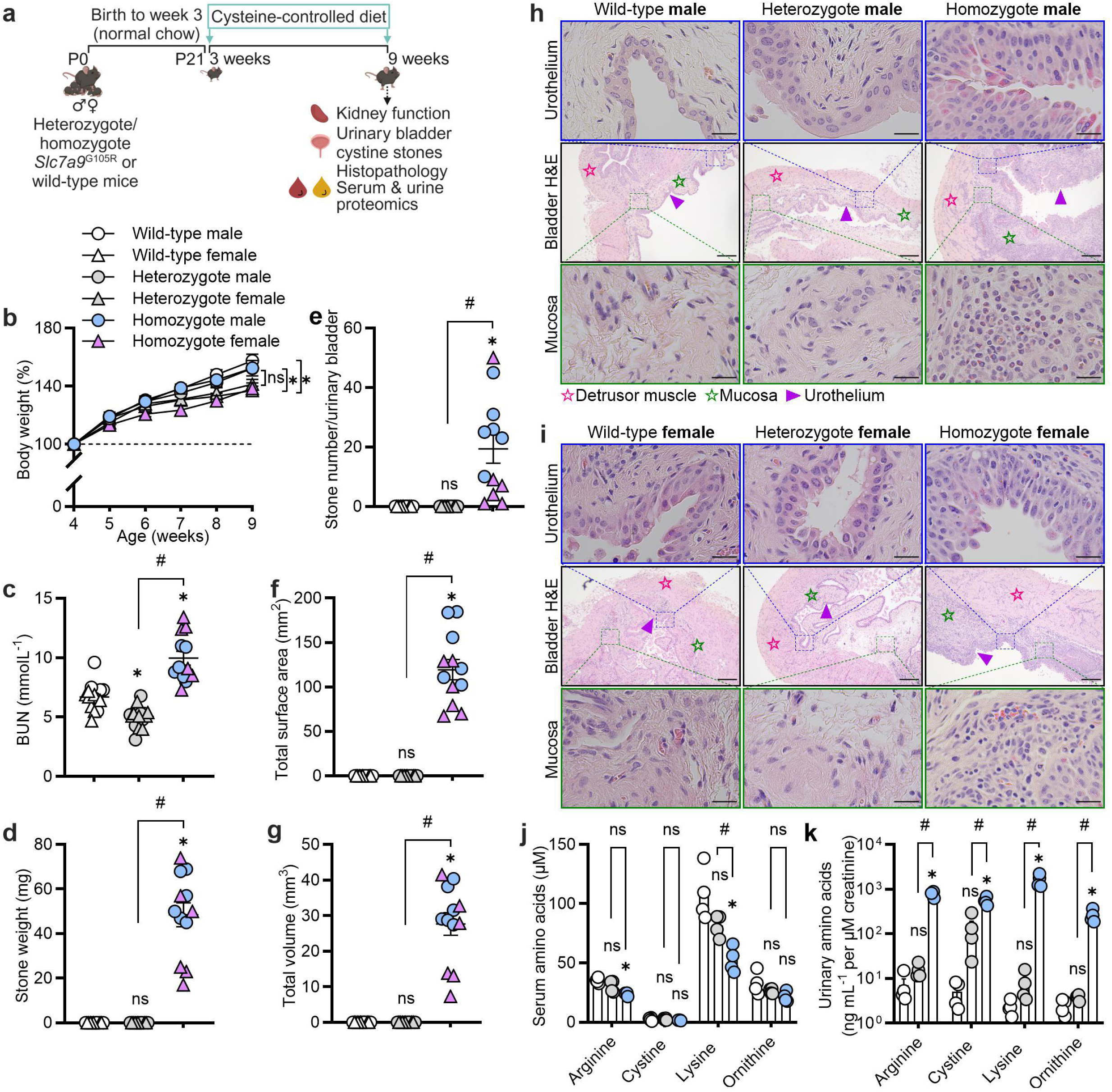
Homozygote *Slc7a9*^G105R^ mice have increased blood urea nitrogen, bladder histopathology, and urinary levels of arginine, cystine, lysine, and ornithine. Kidney function, cystine stone formation, urinary bladder histopathology, and cystine and dibasic amino acid levels were assessed in 9-week-old homozygote *Slc7a9*^G105R^ mice compared to age and sex-matched heterozygote *Slc7a9*^G105R^ and wild-type controls. (**a**) Schematic representation of the experimental model. (**b**) Percentage body weight change in male and female wild-type, heterozygote *Slc7a9*^G105R^, and homozygote *Slc7a9*^G105R^ mice over time (*n* = 13 males and *n* = 13 females per group). (**c**) Serum blood urea nitrogen (BUN) levels at week 9 (*n* = 6 males and *n* = 6 females per group). Urinary bladder cystine stone (**d**) weight, (**e**) number, (**f**) total surface area, and (**g**) total volume in males (*n* = 6) and females (*n* = 6). Representative images of hematoxylin and eosin (H&E)-stained transverse sections (4 µm) of the urinary bladder in (**h**) males and (**i**) females. Levels of cystine and the dibasic amino acids—arginine, lysine, and ornithine—in (**j**) serum (*n* = 4) and (**k**) urine (*n* = 4). Ill, Ill, and ► denote the detrusor muscle, mucosa, and urothelium, respectively. The scale bar indicates 200 µm for the bladder H&E section and 25 µm for the magnified sections of urothelium and mucosa, respectively. Data were obtained from two independent experiments. Data are presented as the mean ± the standard error of the mean (SEM). Statistical significance was calculated using a two-way ANOVA with a Tukey multiple comparisons test (**b**) or a non-parametric unpaired Mann-Whitney t-test (**c**-**g** and **j**-**k**). * denotes a statistical difference compared to wild-type and heterozygote controls (**b**). * also denotes a statistical difference compared to wild-type controls (**c**-**g** and **j**-**k**). ^#^ denotes statistical differences between two consecutive groups. ns denotes not significant.

Cystine stone weight, number, surface area, and volume were significantly increased in homozygote *Slc7a9*^G105R^ mice compared to wild-type and heterozygote controls (**Figure 2d**-**g**). The urinary bladder from homozygote *Slc7a9*^G105R^ mice displayed profound histopathology, characterized by substantial mucosal inflammation and hyperplasia/metaplasia of the urothelium compared to wild-type or heterozygote controls (**Figure 2h**-**i**, **Supplementary Figure S10**). Occasional kidney histopathology, including inflammatory cell infiltrate, tubular dilation, and cast formation, was observed in homozygous *Slc7a9*^G105R^ mice, which was absent in wild-type or heterozygous controls (**Supplementary Figure S11**). Homozygote *Slc7a9*^G105R^ mice had a modest increase in kidney proinflammatory gene expression, including *Cxcl1, Cxcl2, Ccl2,* and *Il6,* compared to heterozygote controls (**Supplementary Figure S12**, **Supplementary Table S3**-**4**).

Serum arginine and lysine, but not cystine and ornithine, were decreased in homozygote *Slc7a9*^G105R^ mice compared to wild-type controls (**Figure 2j**). Homozygote *Slc7a9*^G105R^ mice had decreased serum lysine compared to heterozygote controls (**Figure 2j**). Urinary arginine, cystine, lysine, and ornithine levels increased in homozygote *Slc7a9*^G105R^ mice compared to wild-type or heterozygote controls (**Figure 2k**). Cystine and all dibasic amino acid levels in the serum and urine were comparable between heterozygote and wild-type controls (**Figure 2j**-**k**). Homozygote *Slc7a9*^G105R^ mice also display classical hexagonal cystine crystals in their urine that were absent in the urine of wild-type and heterozygote controls (**Supplementary Figure S13**).

Collectively, the cystinuria phenotype was similar in both males and females (**Figure 1-2**). Therefore, subsequent experiments used males only to reduce animal numbers and comply with animal ethics requirements. Additionally, wild-type controls were comparable to heterozygote controls (**Figure 2**), so subsequent experiments used heterozygote controls only.

### Homozygote *Slc7a9*^G105R^ mice display distinct serum and urinary metabolite profiles

Untargeted serum and urine metabolomics were performed at week 9 (**Figure 3a**). Homozygote *Slc7a9*^G105R^ mice displayed significantly different serum and urine metabolite profiles compared to heterozygote controls (**Figure 3b**-**c**). Differentially abundant metabolites were mapped to relevant metabolic pathways (**Figure 3d**, **Supplementary Figure S14a**-**b**) and metabolite interactions for both serum and urine (**Supplementary Figure S14c**). There were more differentially abundant metabolites in the urine than in serum, with most mapping to arginine, cysteine, and lysine pathways (**Figure 3d**).

**Figure 3.**
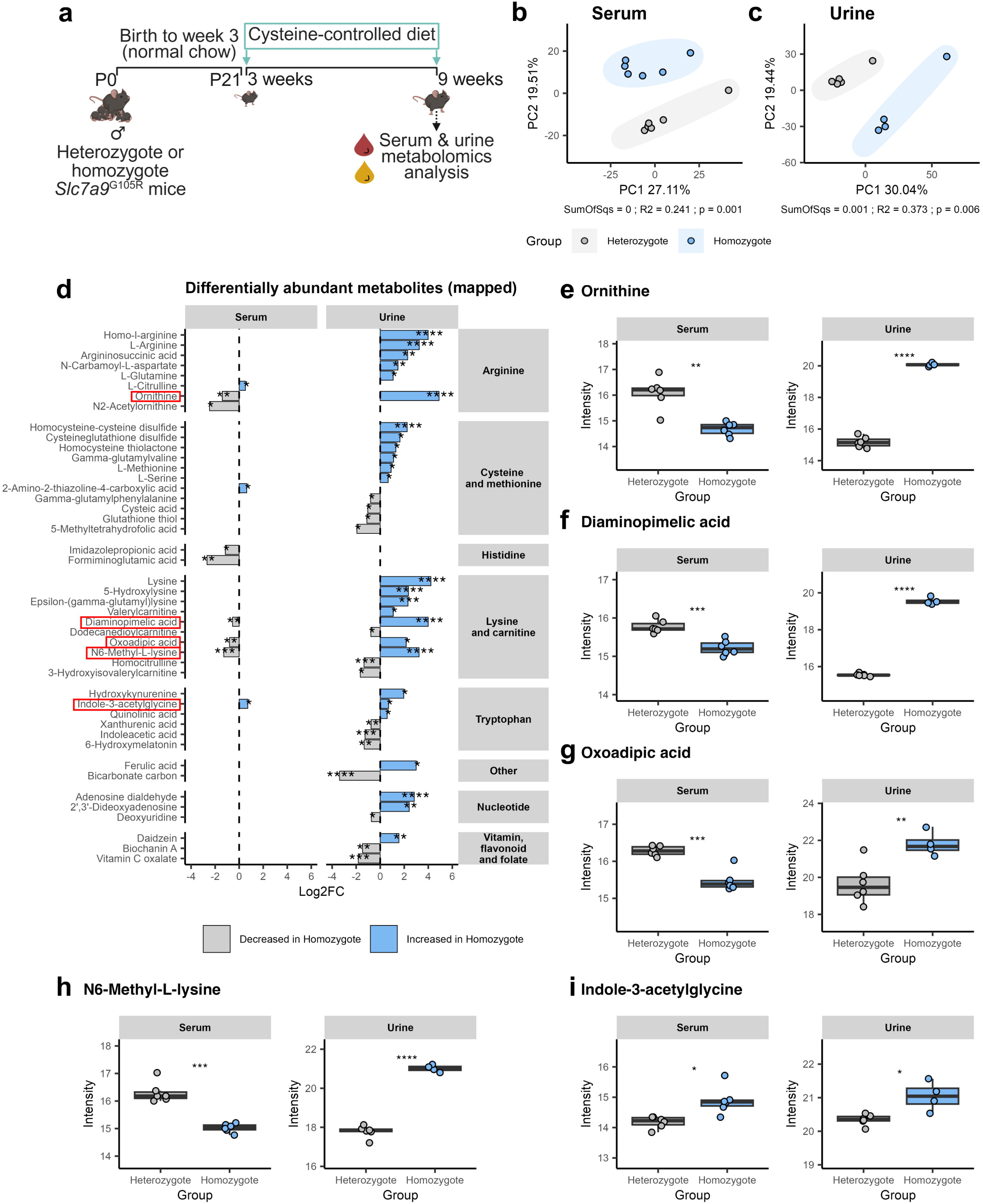
Homozygote *Slc7a9*^G105R^ mice display distinct serum and urinary metabolite profiles. Serum and urine were collected from male 9-week-old heterozygote and homozygote *Slc7a9*^G105R^ mice for untargeted metabolomics analysis. (**a**) Schematic representation of the experimental model. (**b**) PCA plot for serum metabolites. Ellipses represent the 95% confidence interval around the group centroid. PERMANOVA test results for the group. SumOfSqs (sum of squares): effect size; R2: variance explained, p: *P* value. (**c**) PCA plot for urine metabolites. (**d**) Differentially abundant serum (left) and urine (right) metabolites mapped to metabolic pathways (Log2FC > 0.5, FDR < 0.05). Box plots showing common metabolites in the serum (left) and urine (right), including (**e**) ornithine, (**f**) diaminopimelic acid, (**g**) oxoadipic acid, (**h**) N6-methyl-L-lysine, and (**i**) indole-3-acetylglycine. Serum samples: *n* = 6 per group. Urine samples: *n* = 6 for heterozygote and *n* = 4 for homozygote groups. Data are presented as the median, interquartile range (IQR) (boxes) and 1.5x IQR (whiskers). Statistical significance between homozygote and heterozygote *Slc7a9*^G105R^ mice was calculated using a Student’s t-test; *, **, ***, and **** denote *P* < 0.05, 0.01, 0.001, and 0.0001, respectively.

Of particular interest were the differentially abundant metabolites that mapped to the cysteine and methionine pathways. In urine, homocysteine-cysteine disulfide, cysteine glutathione disulfide, homocysteine thiolactone, gamma-glutamylvaline, L-methionine, and L-serine were significantly increased, whilst gamma-glutamylphenylalanine, cysteic acid, glutathione thiol, and 5-methyltetrahydrofolic acid were significantly decreased (**Figure 3d**). 2-Amino-2-thiazoline-4-carboxylic acid was the only differentially abundant serum metabolite mapped to the cysteine and methionine pathway (**Figure 3d**). This untargeted metabolomics approach also confirmed increased urinary levels of the dibasic amino acids L-arginine, ornithine, and lysine (**Figure 3d**).

Four metabolites were decreased in serum but increased in urine, including ornithine, diaminopimelic acid, oxoadipic acid, and N6-methyl-L-lysine (**Figure 3e**-**h**). Indole-3-acetylglycine was the only metabolite increased in serum and urine (**Figure 3i)**.

### Fecal microbiota and functional pathways are altered in homozygote *Slc7a9*^G105R^ mice

The gastrointestinal microbiota influences the severity of other forms of lithiasis,^25^ and the *Slc7a9* gene is also expressed in the apical membrane of epithelial cells of the small intestine.^35^ To investigate if the G105R mutation in *Slc7a9* alters the gastrointestinal microbiome, fecal pellets were collected from individual mice at 9 weeks for metagenomics (**Figure 4a**).

**Figure 4.**
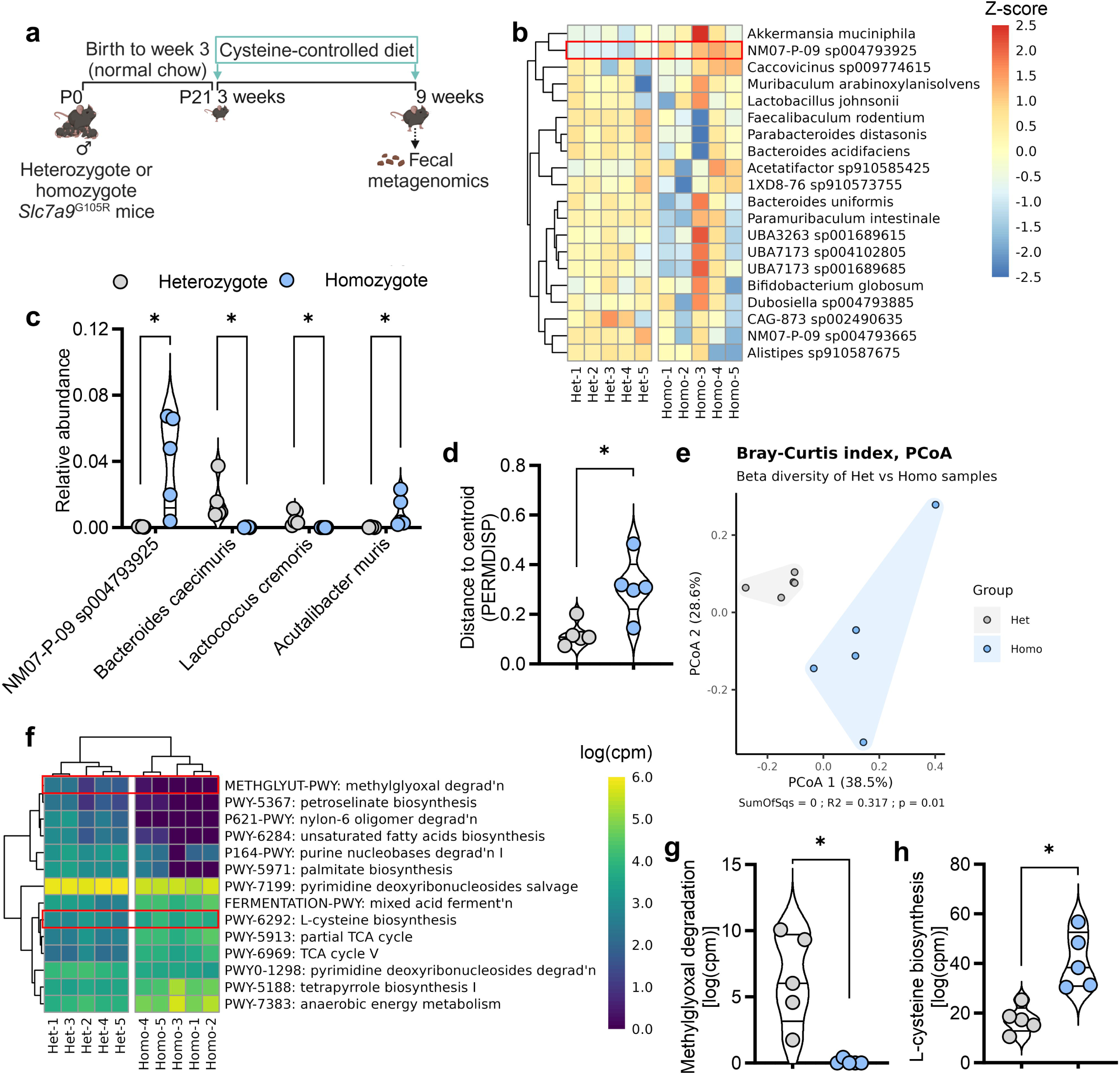
Fecal microbiota and functional pathways are altered in homozygote *Slc7a9*^G105R^ mice. Fecal samples were collected from male heterozygote and homozygote *Slc7a9*^G105R^ mice at 9 weeks, and the microbiome was assessed using metagenomics. (**a**) Schematic representation of the experimental model. (**b**) Heatmap of the 20 most abundant taxa by species. (**c**) Relative abundance of significantly different species (FDR ≤ 0.05). (**d**) Group heterogeneity is the distance from the centroid of each group in the Bray-Curtis index Principle Coordinate Analysis (PCoA). (**e**) A PCoA plot of the Bray-Curtis dissimilarity index. (**f**) Heatmap showing the relative copies of the significantly different functional pathways (FDR ≤ 0.05). (**g**-**h**) Violin plot showing pathway copies per million for the methylglyoxal degradation and the L-cysteine biosynthesis pathways. All data are *n* = 5 per group. Data are presented as the mean ± the standard error of the mean (SEM). Z-scores of the centered log-ratio (CLR)-transformed relative abundances were used (**b**). Data are presented as the mean ± the standard error of the mean (SEM). Statistical differences were calculated using a non-parametric unpaired Mann-Whitney t-test (**c**-**d**, **g**-**h**). * indicates *P* value < 0.05.

Assessment of the top 20 abundant species in individual heterozygote and homozygote *Slc7a9*^G105R^ mice indicated heterogeneity in the fecal microbiome (**Figure 4b**, **Supplementary Figure S15a**). The diversity index (Shannon) at the species level was comparable in homozygotes and heterozygotes (**Supplementary Figure S15b**). The relative abundance of NM07-P-09 sp004793925 (also known as *Leptogranulimonas caecicola*)^36^ and *Acutalibacter muris* were significantly increased, whilst *Bacteroides caecimuris* and *Lactococcus cremoris*^37^ were significantly decreased in homozygote *Slc7a9*^G105R^ mice compared to heterozygote controls (**Figure 4c**). The Bray-Curtis index was used to assess group heterogeneity and identified a significant increase in microbiome heterogeneity in homozygotes compared to heterozygotes (**Figure 4d**-**e**).

Further pathway analysis revealed 14 significantly altered pathways in the gut microbial community of homozygote *Slc7a9*^G105R^ mice compared to heterozygote controls (**Figure 4f**, **Supplementary Figure S16a**-**l**). The METHGLYUT-PWY: methylglyoxal degradation pathway was significantly reduced, and PWY-6292: L-cysteine biosynthesis pathway was significantly increased in homozygote *Slc7a9*^G105R^ mice compared to heterozygote controls (**Figure 4g**-**h**). The other 12 significantly altered pathways, with no known relevance to cysteine metabolism, included TCA cycle V, partial TCA cycle, tetrapyrrole biosynthesis I, anaerobic energy metabolism, mixed acid fermentation, pyrimidine deoxyribonucleosides salvage, unsaturated fatty acids biosynthesis, petroselinate biosynthesis, palmitate biosynthesis, pyrimidine deoxyribonucleosides degradation, purine nucleobases degradation I, and nylon-6 oligomer degradation (**Figure 4f**, **Supplementary Figure S16a**-**l**).

### Prophylactic antibiotic treatment reduced fecal microbiome diversity and urinary tract inflammation but did not affect kidney function or urinary bladder cystine stone growth

Given the changes in the fecal microbiome, we hypothesized that depletion of the microbiome may alter cystine stone growth. To achieve this, mice were treated with antibiotics from weeks 3 to 9 (**Figure 5a**). Antibiotic-treated homozygote *Slc7a9*^G105R^ mice had similar body weight, but significantly reduced microbiome diversity compared to vehicle-treated controls (**Figure 5b**-**d**). Serum BUN, stone appearance, weight, number, surface area, and volume were unaltered compared to vehicle-treated controls (**Figure 5e**-**j**).

**Figure 5.**
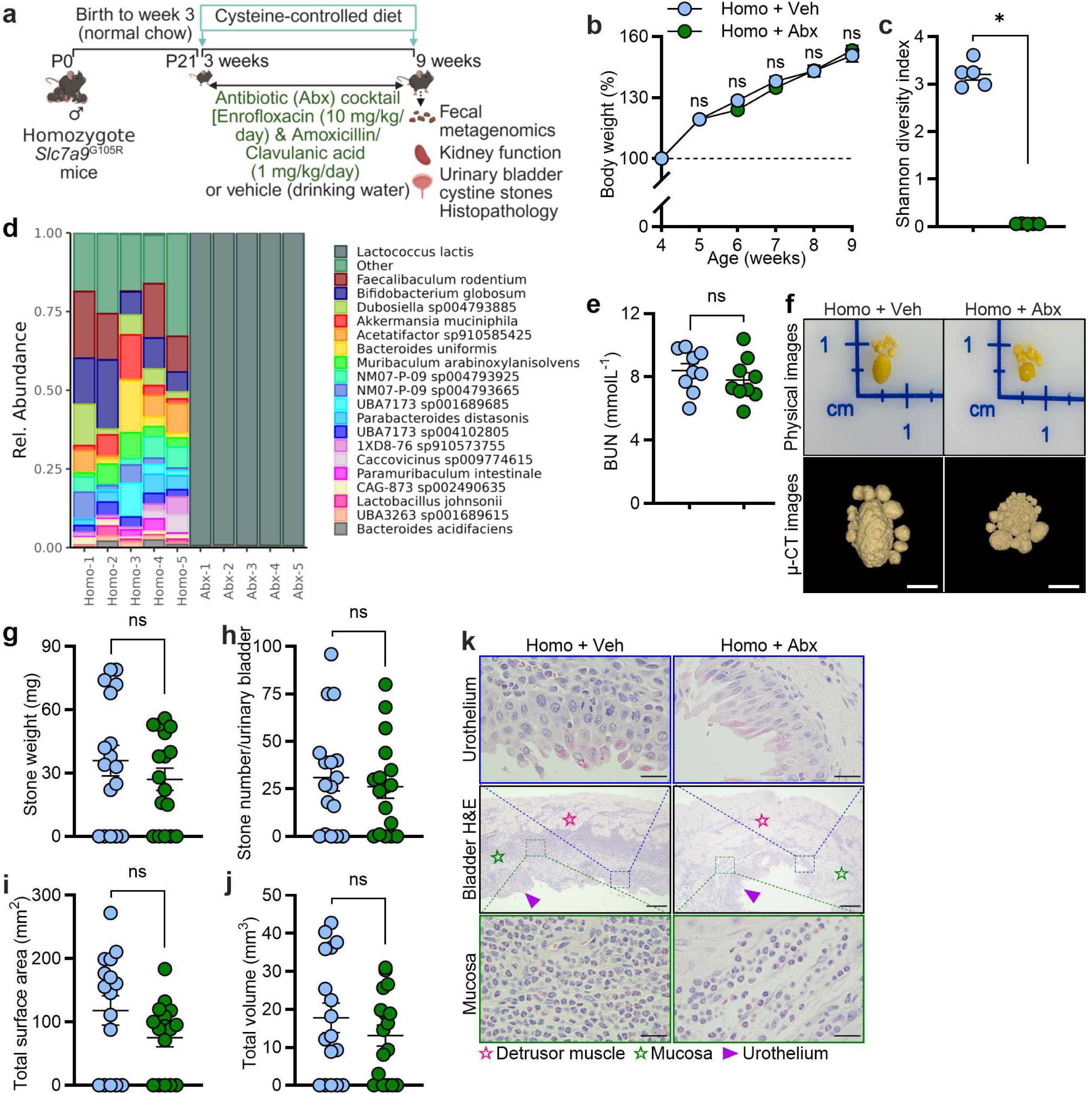
Prophylactic antibiotic treatment reduced fecal microbiome diversity and inflammation of the urinary bladder but did not affect kidney function or cystine stone growth. The impact of broad-spectrum antibiotic (Abx) administration on the fecal microbiome and features of cystinuria were assessed. (**a**) Schematic representation of Abx administration in male homozygote *Slc7a9*^G105R^ mice from weeks 3 to 9. (**b**) Percentage body weight change over time (*n* = 16-17 per group). (**c**) Shannon diversity index (*n* = 5 per group). (**d**) Bar plot showing the relative abundance of the top 20 species. (**e**) Serum blood urea nitrogen (BUN) levels (*n* = 10 per group). (**f**) Representative matched physical and µ-CT images of urinary cystine stones. Urinary bladder cystine stone (**g**) weight, (**h**) number, (**i**) total surface area, and (**j**) total stone volume (*n* = 16-17 per group). (**k**) Representative hematoxylin and eosin (H&E)-stained bladder sections. Ill, Ill, and ► denote the detrusor muscle, mucosa, and urothelium in the bladder section, respectively. The scale bar indicates 2.5 mm for the representative urinary bladder cystine stone µ-CT images, 200 µm for the bladder H&E section, and 25 µm for the magnified sections of urothelium and mucosa, respectively. Data was obtained from two independent experiments. Data are presented as the mean ± the standard error of the mean (SEM). Statistical differences were calculated using a two-way ANOVA with a Tukey multiple comparisons test (**b**) and a non-parametric unpaired Mann-Whitney t-test (**c**, **e**, **g**-**j**). *P* value < 0.05. * denotes a significant difference between the groups. ns indicates not significant.

Antibiotic treatment visually reduced mucosal inflammation in the urinary bladder compared to vehicle-treated controls (**Figure 5k**, **Supplementary Figure S17**). Antibiotic treatment also reduced inflammatory gene expression in the kidney, including *Cxcl1*, *Ccl2*, and *Il6* (**Supplementary Figure S18**, **Supplementary Table S5**-**6**). Furthermore, as mice are coprophagic, we assessed the impact of cohousing for 6 weeks on disease burden. Cohousing of wild-type and homozygote *Slc7a9*^G105R^ mice did not impact the severity of cystinuria or mucosal inflammation (**Supplementary Figure S19**).

### Prophylactic dietary supplementation with ALA improved kidney function, prevented urinary cystine stone growth, and reduced urinary bladder mucosal inflammation but caused urothelium disruption in homozygote *Slc7a9*^G105R^ mice

ALA has been shown to reduce cystine stone growth in *Slc3a1*-deficient mice,^20^ while other clinically used drugs for cystinuria, such as D-penicillamine and tiopronin, were ineffective.^13, 20, 38, 39^ ALA is now in a clinical trial for adult cystinuria (Trial # NCT02910531). We chose ALA as a positive control to ensure our new model was amenable to therapeutic intervention. Male homozygote *Slc7a9*^G105R^ mice were weaned onto an ALA-supplemented (**Supplementary Table S2**) or regular cysteine-controlled diet (control) at 3 weeks of age (**Figure 6a**). Homozygote *Slc7a9*^G105R^ mice fed an ALA-supplemented diet had significantly increased weight gain at 6, 7, 8, and 9 weeks compared to controls (**Figure 6b**). Homozygote *Slc7a9*^G105R^ mice fed an ALA-supplemented diet also had significantly reduced BUN levels (**Figure 6c**).

**Figure 6.**
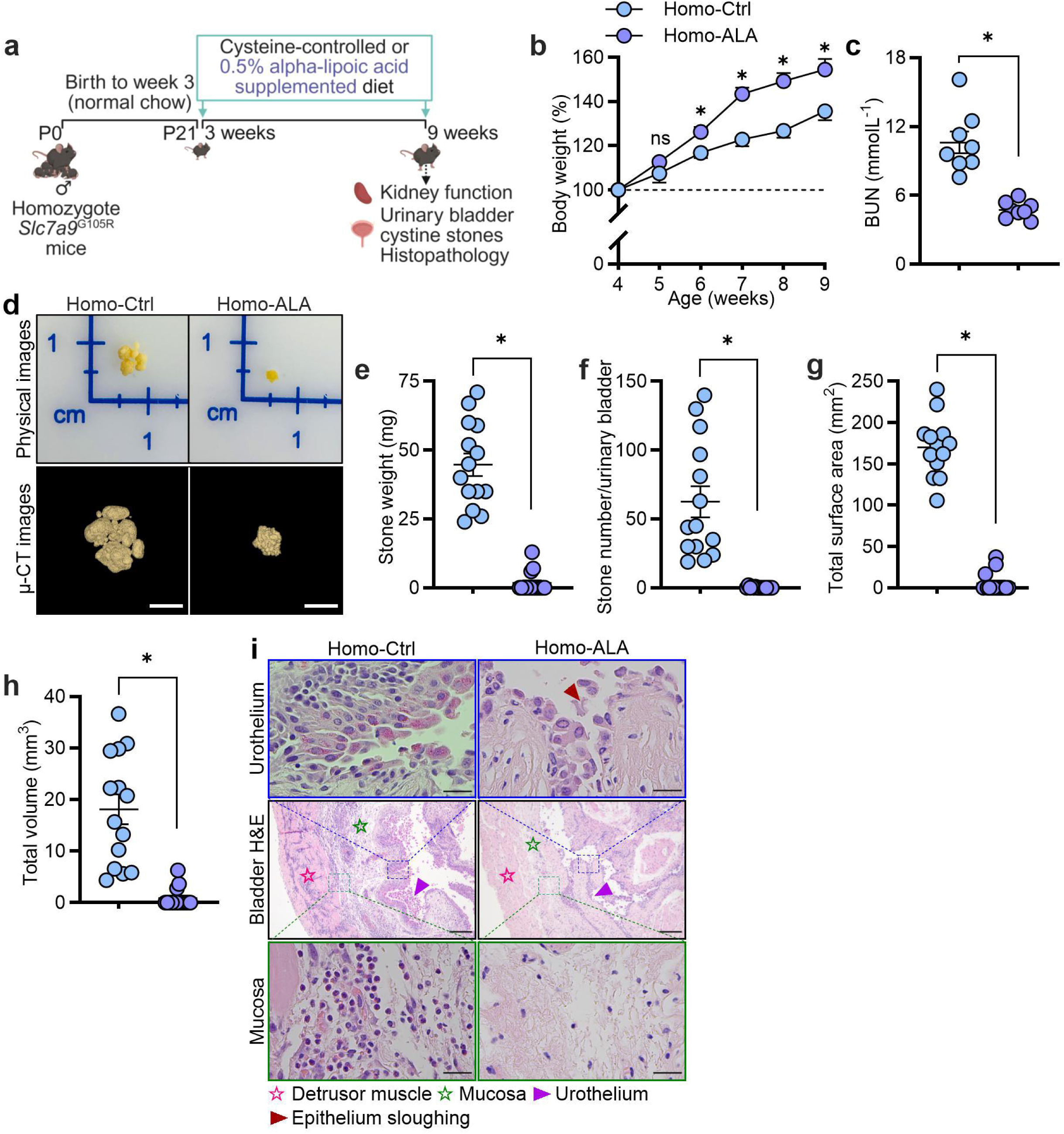
Prophylactic dietary supplementation with alpha-lipoic acid improved kidney function, prevented urinary cystine stone growth, and reduced urinary bladder mucosal inflammation but caused urothelium destruction in homozygote *Slc7a9*^G105R^ mice. The effect of prophylactic alpha-lipoic acid (ALA) dietary supplementation on kidney function, urinary cystine stone growth, and urinary bladder histopathology was assessed in male homozygote *Slc7a9*^G105R^ mice at 9 weeks. (**a**) Schematic representation of dietary ALA supplementation in homozygote *Slc7a9*^G105R^ mice. (**b**) Percentage body weight change over time (*n* = 14-15 per group). (**c**) Serum blood urea nitrogen (BUN) levels (*n* = 8 per group). (**d**) Matched representative physical and µ-CT images of urinary bladder cystine stones. Cystine stone (**e**) weight, (**f**) number, (**g**) total surface area, and (**h**) total volume (*n* = 14-15 per group). (**i**) Representative hematoxylin and eosin (H&E)-stained bladder sections. Ill, Ill, and ► denote the detrusor muscle, mucosa, and urothelium, respectively. The scale bar indicates 2.5 mm for µ-CT urinary bladder cystine stone image, 200 µm for the bladder H&E section, and 25 µm for the magnified sections of urothelium and mucosa, respectively. Data were obtained from two independent experiments. Data are presented as the mean ± the standard error of the mean (SEM). Statistical differences were calculated using a two-way ANOVA with a Tukey multiple comparison test (**b**) and non-parametric unpaired Mann-Whitney t-test (**c**, **e**-**h**). *P* value < 0.05. * denotes a statistical difference between the groups from weeks 6 to 9 (**b**). * also denotes a significant difference between the groups (**c**, **e**-**h**). ns indicates not significant.

Representative physical and µ-CT images showed urinary cystine stones were markedly reduced in homozygote *Slc7a9*^G105R^ mice fed an ALA-supplemented diet (**Figure 6d**). Homozygote *Slc7a9*^G105R^ mice fed an ALA-supplemented diet had significantly reduced cystine stone weight, number, surface area, and volume compared to controls (**Figure 6e**-**h**). BUN positively correlated with cystine stone weight, number, surface area, and volume (**Supplementary Figure S20a-d**).

Representative H&E-stained urinary bladder images showed that homozygote *Slc7a9*^G105R^ mice fed an ALA-supplemented diet had reduced mucosal inflammation but substantial urothelium destruction compared to controls (**Figure 6i**, **Supplementary Figure S21**). The ALA-supplemented diet also reduced the expression of inflammatory genes, including *Cxcl1*, *Cxcl2*, *Ccl2*, and *Il6*, in the kidneys of homozygote *Slc7a9*^G105R^ mice compared to controls (**Supplementary Figure S22**, **Supplementary Table S7**-**8**).

### Prophylactic dietary supplementation with ALA drastically changes serum and urinary metabolites in homozygote *Slc7a9*^G105R^ mice

The original work using ALA in *Slc3a1*^−/−^ mice suggested that unknown metabolites resulting from ALA treatment were likely responsible for the therapeutic effects of ALA.^20^ To explore this, untargeted serum and urine metabolomics were performed at week 9 (**Figure 7a**). ALA-supplemented diet significantly altered serum and urinary metabolites compared to homozygote controls (**Figure 7b**-**c**). The effect of ALA on urine metabolites was greater than on serum metabolites (**Supplementary Figure S23a**). ALA also had a more profound impact on serum and urinary metabolites than the cystinuria phenotype (**Supplementary Figure S23b**-**c**). Differentially abundant metabolites were mapped to metabolic pathways (**Figure 7d**, **Supplementary Figure S24a**-**b**) and metabolite interactions for both serum and urine (**Supplementary Figure S24c**).

**Figure 7.**
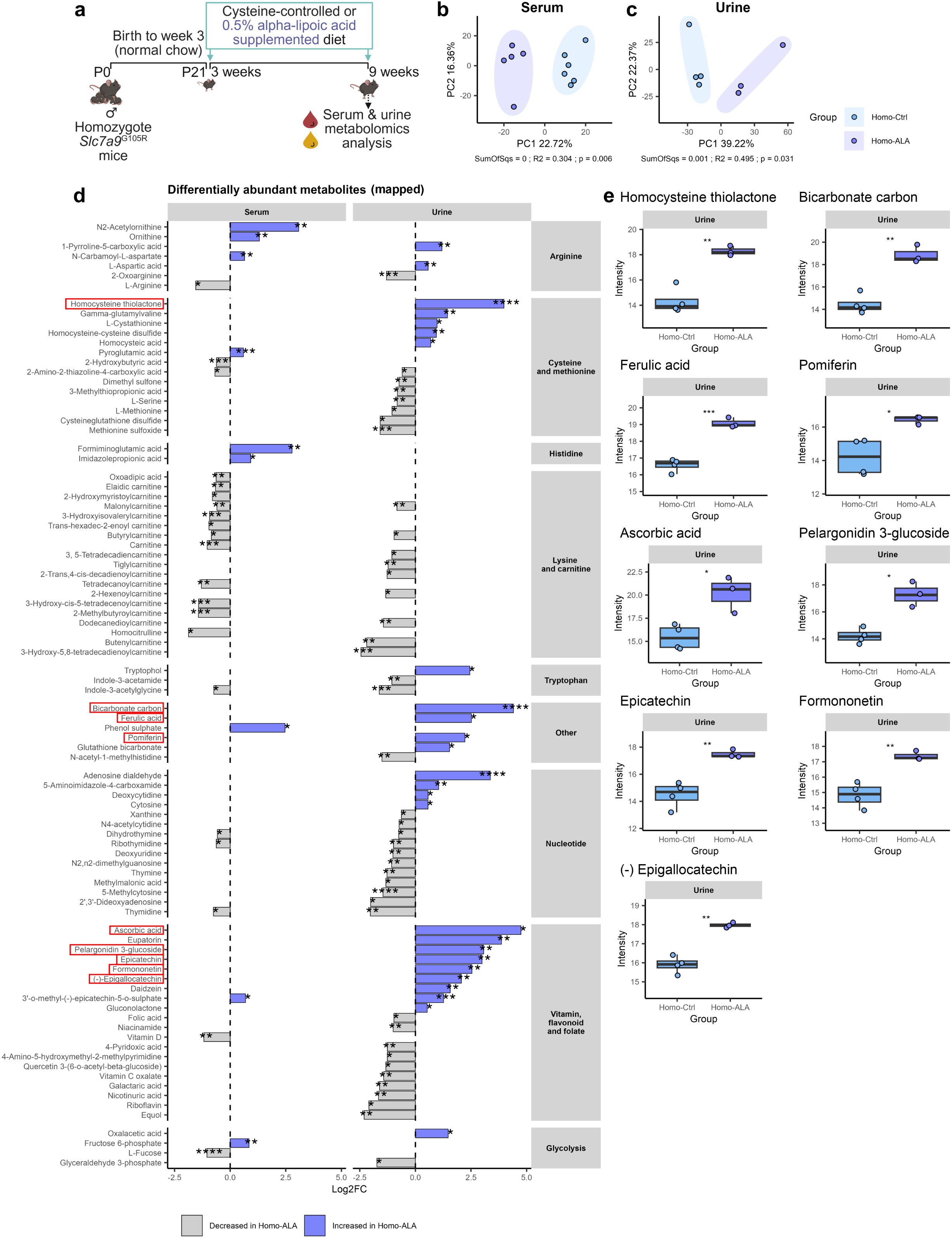
Prophylactic dietary supplementation with alpha-lipoic acid drastically changes serum and urinary metabolites in homozygote *Slc7a9*^G105R^ mice. Serum and urine were collected from male 9-week-old homozygote *Slc7a9*^G105R^ mice fed control (Homo-Ctrl) or ALA-supplemented diet (Homo-ALA) for untargeted metabolomics analysis. (**a**) Schematic representation of the experimental model. PCA plot for (**b**) serum and (**c**) PCA plot for urine metabolites. Ellipses represent the 95% confidence interval around the group centroid. PERMANOVA test results for the group. SumOfSqs (sum of squares): effect size; R2: variance explained, p: *P* value. (**d**) Differentially abundant serum (left) and urine (right) metabolites mapped to metabolic pathways (Log2FC > 0.5, FDR < 0.05). (**e**) Box plots showing select urinary metabolites linked to epithelial cell damage, cystine solubility, or antioxidant/anti-inflammatory effects. Serum samples: *n* = 5-6 per group. Urine samples: *n* = 3-4 per group. Data are presented as the median, interquartile range (IQR) (boxes) and 1.5x IQR (whiskers). Statistical significance between homozygote *Slc7a9*^G105R^ fed a control or ALA-supplemented diet was calculated using a Student’s t-test; *, **, ***, and **** denote *P* < 0.05, 0.01, 0.001, and 0.0001, respectively.

Urine metabolites increased by ALA were of particular interest, with several relating to epithelial cell damage (e.g., homocysteine thiolactone), cystine solubility (e.g., bicarbonate carbon, ascorbic acid) or antioxidant effects (e.g., ferulic acid, pomiferin, pelargonidin 3-glucoside, epicatechin, formononetin, (-) epigallocatechin) (**Figure 7e**). ALA also significantly decreased multiple lysine and carnitine metabolites in serum and urine (**Figure 7d**). Finally, ALA increased serum levels of the histidine metabolites formiminoglutamic acid and imidazolepropionic acid (**Figure 7d**).

### Untargeted serum and urine metabolite profiling identifies new biomarkers for cystinuria

To determine if serum and urinary metabolites could serve as new biomarkers for cystinuria, correlation analyses of metabolites with cystine stone number, surface area, and volume were performed and mapped to relevant metabolic pathways (**Figure 8a**). Those correlated with cystine stone number, surface area, and volume are shown as scatter plots (**Figure 8b**-**c**). Six serum metabolites correlated with cystine stone number, surface area, and volume (**Figure 8b**). Three were positively correlated (2-hydroxybutyric acid, 2-amino-2-thiazoline-4-carboxylic acid, and indole-3-acetylglycine), and three were negatively correlated (imidazolepropionic acid, ornithine, and formiminoglutamic acid). Similarly, six urine metabolites correlated with cystine stone number, surface area, and volume (**Figure 8c**). Three were positively correlated (L-serine, cysteineglutathione disulfide, and 2-hydroxybutyric acid), and three were negatively correlated (glutathione thiol, cysteic acid, and ascorbic acid). Additional serum and urine metabolites correlated with cystine stone number, surface area, and volume but did not map to relevant metabolic pathways (**Supplementary Figure S25a**-**b**).

**Figure 8.**
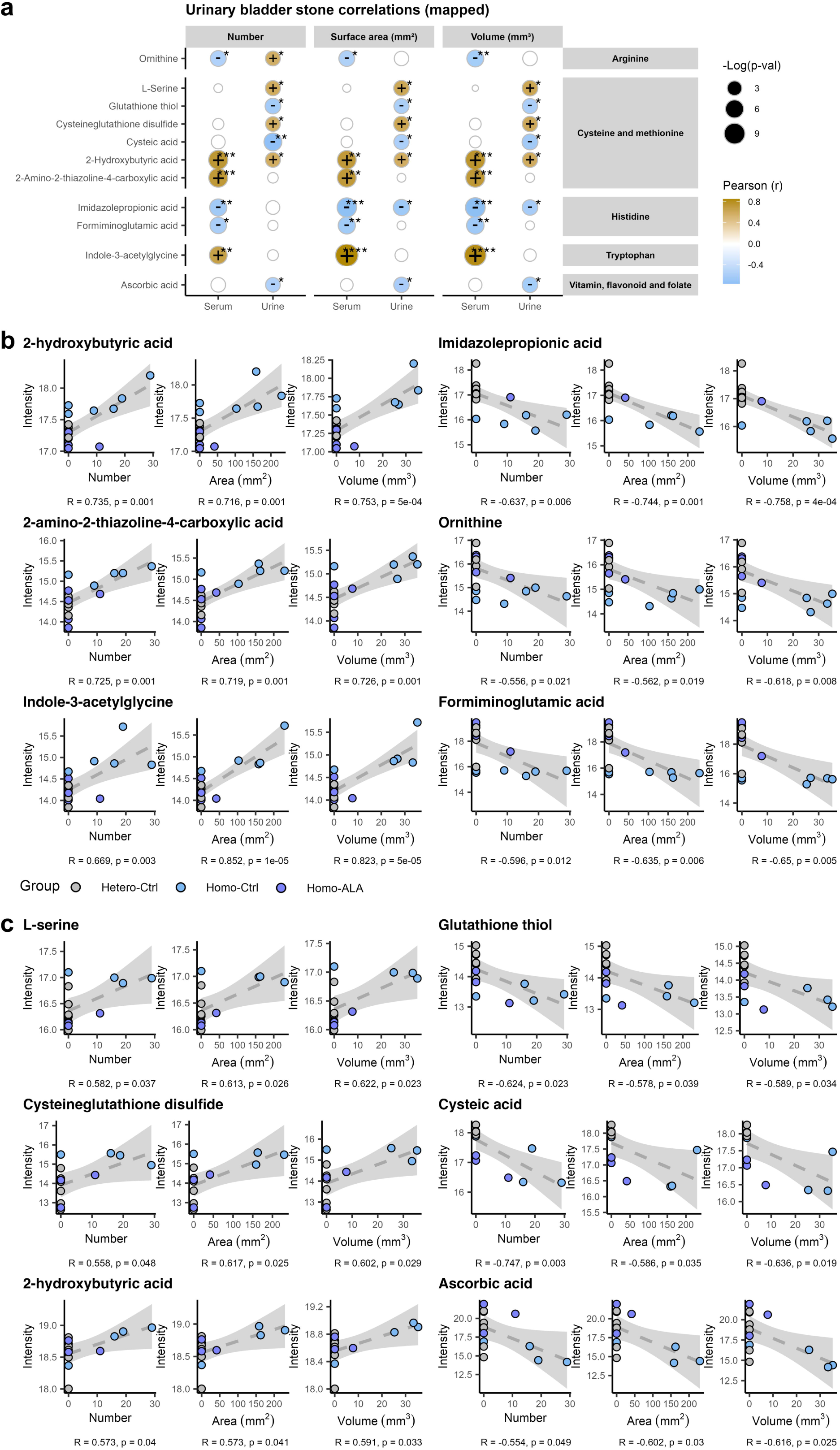
Untargeted serum and urine metabolite profiling identifies new biomarkers for cystinuria. Serum and urine were collected from male 9-week-old heterozygote (Hetero-Ctrl), and homozygote *Slc7a9*^G105R^ mice fed control (Homo-Ctrl) or ALA-supplemented diet (Homo-ALA) for untargeted metabolomics analysis. (**a**) Summary of urinary bladder cystine stone number, surface area, and volume correlated with serum and urinary metabolites that map to metabolic pathways using Pearson correlation. Empty circles with a grey outline in the plot indicate metabolites for which the Pearson correlation was not significant. Empty spots in the plot indicate that a metabolite was not detected in the serum or urine. (**b**) Serum metabolites that correlate with cystine stone number, surface area, and volume. (**c**) Urine metabolites that correlate with cystine stone number, surface area, and volume. Data are presented as the Pearson correlation (R) coefficient, and *, **, ***, and **** denote *P* values of 0.05, 0.01, 0.001, and 0.0001, respectively. Grey shading represents the 95% confidence interval.

### Therapeutic dietary supplementation with ALA reduced cystine stone burden and urinary bladder mucosal inflammation, but caused damage to the urothelium at 12 weeks

Given that ALA effectively prevented cystine stone growth in homozygote *Slc7a9*^G105R^ mice, we sought to determine if ALA can effectively treat established disease. For this, a subset of mice were switched to an ALA-supplemented cysteine-controlled diet at 9 weeks, while controls were kept on the cysteine-controlled diet until week 12 (**Figure 9a**).

**Figure 9.**
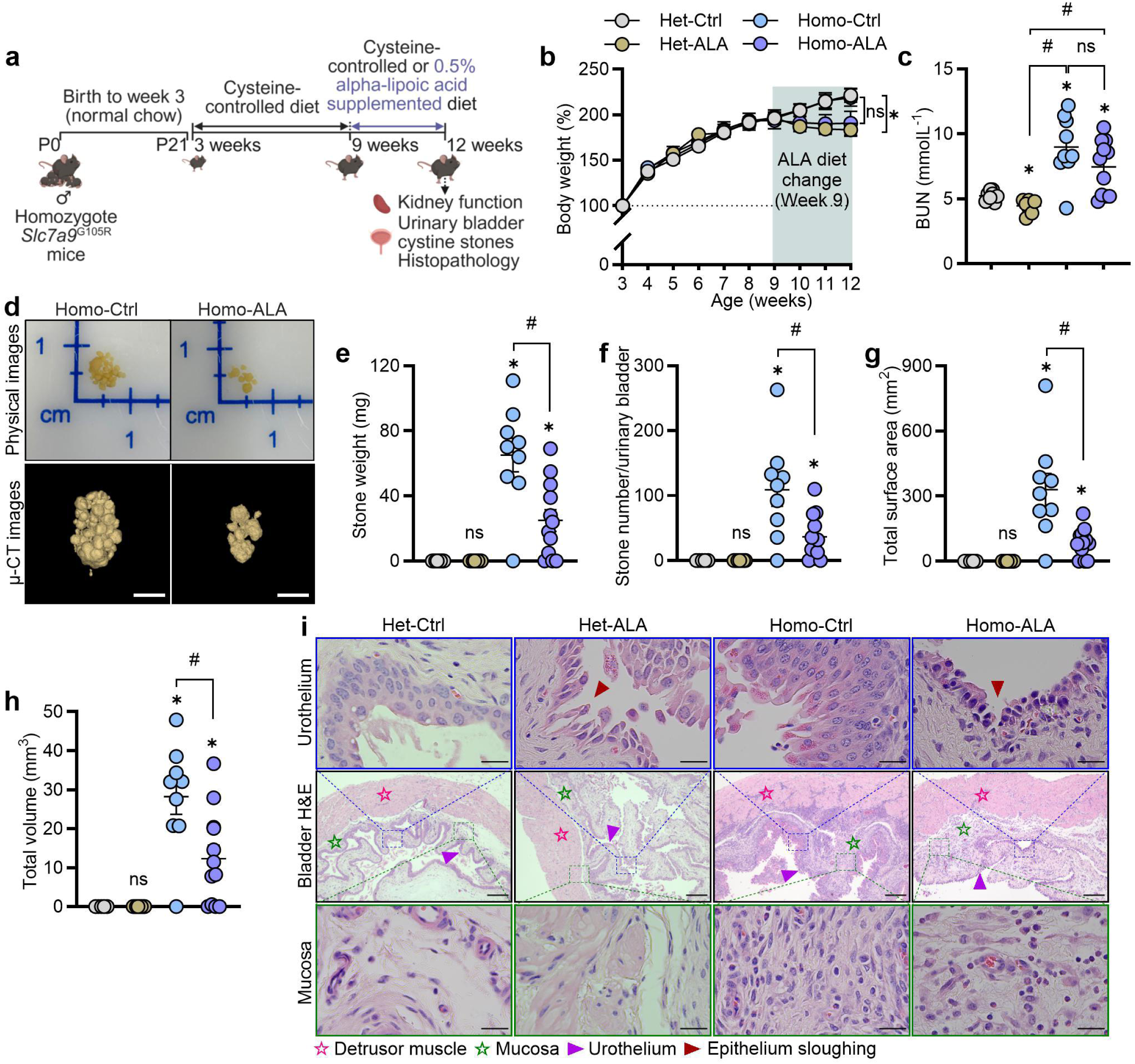
Therapeutic dietary supplementation with alpha-lipoic acid reduced cystine stone growth and urinary bladder mucosal inflammation in homozygote *Slc7a9*^G105R^ mice at 12 weeks. The therapeutic effect of alpha-lipoic acid (ALA) dietary supplementation on kidney function, urinary cystine stone growth, and urinary bladder histopathology was assessed in male homozygote *Slc7a9*^G105R^ mice at 12 weeks after 3 weeks of treatment. (**a**) Schematic representation of ALA supplementation in homozygote *Slc7a9*^G105R^ mice. (**b**) Percentage body weight change over time (*n* = 8-12 per group). (**c**) Serum blood urea nitrogen (BUN) levels (*n* = 7-11 per group). (**d**) Matched representative physical and µ-CT images of urinary cystine stones. Urinary cystine stone (**e**) weight, (**f**) number, (**g**) total surface area, and (**h**) total stone volume (heterozygote controls *n* = 5 and homozygotes *n* = 9-12 per group). (**i**) Representative hematoxylin and eosin (H&E)-stained bladder sections. Ill, Ill, and ► denote the detrusor muscle, mucosa, and urothelium, respectively. The scale bar indicates 2.5 mm for urinary cystine stones, 200 µm for the bladder H&E section, and 25 µm for the magnified sections of urothelium and mucosa. Data were obtained from two independent experiments. Data are presented as the mean ± the standard error of the mean (SEM). Statistical differences were calculated using a two-way ANOVA with a Tukey multiple comparison test (**b**) and non-parametric unpaired Mann-Whitney t-test (**c**, **e**-**h**). *P* value < 0.05. * denotes a statistical difference between heterozygote controls and heterozygote ALA-supplemented group at week 12 (**b**). * also denotes a significant difference compared to heterozygote controls (**c**, **e**-**h**). ^#^ denotes statistical differences between two consecutive groups. ns indicates not significant.

Heterozygote *Slc7a9*^G105R^ (Het-ALA) fed an ALA-supplemented diet had significantly decreased body weight compared to the heterozygote controls (Het-Ctrl), whilst homozygote *Slc7a9*^G105R^ (Homo-ALA) fed an ALA-supplemented diet were comparable to the homozygote controls (Homo-Ctrl) (**Figure 9b**). Heterozygote *Slc7a9*^G105R^ mice fed an ALA-supplemented diet had significantly reduced BUN compared to the heterozygote controls (**Figure 9c**). Homozygote *Slc7a9*^G105R^ mice fed an ALA-supplemented diet had similar BUN levels to homozygote *Slc7a9*^G105R^ controls (**Figure 9c**). Both homozygote controls and homozygotes fed an ALA-supplemented diet had increased BUN compared to their respective heterozygote controls (**Figure 9c**).

Representative physical and µ-CT images of urinary bladder cystine stones show that homozygote *Slc7a9*^G105R^ mice fed an ALA-supplemented diet had visibly reduced stones compared to the homozygote controls (**Figure 9d**). Homozygote *Slc7a9*^G105R^ mice fed an ALA-supplemented diet had significantly decreased cystine stone weight, number, surface area, and volume compared to the homozygote controls (**Figure 9e**-**h**).

Representative H&E-stained urinary bladder sections showed a marked reduction in mucosal inflammation in homozygote *Slc7a9*^G105R^ mice fed an ALA-supplemented diet compared to homozygote controls (**Figure 9i**, **Supplementary Figure S26**). Notably, there were visually striking urothelial changes resembling urothelium barrier dysfunction in heterozygote *Slc7a9*^G105R^ mice fed an ALA-supplemented diet compared to heterozygote controls (**Figure 9i**). Consistent with our week 9 data, homozygote *Slc7a9*^G105R^ controls displayed substantial hyperplasia of the urothelium compared to heterozygote controls (**Figure 9i**). Homozygote *Slc7a9*^G105R^ mice fed an ALA-supplemented diet exhibited visibly reduced urothelial thickness but showed signs of urothelial barrier disruption compared to heterozygote controls (**Figure 9i**).

## Discussion

Cystinuria remains a challenging genetic disorder with limited treatment options and a high burden of recurrent stone formation. Although its genetic underpinnings are well characterized, progress in developing targeted therapies has been constrained by the lack of preclinical models that accurately reflect human pathogenic mutations. In this study, we developed and characterized a *Slc7a9*^G105R^ knock-in mouse model that recapitulates the most common disease-causing variant in *SLC7A9* in human cystinuria. This model exhibits a robust and early-onset phenotype characterized by elevated urinary cystine and dibasic amino acids, substantial cystine stone formation, and alterations in metabolites and the fecal microbiota. This new model offers a genetically precise platform for investigating disease mechanisms, evaluating potential therapeutic strategies, and identifying candidate biomarkers that warrant further validation in human cohorts.

In contrast to existing mouse models of cystinuria, which often exhibit delayed disease onset and low penetrance of stone formation,^4, 16–18, 23, 40^ the *Slc7a9*^G105R^ knock-in mice developed early and highly penetrant cystine stone formation, with over 90% of homozygous animals (both male and female) exhibiting substantial stone burden by just 9 weeks of age, equivalent to late teens or early adulthood in humans. This uniformity across sexes contrasts with previous clinical and preclinical reports indicating a higher stone burden in males.^4, 10, 17, 23, 40^ Epidemiological studies have indicated a male predominance in cystine stone formation beginning in early childhood, before puberty, which suggests a limited role for sex hormones in disease onset.^10^ Consistent with this, stone formation in *Slc7a9*^G105R^ mice was detected as early as 3 weeks of age, underscoring the model’s relevance for investigating pediatric and adolescent cystinuria. Additionally, we found that pharmacological suppression of testosterone using spironolactone in male *Slc7a9*^G105R^ mice reduced the number of stones but not the total stone volume. This finding parallels the phenotype observed in female *Slc7a9*^G105R^ mice, who typically develop fewer but larger stones. These data suggest that while sex hormones may modulate certain aspects of stone morphology and formation dynamics, they are unlikely to be the primary determinants of disease severity. Recent mechanistic insights have further illuminated the molecular underpinnings of sex differences in cystinuria. Notably, male mice exhibit increased expression of mitochondrial *Slc3a1*, which enhances cystine uptake and promotes stone formation. Conversely, females display lower mitochondrial *Slc3a1* expression, correlating with decreased cystine accumulation and relative protection from severe stone burden.^15^ This discovery highlights intrinsic sex-specific differences in transporter expression and mitochondrial function that operate independently of circulating sex hormones, thereby refining our understanding of cystinuria pathogenesis, at least when driven by *SLC3A1*.^15^ Together, these findings underscore the need to consider both hormonal and cell-intrinsic factors in developing therapeutic strategies to mitigate cystine stone disease, which is likely to be genotype-specific.

Consistent with previous cystinuria mouse models,^15, 17, 19, 20^ cystine stone formation in *Slc7a9*^G105R^ mice occurred predominantly in the urinary bladder, with no detectable renal calculi. This pattern contrasts with human cystinuria, where cystine stones most commonly form in the kidneys.^3^ The basis for this species-specific difference likely reflects anatomical and physiological distinctions, including variations in urinary tract structure, flow dynamics, and crystal aggregation properties between mice and humans. Although renal stone deposition was absent, the continuous passage of cystine crystals through the upper urinary tract may contribute to renal tubular stress or injury.

Increased BUN has been reported in existing cystinuria mouse models, indicating renal impairment associated with stone disease.^15, 16^ However, we are the first to longitudinally profile BUN and demonstrate a strong positive correlation between BUN levels and cystine stone burden. In our model, BUN progressively increased with stone growth, suggesting that ongoing lithogenesis may contribute to renal dysfunction or subclinical tubular stress, even in the absence of overt renal failure. This observation aligns with prior evidence that the deposition of calcium oxalate or cystine crystals induces tubular injury and oxidative stress, thereby disrupting urea handling and causing early changes in renal biomarkers.^16, 41–43^ Indeed, increased amino acid loss can lead to compensatory protein catabolism, particularly when the body is attempting to maintain amino acid pools. Increased urea cycle activity, particularly from arginine and ornithine metabolism, can lead to increased urea (and BUN) production. This may explain why we observe elevated BUN levels even when the GFR remains stable. Importantly, prophylactic ALA treatment restored BUN to baseline levels, supporting its potential as a marker of disease burden and therapeutic response. While BUN is a routine clinical biomarker of renal function with limited specificity, its strong association with stone burden in *Slc7a9*^G105R^ mice, combined with its widespread clinical use, highlights its promise as a simple, non-invasive biomarker for monitoring cystinuria progression and treatment efficacy.

*Slc7a9*^G105R^ mice primarily exhibited bladder pathology, including urothelial disruption and mucosal inflammation, with only mild and infrequent renal tubular injury observed histologically at 9 weeks. This contrasts with other cystinuria models, such as *Slc3a1*^−/−^ and *Slc7a9*^−/−^ mice, which display progressive renal pathology over time, including interstitial fibrosis and tubular atrophy.^16, 18^ Notably, disease onset in *Slc7a9*^G105R^ mice occurs early, with a substantial cystine stone burden present in over 90% of mice by 9 weeks, allowing for the assessment of active lithogenesis and tissue responses without the need for prolonged aging. Despite limited histological evidence of kidney injury, we observed increased expression of proinflammatory genes in the kidney, including *Cxcl2*, *Ccl2*, and *Il6*. These chemokines are well-established mediators of sterile inflammation and are implicated in renal injury induced by calcium oxalate crystals.^44–46^ Similar patterns have been reported in models of calcium oxalate nephropathy and human stone disease, suggesting that innate immune responses may be a conserved feature of crystal-associated kidney injury.^41, 47^ Interestingly, the expression of these inflammatory genes was reduced by both antibiotic and ALA treatment, suggesting potential contributions from gut microbial factors and oxidative stress. While inflammation in this model is subtle and not associated with significant histopathological renal changes at 9 weeks, our findings suggest that cystine lithogenesis may trigger early immune activation in the kidney. These responses could reflect a broader renal stress signature and may offer a mechanistic link between stone burden and subclinical renal dysfunction.

Consistent with previous reports, homozygous *Slc7a9*^G105R^ mice exhibited significantly increased urinary levels of cystine, ornithine, lysine, and arginine.^10, 16, 18, 23, 48^ The presence of classic hexagonal cystine crystals in the urine of these mice further confirms the elevated urinary cystine concentrations and mirrors a hallmark diagnostic feature observed in human patients.^15, 18, 20, 23^ These findings validate the biochemical fidelity of the *Slc7a9*^G105R^ model and support its utility for studying cystine handling and crystal formation in a genetically accurate context.

Metabolomics has emerged as a powerful approach for uncovering disease mechanisms, including those driving kidney stone formation, by revealing perturbations in key metabolic pathways such as amino acid metabolism, fatty acid oxidation, and redox homeostasis.^24^ Previous work using *Slc7a9^−/−^* mice identified the urinary ratio of S-methyl-L-ergothioneine to L-ergothioneine as a potential biomarker for cystinuria.^49^ In contrast, our untargeted metabolomics analysis of *Slc7a9*^G105R^ mice detected L-ergothioneine in both serum and urine but not its methylated derivative, and levels of L-ergothioneine were not significantly altered in mice with cystinuria (FDR > 0.05). These findings suggest that the *Slc7a9*^G105R^ model may exhibit distinct metabolic signatures compared to knockout models, highlighting the importance of mutation-specific approaches in biomarker discovery.

In our current study, serum 2-hydroxybutyric acid emerged as a strong positive correlate of cystine stone burden in the *Slc7a9*^G105R^ model, highlighting its potential as a non-invasive biomarker for disease progression. Biochemically, 2-hydroxybutyric acid is generated as a byproduct during the conversion of cystathionine to cysteine in the transsulfuration pathway—a process up-regulated during increased glutathione synthesis. This links 2-hydroxybutyric acid directly to sulfur amino acid metabolism and redox homeostasis.^50^ Elevated circulating 2-hydroxybutyric acid has been documented in various pathological states characterized by oxidative stress, mitochondrial dysfunction, and impaired glucose metabolism,^51–53^ conditions that may plausibly intersect with the metabolic and inflammatory disturbances observed in cystinuria. Although not previously associated with this disease, the biochemical proximity of 2-hydroxybutyric acid to cysteine metabolism and glutathione turnover provides a mechanistic rationale for its elevation in cystinuria. These findings position 2-hydroxybutyric acid as a candidate biomarker for monitoring disease activity and treatment responses.

In our untargeted serum metabolomics analysis, we also identified a significant increase in serum 2-amino-2-thiazoline-4-carboxylic acid, a stable metabolite formed by the reaction of cysteine with cyanide. Notably, serum 2-amino-2-thiazoline-4-carboxylic acid levels correlated positively with cystine stone burden, suggesting a potential link between altered cysteine metabolism and disease severity. Beyond its role in stone formation, cysteine also participates in essential metabolic processes, including antioxidant defense (as a precursor of glutathione) and detoxification.^54, 55^ Elevated 2-amino-2-thiazoline-4-carboxylic acid may therefore reflect a diversion of free cysteine toward detoxification pathways, potentially driven by inflammation, oxidative stress, or excess cystine accumulation in the kidney. While 2-amino-2-thiazoline-4-carboxylic acid is best known as a biomarker of cyanide exposure,^56,57^ our findings suggest it may also serve as an indirect marker of metabolic adaptations to impaired cystine reabsorption and stone formation. These results highlight a previously unrecognized metabolic consequence of cystinuria and raise the possibility that 2-amino-2-thiazoline-4-carboxylic acid could be developed as a non-invasive biomarker of disease burden.

Interestingly, we observed elevated levels of indole-3-acetylglycine in serum, which positively correlated with cystine stone burden. Indole-3-acetylglycine is a conjugated metabolite derived from indole-3-acetic acid, itself a product of microbial tryptophan metabolism. Although indole-3-acetylglycine has not previously been associated with cystinuria, its increase may reflect broader alterations in host–microbiome metabolic interactions secondary to tubulointerstitial injury or subclinical renal dysfunction. Indole derivatives such as indoxyl sulfate and indole-3-acetic acid are increasingly recognized as uremic solutes that accumulate in kidney disease and contribute to oxidative stress, inflammation, and fibrosis.^58–60^ Moreover, gut microbial metabolism of tryptophan influences urinary composition, including pH and citrate levels, which are essential modifiers of cystine stone risk.^61, 62^ Thus, elevated indole-3-acetylglycine may represent a biomarker of microbial metabolic perturbation or impaired renal clearance in cystinuria.

The gut microbiota is increasingly implicated in kidney stone disease, particularly through observational and experimental studies in calcium oxalate nephrolithiasis.^25, 63, 64^ In this context, dysbiosis, characterized by the loss of oxalate-degrading and butyrate-producing bacteria, has been associated with increased stone risk and inflammation.^27, 65^ In *Slc7a9*^G105R^ mice, fecal metagenomic analysis revealed microbial shifts marked by reduced abundance of *Lactococcus cremoris* and *Bacteroides caecimuris*, alongside increased abundance of *Acutalibacter muris* and *NM07-P-09 sp004793925*. The loss of *L. cremoris*, a species shown to reduce renal inflammation in hyperuricemia models via increased short-chain fatty acid production and improved host metabolism^66^, may contribute to a pro-inflammatory or metabolically unfavorable state in cystinuria. Similarly, a decrease in *B. caecimuris*, a commensal organism associated with metabolic regulation and epithelial interaction^67, 68^ may reflect a shift away from microbial profiles supportive of homeostasis. In contrast, enrichment of *A. muris* and *Leptogranulimonas caecicola* (formerly *NM07-P-09 sp004793925*), while not functionally characterized, raises the possibility of involvement in microbial succession, a process whereby changes in environmental or host conditions drive temporal shifts in microbial community composition and function.^25^

Interestingly, antibiotic-mediated microbiota depletion did not reduce stone burden but led to significant improvement in bladder histopathology, suggesting that the microbiome may influence local inflammatory tone rather than directly modulating cystine stone formation. Functional pathway analysis of fecal metagenomes revealed upregulation of bacterial L-cysteine biosynthesis pathways in homozygous *Slc7a9*^G105R^ mice. This may reflect a microbiota-level compensatory response to urinary cystine loss or an adaptive mechanism to buffer oxidative stress through enhanced glutathione precursor availability. Conversely, down-regulation of microbial methylglyoxal detoxification pathways may impair redox regulation and contribute to the accumulation of reactive dicarbonyls with potential renal toxicity. Together, these findings support a complex interplay between microbiome composition, microbial metabolism, and inflammatory tone in cystinuria, providing a foundation for future studies investigating microbe–host–metabolite interactions and their therapeutic relevance.

Current treatment options for cystinuria primarily rely on urinary alkalinization and thiol-based agents, such as tiopronin and D-penicillamine; however, their use is frequently limited by gastrointestinal intolerance, dermatologic reactions, and poor long-term adherence.^38, 39, 69^ ALA, a mitochondrial antioxidant with thiol-scavenging and metal-chelating properties, has previously been shown to reduce cystine stone formation in *Slc3a1* knockout mice.^20^ However, subsequent studies noted that synthetic ALA itself does not directly alter cystine solubility, suggesting that its protective effect may be mediated by downstream metabolites rather than the parent compound.^20^ Notably, ALA has not previously been tested in *Slc7a9*-deficient models, which represent a substantial proportion of patients with cystinuria. In the current study, we evaluated ALA for the first time in *Slc7a9*^G105R^ mice. ALA significantly attenuated stone growth in both preventive and therapeutic settings, accompanied by shifts in systemic and urinary metabolite profiles. Among the metabolites elevated in ALA-treated mice were bicarbonate carbon, and ascorbic acid, both of which have been independently associated with increased cystine solubility and reduced stone formation.^70, 71^ A phase II clinical trial is currently underway to evaluate the efficacy of daily oral ALA in reducing cystine stone recurrence, further underscoring the relevance of these findings to translational efforts.^29^

However, our data in mice also raises important safety considerations. Despite its efficacy in reducing stone burden, ALA treatment was associated with histological evidence suggestive of urothelial barrier disruption. These findings suggest that while ALA may confer metabolic and lithogenic benefits, it could also compromise epithelial homeostasis in the urinary tract. Given that the urothelium plays a critical role in maintaining barrier function and modulating local immune responses, potential epithelial toxicity should be carefully monitored as ALA advances through clinical trials. Future studies should aim to delineate the dose-response relationship, the reversibility of epithelial changes, and whether co-therapies can mitigate these effects. Overall, while ALA represents a promising therapeutic candidate, its dual impact on stone formation and urothelial integrity underscores the need for a balanced risk–benefit assessment in translational and clinical contexts.

In conclusion, the generation of a novel *Slc7a9*^G105R^ mouse model provides a robust and translational platform for studying cystinuria driven by a clinically relevant pathogenic point mutation. This model exhibits very early-onset cystinuria, faithfully recapitulating the genetic and metabolic hallmarks of the human condition, while also unveiling distinct metabolomic signatures that may serve as novel biomarkers. Our findings emphasize the intricate interplay among genetic, metabolic, and microbial factors in cystinuria and highlight the therapeutic promise of ALA in modulating these pathways to reduce stone formation and inflammation. Ultimately, this model lays a critical foundation for evaluating new therapeutic approaches, including precision medicine strategies, such as targeted gene therapy, that can propel future translational research in cystinuria. Collectively, the *Slc7a9*^G105R^ is well-positioned to serve as the new gold standard model for preclinical and translational research in cystinuria.

## Data statement

All research data supporting the findings of this study are included in the main figures and the online supplementary material. The raw metagenomics data are available at NCBI SRA (Accession number: PRJNA1131664).

## Acknowledgements

The authors would like to acknowledge the Australian Phenomics Facility, Alfred Research Alliance-Monash Biomedical Imaging (ARA-MBI), Monash Genomics and Bioinformatics Platform, Monash Proteomics and Metabolomics Platform, and Monash Micro Imaging for their contributions. Schematics were created using Biorender (License # Bhatt, N., 2025; https://BioRender.com/jbaiy2p, <u>/izu2u4i</u>, <u>/rzta4m8</u>, <u>/ran1nie</u>, <u>/uo3dhz8</u>, <u>/ihl502m</u>, <u>/33x9bv9</u>, /tdjtm2n).

## Author contributions

Conceptualization: NPB, AVD, GB, SHJ, and MRS. Experiments: NPB, GRR, and TTHN. Data analysis: NPB, GRR, GI, TTHN, CRBA, AP, CKB and MRS. Result interpretation: NPB, GRR, GI, TTHN, AP, CKB, BJM, and MRS. Manuscript writing and editing: NPB, GI, AP, CKB, and MRS. All authors read and approved the final manuscript.

Supplementary material is available online at www.kidney-international.org.

## References

1. Calonge MJ, Gasparini P, Chillarón J, et al. Cystinuria caused by mutations in rBAT, a gene involved in the transport of cystine. Nat Genet. 1994;6(4):420–5.

2. Feliubadaló L, Font M, Purroy J, et al. Non-type I cystinuria caused by mutations in SLC7A9, encoding a subunit (bo,+AT) of rBAT. Nat Genet. 1999;23(1):52–7.

3. Servais A, Thomas K, Dello Strologo L, et al. Cystinuria: clinical practice recommendation. Kidney International. 2021;99(1):48–58.

4. Ercolani M, Sahota A, Schuler C, et al. Bladder outlet obstruction in male cystinuria mice. Int Urol Nephrol. 2010;42(1):57–63.

5. Evan AP, Coe FL, Lingeman JE, et al. Renal crystal deposits and histopathology in patients with cystine stones. Kidney International. 2006;69(12):2227–2235.

6. Seyedzadeh A, Momtaz HE, Moradi MR, et al. Pediatric cystine calculi in west of Iran: a study of 22 cases. Urol J. 2006;3(3):134–7.

7. Gambaro G, Favaro S, D’Angelo A. Risk for renal failure in nephrolithiasis. Am J Kidney Dis. 2001;37(2):233–43.

8. Worcester EM, Coe FL, Evan AP, et al. Reduced renal function and benefits of treatment in cystinuria vs other forms of nephrolithiasis. BJU Int. 2006;97(6):1285–90.

9. Lindell A, Denneberg T, Granerus G. Studies on renal function in patients with cystinuria. Nephron. 1997;77(1):76–85.

10. Dello Strologo L, Pras E, Pontesilli C, et al. Comparison between SLC3A1 and SLC7A9 cystinuria patients and carriers: a need for a new classification. J Am Soc Nephrol. 2002;13(10):2547–53.

11. Chillarón J, Font-Llitjós M, Fort J, et al. Pathophysiology and treatment of cystinuria. Nat Rev Nephrol. 2010;6(7):424–34.

12. Saravakos P, Kokkinou V, Giannatos E. Cystinuria: Current Diagnosis and Management. Urology. 2014;83(4):693–699.

13. Bhatt NP, Deshpande AV, Starkey MR. Pharmacological interventions for the management of cystinuria: a systematic review. J Nephrol. 2024;37(2):293–308.

14. Warren H, Poon D, Srinivasan R, et al. Non-contrast computed tomography characteristics in a large cohort of cystinuria patients. World J Urol. 2021;39(7):2753–2757.

15. Su J, Pan Y, Zhong F, et al. Mitochondrial SLC3A1 regulates sexual dimorphism in cystinuria. Genes Dis. 2025;12(3):101472.

16. Woodard LE, Welch RC, Veach RA, et al. Metabolic consequences of cystinuria. BMC Nephrology. 2019;20(1):227.

17. Sasaki H, Sasaki T, Hiura K, et al. A mouse model of type B cystinuria due to spontaneous mutation in FVB/NJcl mice. Urolithiasis. 2022;50(6):679–684.

18. Feliubadaló L, Arbonés ML, Mañas S, et al. *Slc7a9*-deficient mice develop cystinuria non-I and cystine urolithiasis. Hum Mol Genet. 2003;12(17):2097–108.

19. Font-Llitjós M, Feliubadaló L, Espino M, et al. *Slc7a9* knockout mouse is a good cystinuria model for antilithiasic pharmacological studies. Am J Physiol Renal Physiol. 2007;293(3):F732–40.

20. Zee T, Bose N, Zee J, et al. α-Lipoic acid treatment prevents cystine urolithiasis in a mouse model of cystinuria. Nat Med. 2017;23(3):288–290.

21. Di Giacopo A, Rubio-Aliaga I, Cantone A, et al. Differential cystine and dibasic amino acid handling after loss of function of the amino acid transporter b0,+AT (*Slc7a9*) in mice. Am J Physiol Renal Physiol. 2013;305(12):F1645–55.

22. Kum F, Wong K, Game D, et al. Hypertension and renal impairment in patients with cystinuria: findings from a specialist cystinuria centre. Urolithiasis. 2019;47(4):357–363.

23. Peters T, Thaete C, Wolf S, et al. A mouse model for cystinuria type I. Human Molecular Genetics. 2003;12(17):2109–2120.

24. Xiong Y, Song Q, Zhao S, et al. Serum metabolomics study reveals a distinct metabolic diagnostic model for renal calculi. Heliyon. 2024;10(11):e32482.

25. Jones-Freeman B, Chonwerawong M, Marcelino VR, et al. The microbiome and host mucosal interactions in urinary tract diseases. Mucosal Immunol. 2021;14(4):779–792.

26. Hunthai S, Usawachintachit M, Taweevisit M, et al. Unraveling the role of gut microbiota by fecal microbiota transplantation in rat model of kidney stone disease. Sci Rep. 2024;14(1):21924.

27. Denburg MR, Koepsell K, Lee JJ, et al. Perturbations of the Gut Microbiome and Metabolome in Children with Calcium Oxalate Kidney Stone Disease. J Am Soc Nephrol. 2020;31(6):1358–1369.

28. Cil O, Perwad F. α-Lipoic Acid (ALA) Improves Cystine Solubility in Cystinuria: Report of 2 Cases. Pediatrics. 2020;145(5).

29. Chi T. Lipoic Acid Supplement for Cystine Stone (ALA). ClinicalTrialsgov. 2023;ID NCT02910531

30. Blanco-Míguez A, Beghini F, Cumbo F, et al. Extending and improving metagenomic taxonomic profiling with uncharacterized species using MetaPhlAn 4. Nature Biotechnology. 2023;41(11):1633–1644.

31. Lin H, Peddada SD. Multigroup analysis of compositions of microbiomes with covariate adjustments and repeated measures. Nature Methods. 2024;21(1):83–91.

32. Caspi R, Billington R, Keseler IM, et al. The MetaCyc database of metabolic pathways and enzymes - a 2019 update. Nucleic Acids Research. 2019;48(D1):D445–D453.

33. Beghini F, McIver LJ, Blanco-Míguez A, et al. Integrating taxonomic, functional, and strain-level profiling of diverse microbial communities with bioBakery 3. eLife. 2021;10:e65088.

34. Mallick H, Rahnavard A, McIver LJ, et al. Multivariable association discovery in population-scale meta-omics studies. PLoS Comput Biol. 2021;17(11):e1009442.

35. Tostivint I, Royer N, Nicolas M, et al. Spectrum of mutations in cystinuria patients presenting with prenatal hyperechoic colon. Clin Genet. 2017;92(6):632–638.

36. Morinaga K, Kusada H, Sakamoto S, et al. Granulimonas faecalis gen. nov., sp. nov., and Leptogranulimonas caecicola gen. nov., sp. nov., novel lactate-producing Atopobiaceae bacteria isolated from mouse intestines, and an emended description of the family Atopobiaceae. International Journal of Systematic and Evolutionary Microbiology. 2022;72(10).

37. Darby TM, Owens JA, Saeedi BJ, et al. Lactococcus Lactis Subsp. cremoris Is an Efficacious Beneficial Bacterium that Limits Tissue Injury in the Intestine. iScience. 2019;12:356–367.

38. Pak CY, Fuller C, Sakhaee K, et al. Management of cystine nephrolithiasis with alpha-mercaptopropionylglycine. J Urol. 1986;136(5):1003–8.

39. Prot-Bertoye C, Lebbah S, Daudon M, et al. Adverse events associated with currently used medical treatments for cystinuria and treatment goals: results from a series of 442 patients in France. BJU Int. 2019;124(5):849–861.

40. Bai Y, Tang Y, Wang J, et al. Tolvaptan treatment of cystine urolithiasis in a mouse model of cystinuria. World J Urol. 2021;39(1):263–269.

41. Khan SR. Reactive oxygen species, inflammation and calcium oxalate nephrolithiasis. Transl Androl Urol. 2014;3(3):256–276.

42. Sigurjonsdottir VK, Runolfsdottir HL, Indridason OS, et al. Impact of nephrolithiasis on kidney function. BMC Nephrol. 2015;16:149.

43. Mulay SR, Anders HJ. Crystal nephropathies: mechanisms of crystal-induced kidney injury. Nat Rev Nephrol. 2017;13(4):226–240.

44. Han M, Zhang D, Ji J, et al. Downregulating miR-184 relieves calcium oxalate crystal-mediated renal cell damage via activating the Rap1 signaling pathway. Aging (Albany NY). 2023;15(24):14749–14763.

45. Wang B, Tan Z, She W, et al. Characterizing Chemokine Signaling Pathways and Hub Genes in Calcium Oxalate-Induced Kidney Stone Formation: Insights from Rodent Models. Biochem Genet. 2025.

46. Mulay SR, Kulkarni OP, Rupanagudi KV, et al. Calcium oxalate crystals induce renal inflammation by NLRP3-mediated IL-1β secretion. J Clin Invest. 2013;123(1):236–46.

47. Joshi S, Wang W, Peck AB, et al. Activation of the NLRP3 inflammasome in association with calcium oxalate crystal induced reactive oxygen species in kidneys. J Urol. 2015;193(5):1684–91.

48. Espino M, Font-Llitjós M, Vilches C, et al. Digenic Inheritance in Cystinuria Mouse Model. PLoS One. 2015;10(9):e0137277.

49. López de Heredia M, Muñoz L, Carru C, et al. S-Methyl-L-Ergothioneine to L-Ergothioneine Ratio in Urine Is a Marker of Cystine Lithiasis in a Cystinuria Mouse Model. Antioxidants (Basel). 2021;10(9).

50. Brosnan JT, Brosnan ME. The sulfur-containing amino acids: an overview. J Nutr. 2006;136(6 Suppl):1636s–1640s.

51. Gall WE, Beebe K, Lawton KA, et al. alpha-hydroxybutyrate is an early biomarker of insulin resistance and glucose intolerance in a nondiabetic population. PLoS One. 2010;5(5):e10883.

52. Ferrannini E, Natali A, Camastra S, et al. Early metabolic markers of the development of dysglycemia and type 2 diabetes and their physiological significance. Diabetes. 2013;62(5):1730–7.

53. Ye D, Huang J, Wu J, et al. Integrative metagenomic and metabolomic analyses reveal gut microbiota-derived multiple hits connected to development of gestational diabetes mellitus in humans. Gut Microbes. 2023;15(1):2154552.

54. Stipanuk MH, Ueki I. Dealing with methionine/homocysteine sulfur: cysteine metabolism to taurine and inorganic sulfur. Journal of Inherited Metabolic Disease. 2011;34(1):17–32.

55. Lu SC. Glutathione synthesis. Biochim Biophys Acta. 2013;1830(5):3143–53.

56. Logue BA, Maserek WK, Rockwood GA, et al. The analysis of 2-amino-2-thiazoline-4-carboxylic acid in the plasma of smokers and non-smokers. Toxicol Mech Methods. 2009;19(3):202–8.

57. Nishio T, Toukairin Y, Hoshi T, et al. Quantification of 2-aminothiazoline-4-carboxylic acid as a reliable marker of cyanide exposure using chemical derivatization followed by liquid chromatography–tandem mass spectrometry. Journal of Pharmaceutical and Biomedical Analysis. 2022;207:114429.

58. Vanholder R, Schepers E, Pletinck A, et al. The uremic toxicity of indoxyl sulfate and p-cresyl sulfate: a systematic review. J Am Soc Nephrol. 2014;25(9):1897–907.

59. Ramezani A, Raj DS. The gut microbiome, kidney disease, and targeted interventions. J Am Soc Nephrol. 2014;25(4):657–70.

60. Cernaro V, Calabrese V, Loddo S, et al. Indole-3-acetic acid correlates with monocyte-to-high-density lipoprotein (HDL) ratio (MHR) in chronic kidney disease patients. Int Urol Nephrol. 2022;54(9):2355–2364.

61. Meijers BKI, Evenepoel P. The gut–kidney axis: indoxyl sulfate, p-cresyl sulfate and CKD progression. Nephrology Dialysis Transplantation. 2011;26(3):759–761.

62. Oshima S, Shiiya S, Nakamura Y. Combined Supplementation with Glycine and Tryptophan Reduces Purine-Induced Serum Uric Acid Elevation by Accelerating Urinary Uric Acid Excretion: A Randomized, Single-Blind, Placebo-Controlled, Crossover Study. Nutrients. 2019;11(11):2562.

63. Mehta M, Goldfarb DS, Nazzal L. The role of the microbiome in kidney stone formation. Int J Surg. 2016;36(Pt D):607–612.

64. Stern JM, Moazami S, Qiu Y, et al. Evidence for a distinct gut microbiome in kidney stone formers compared to non-stone formers. Urolithiasis. 2016;44(5):399–407.

65. Knauf F, Brewer JR, Flavell RA. Immunity, microbiota and kidney disease. Nat Rev Nephrol. 2019;15(5):263–274.

66. Wang Z, Huang Y, Yang T, et al. Lactococcus cremoris D2022 alleviates hyperuricemia and suppresses renal inflammation via potential gut-kidney axis. Food Funct. 2024;15(11):6015–6027.

67. Smith PM, Howitt MR, Panikov N, et al. The microbial metabolites, short-chain fatty acids, regulate colonic Treg cell homeostasis. Science. 2013;341(6145):569–73.

68. Urbauer E, Aguanno D, Mindermann N, et al. Mitochondrial perturbation in the intestine causes microbiota-dependent injury and gene signatures discriminative of inflammatory disease. Cell Host Microbe. 2024;32(8):1347–1364.

69. Clark CS, Gnanappiragasam S, Thomas K, et al. Cystinuria: An Overview of Challenges and Surgical Management. Front Surg. 2022;9:812226.

70. Birwé H, Schneeberger W, Hesse A. Investigations of the efficacy of ascorbic acid therapy in cystinuria. Urol Res. 1991;19(3):199–201.

71. Fjellstedt E, Denneberg T, Jeppsson JO, et al. A comparison of the effects of potassium citrate and sodium bicarbonate in the alkalinization of urine in homozygous cystinuria. Urol Res. 2001;29(5):295–302.

