## Supplement for "Generation of a novel *Slc7a9*^G105R^ mutant mouse identifies new biomarkers for cystinuria"

P: +61 437 593 702.

E:

**Running headline**: *Slc7a9*^G105R^ mouse identifies cystinuria biomarkers

**Supplementary detailed methods**

**Mouse husbandry**

Mouse breeding and experiments were conducted at the Alfred Medical Research and Education Precinct, Australia (AMREP AS) Pty Ltd, a facility affiliated with Monash University. Three-week-old male and female heterozygous or homozygous *Slc7a9*^G105R^ mutant mice, weighing approximately 10-12 grams, were allocated to the experiments. C57BL/6JAusb mice were used as wild-type controls. All animals were housed under standard laboratory conditions, with a 12-h day and night cycle, and had access to food and water *ad libitum*. Mice were housed in individually ventilated cages (IVC) under specific pathogen-free, physical containment 2 (PC2) conditions. Mice were monitored daily for adverse events and provided additional supportive care as required, including placing cages in a heating chamber, providing a mashed diet, and administering 0.9% sodium chloride (Alpha Medical, NSW, Australia) or 0.1 mg/kg buprenorphine (Provet, VIC, Australia) subcutaneously.

**Generation of *Slc7a9^G105R^* mice**

*Slc7a9^G105R^* mice were generated in collaboration with the Australian National University's Australian Phenomics Facility, using CRISPR-Cas9-mediated gene editing technology as previously described (**Supplementary Figure S1a**).^1^ Mouse genomic sequences were obtained from Ensembl (Ensembl.org). Cas9 protein and the single guide RNA (sgRNA) were purchased from Integrated DNA Technology (IDT) with the target sequences and single-stranded oligonucleotide donor (ssODN) (**Supplementary Table S1**). The Cas9, sgRNA 5’-GTAGGCAGGGATGGGGCCAA AGG-3’, and ssODN were delivered into the fertilized pronucleus of the fertilized zygotes with the following concentrations: Cas9 protein (50 ng/µL) was co-injected with the sgRNA (2.5 ng/µL) and the ssODN (25 ng/µL). After the micro-injection, the zygotes were incubated overnight at 37°C in 5% CO_2_, and two-cell stage embryos were surgically transferred into the ampulla of a pseudo-pregnant (CFW/crl-C57BL/6crl) F1 female mouse. DNA was extracted from the ear punches of the mice using a crude DNA extraction protocol and subsequently amplified by PCR. The PCR products were then purified with a PCR Clean-Up System kit (Promega, Sydney, Australia) according to the manufacturer's instructions using the forward (5’-TGGACCTTTGGAGAGTGGAA-3’) and reverse (5’-AGGCTGCACACACATACTCT-3’) *Slc7a9* target primers (**Supplementary Table S1**). Sanger sequencing was performed in the Biomolecular Resource facilities at the Australian National University (**Supplementary Figure S1b**-**c**). Once *Slc7a9*^G105R^ mice were generated, they were relocated to the AMREP AS Pty Ltd, at Monash University.

**Genotyping protocol**

A mouse tail tip was collected and placed into a pre-labelled Transnetyx 96-well plate and sent to Transnetyx (Transnetyx, Memphis, Tennessee, USA) for genotyping. PCR was performed with an initial denaturation at 94°C for 10 seconds, followed by 30 cycles of amplification consisting of denaturation at 94°C for 10 seconds, annealing at 55-65°C for 30 seconds, and extension at 60°C for 1 minute, with a final extension at 72°C for 3 minutes. PCR primers:

*Slc7a9*^G105R^ Forward: CTATCTGATGGAGGCCTTTC

*Slc7a9*^G105R^ Reverse: CGATCAGGCTGGTCCAGGAG

**Diets**

Mating pairs received a standard irradiated mouse diet. Pups also received a standard irradiated mouse diet until week 3 (Specialty Feeds, Glen Forest, WA, Australia). The cysteine-controlled diet (Code# SF20-009 [Specialty Feeds]) contained 0.4% cysteine. The alpha-lipoic acid (ALA)-supplemented diet (Code# SF21-078 [Specialty Feeds]) contained 0.4% cysteine and 0.5% ALA (Sigma-Aldrich, Darmstadt, Germany). The nutrient compositions of the standard, cysteine-controlled, and ALA-supplemented diets are outlined in **Supplementary Table S2**.

**Tissue collection**

On the morning of the scheduled endpoint, mice were restrained, and gentle pressure was applied to the lower abdomen to induce urination. Urine was collected in a sterile petri dish and immediately transferred into 1.5 mL microcentrifuge tubes (SSIbio, Scientific specialties, USA) for storage at -80°C until further analysis. Following urine collection, mice were euthanized with an intraperitoneal injection of 200 mg/kg pentobarbitone (Virbac, Carros, France). Blood was collected by cardiac puncture, left to clot at room temperature for at least 30 minutes, centrifuged at 3,000 x *g* for 10 minutes at 4°C, and serum was collected and stored at -80°C. The urinary bladder was excised and suspended in phosphate-buffered saline (PBS) for subsequent micro-computed tomography (μ-CT) imaging. After imaging, the urinary bladder was opened to extract cystine calculi, and then it was fixed in 4% formaldehyde (Sigma-Aldrich) for histological evaluation. The left kidney was fixed in 4% formaldehyde (Sigma-Aldrich) for histological analysis following μ-CT imaging in a subset of samples, while the right kidney was stored at -80°C for RNA extraction.

**Micro-computed tomography**

μ-CT imaging of the mouse urinary bladder or kidney was performed using the NanoScan PET/CT machine (Mediso, Budapest, Hungary) at the Alfred Research Alliance-Monash Biomedical Imaging platform (ARA-MBI). The following μ-CT imaging parameters were used: tube voltage, 70 kVp; tube current, 680 μA; exposure time, 300 ms; 1440 projections, 1 rotation; volume size, 22 x 22 x 12 μm; voxel size (reconstructed), 11 μm^3^. μ-CT images were analyzed using 3D Slicer software (version 5.0.3, [www.slicer.org](http://www.slicer.org)). The number, surface area, and volume of urinary bladder stones were quantified across all studies.

**Kidney function**

Approximately 100 μL of serum was used to measure blood urea nitrogen (BUN; Catalogue# 98-11070-01) using individual cassettes on a Catalyst One Veterinary Chemistry Analyzer from IDEXX Laboratories (IDEXX Rydalmere, NSW, Australia). *In vivo* kidney function was measured using transdermal glomerular filtration rate (*t*GFR) at 8 weeks. Mice were anesthetized via 4% isoflurane and maintained briefly on 1-2% isoflurane using a facemask. Fur was removed from the dorsal side of the mouse using an electric razor and depilation cream (Reckitt Benckiser, Sydney, Australia) one day prior to the *t*GFR measurement. *t*GFR was measured the following day by anesthetizing the mouse as previously described^2^ and attaching a mini-transdermal GFR monitor (MediBeacon GmbH, Mannheim, Germany) to the right flank muscle using surgical tape.^2^ A heating pad was applied to the animals’ tails to visualize the prominent vein and ensure accurate intravenous injection. An intravenous bolus of 50 mg/kg of fluorescein isothiocyanate (FITC)-sinistrin (Catalogue# 29389090 [MediBeacon GmbH]) was administered via the tail vein in one smooth injection. Animals were relocated into individual cages for 90 minutes whilst fluorescence decay was measured. Data were read from the device using MB Lab 2 software and analyzed using MB Studio 2 software, employing a three-compartment model with linear baseline correction to obtain the FITC-sinistrin half-life.

**Histology**

Urinary bladders and kidneys were fixed in 4% formaldehyde solution (Sigma-Aldrich), followed by paraffin embedding and sectioning. Transverse or longitudinal sections, 4 μm, were prepared and stained at the Monash Histology Platform (Clayton, Melbourne). Hematoxylin and eosin (H&E) staining was used for urinary bladder sections, while kidney sections were stained using Periodic Acid-Schiff-Alcian Blue (PAS-AB). Representative sections were captured using a Nikon Eclipse E-200 Trinocular Microscope equipped with Mosaic2.3exe software (version 2.3.0.0, Nikon Instruments Inc.). Images were acquired at 4x or 10x magnification, with high magnification regions imaged at 40x. All images were scale-calibrated using ImageJ software (version 1.53).

**Spironolactone administration**

Spironolactone (37.5 mg/kg/day, catalogue# S3378 [Sigma-Aldrich]) was dissolved in 0.1% ethanol (Sigma-Aldrich) in drinking water.^3, 4^ Mice received a working concentration of spironolactone in 150 mL of drinking water in black bottles. Vehicle-treated mice received 0.1% ethanol in 150 mL of drinking water in black bottles. Water bottles were changed every Monday, Wednesday, and Friday from weeks 3 to 9.

**Gene expression analysis**

*RNA extraction*

RNA was extracted using TRI Reagent (Sigma-Aldrich). Each extraction step was carried out at 4°C. The whole right kidney was physically homogenized in a TissueLyser II^TM^ (Qiagen, Clayton, VIC, Australia) using 5 mm stainless steel beads in a single tube containing 400 µL of PBS through 50 oscillations per second for 3 minutes. 200 μL of the kidney homogenate was used for protein extraction, and 1 mL of TRI Reagent was added to the remaining 200 μL of the kidney homogenate, which underwent a further 50 oscillations for 2 minutes until completely homogenized. The RNA sample was vortexed and then incubated at room temperature for 5 minutes, followed by centrifugation at 12,000 x *g* for 10 minutes. The supernatant was added to 250 μL chloroform (Sigma-Aldrich), incubated at room temperature for 10 minutes, and centrifuged at 12,000 x *g* for 15 minutes. Clear supernatant was extracted and mixed with 500 μL isopropanol (Sigma-Aldrich). The RNA sample tube was incubated at room temperature for 10 minutes and centrifuged at 12,000 x *g* for 10 minutes. The supernatant was discarded, and the RNA pellet was washed twice with 1 mL of 70% ethanol (Sigma-Aldrich). At the final step, the supernatant was discarded, and the pellets were briefly air-dried before resuspending them in 100 μL UltraPure RNase-Free H_2_O (Thermo-Fisher Scientific, North Ryde, VIC, Australia). RNA was stored at -80°C.

*Reverse transcription*

RNA concentration was measured using a NanoDrop^TM^ 2000c spectrophotometer (Thermo-Fisher Scientific), and RNA was diluted in RNase-Free H_2_O (Thermo-Fisher Scientific) to make the final concentration 125 ng/mL (1000 ng concentration of loading RNA). 8 μL of the RNA sample and no template control (NTC) were added to the RT plate, followed by the addition of 2 μL of DNase I Mix (Sigma-Aldrich). The plate was sealed and mixed, then incubated at room temperature for 15 minutes. 1 μL of DNase I Stop solution (Sigma-Aldrich) was added, the plate sealed, and mixed. The RT plate was put in the Bio-Rad T100^TM^ Thermal cycler (Bio-Rad Laboratories, South Granville, NSW, Australia) and incubated at 65°C for 10 minutes. 2 μL of Random Primers (Thermo-Fisher Scientific) and dNTP mix (Bioscience, Austin, Texas, USA) was added followed by plate sealing and mixing. The RT plate was returned to the thermal cycler to incubate at 65°C for another 5 minutes. Furthermore, 7 μL of MMLV MasterMix (Thermo-Fisher Scientific) was added to the sample, then sealed and gently mixed, before being returned to the thermal cycler to incubate at 37°C for 2 minutes. After incubation, 1 μL of MMLV (Thermo-Fisher Scientific) was added, followed by sealing and mixing. The plate was put back in the thermal cycler set for incubation at 25°C for 10 minutes, 37°C for 50 minutes and then 70°C for 15 minutes. After the completion of reverse transcription, the plate was spun down, and 79 μL of RNase-free H_2_O (Thermo-Fisher Scientific) was added. The final volume of cDNA was stored in 200 μL RNA-grade tubes at -30°C.

*RT^2^ profiler PCR arrays*

RT^2^ SYBR Green (Catalogue# 330520 [Qiagen]) based real-time RT^2^ profiler PCR arrays were performed using the QuantStudio 6 Flex Real-Time PCR system (Applied Biosystems, Waltham, MA, USA). The mouse inflammatory response and autoimmunity (Gene Globe ID# PAMM-077ZE) gene expression panel was used with the RT^2^ profiler PCR array kit (Catalogue# 330231 [Qiagen]) according to the manufacturer's instructions.

*Targeted quantitative real-time polymerase chain reaction (qPCR)*

Kidneys were harvested and stored at -80°C before processing for reverse transcription and qPCR to assess inflammatory gene expression. Samples were analyzed in duplicate, and cDNA amplification was performed using a QuantStudio 6 Flex Real-Time PCR system (Applied Biosystems). Gene expression was quantified using SYBR Green PowerUp Mix assays (Thermo-Fisher Scientific). The relative gene expression was determined using the 2^−ΔΔCT^ method, with normalization to the geometric mean of the housekeeping gene, glyceraldehyde-3-phosphate dehydrogenase (*Gapdh*). Gene expression levels were then normalized to control mice.

**Amino acid quantification**

*Amino acid measurement in serum*

Arginine, cystine, lysine, and ornithine were quantified by comparison against an eight-point external calibration curve prepared in a matrix. The amino acid calibration curves were prepared by constructing a series of diluted standard solutions of accurately known concentration, which were then added to 8 µL of serum and extracted as described below. For arginine, cystine, and lysine, the curve was constructed using a commercially available mixture of amino acids, Standard H (Thermo-Fisher Scientific). For ornithine, a stock solution was prepared from the hydrochloride salt. For arginine, lysine, and ornithine, these curves show large positive intercepts consistent with significant endogenous concentrations of these amino acids in the serum samples. Consequently, the amino acid concentrations were calculated using the slope of the curves and setting the intercept to zero. Serum samples were extracted by first thawing them on ice and then transferring 10 µL to a fresh tube. 80 µL of ice-cold extraction solvent (1:1 acetonitrile:methanol) was added before mixing for 30 minutes at 4°C. The sample was centrifuged at 20,000 x *g* for 10 minutes at 4°C before transferring the supernatant to a sample vial for Liquid Chromatography-Mass Spectrometry (LC-MS) analysis.

*Amino acid measurement in urine*

Arginine, cystine, lysine, and ornithine were quantified by comparison against an eight-point external calibration curve prepared in a matrix. The amino acid calibration curves were prepared by constructing a series of diluted standard solutions of accurately known concentration, which were then added to 20 µL of 100-fold diluted matrix urine and extracted as described below. In the case of arginine, cystine and lysine, the diluted standard solutions were prepared from a commercially available mixture of amino acids, Standard H (Thermo-Fisher Scientific). For ornithine, a stock was prepared from the hydrochloride salt. Urine samples were mixed by repeated aspiration before transferring 5 µL of urine to a fresh tube and adding 495 µL of LC-MS grade water to obtain a 100-fold diluted sample. 40 µL of this diluted sample or 20 µL of the calibration curve sample was extracted. To each sample, three volumes (120 µL samples/60 µL calibration curve) of ice-cold 75% acetonitrile were added, followed by mixing for 30 minutes at 4°C. The sample was centrifuged at 20,000 x *g* for 10 minutes at 4°C before transferring the supernatant to a sample vial for LC-MS analysis.

*LC-MS analysis*

A Dionex RSLC3000 ultra-high-performance liquid-chromatography (UHPLC) coupled to a Q-Exactive Plus Orbitrap MS (Thermo-Fisher Scientific) was used. Samples were analyzed using hydrophilic interaction liquid chromatography (HILIC), following a previously published method.^5^ The chromatography utilized an iHILIC®-(P) Classic 150 x 4.6 mm, 5 µM, 200Å (Hilicon, Sweden) (25°C). A gradient elution was performed using 20 mM ammonium carbonate as mobile phase A (A) and acetonitrile as mobile phase B (B). The linear gradient involved decreasing the proportion of acetonitrile from 80% at 0 minutes, 50% at 15 minutes, 5% at 18 minutes, held at 5% until 21 minutes, followed by a return to 80% at 24 minutes, which was maintained until 32 minutes to re-equilibrate the column. The flow rate was maintained at 500 μL/minute. Samples were kept in the autosampler (6°C) and 10 μL was injected for analysis. MS was performed at 70,000 resolutions operating in rapid switching positive (4 kV) and negative (3.5 kV) mode electrospray ionization (capillary temperature 300°C; sheath gas flow rate 50; auxiliary gas flow rate 20; sweep gas 2; probe temp 120°C). Peaks corresponding to the extracted ion chromatograms of the protonated amino acids were then integrated in TraceFinder 4.1 (Thermo-Fisher Scientific).

**Untargeted metabolomics**

*Serum and urine metabolite extraction*

20 μL serum and 50 μL urine were thawed and resuspended in a 4:1 ratio of methanol containing 1 µM CCTP internal standards, 5 µM BHT and 13C,15N-amino acid mixture. For urine samples, the extracted volume was adjusted based on the creatinine concentration in each sample. Creatinine concentration was measured using Creatinine Assay—The Creatinine Companion (Ethos Biosciences, USA). Blanks were made with water and extracted alongside. Samples were mixed at 4°C for 60 minutes under agitation and centrifuged at 20,817 x *g* for 10 minutes at 4°C. For serum, the supernatant was transferred to LC-MS vials for acquisition. The urine samples were evaporated using a SpeedVac until dry and resolubilized using 80% MeOH containing 1 µM CAPS, CHAPS and PIPES, 15N, 13C-Amino acid mixture, and 5 µM BHT. QC samples were made by pooling 10 μL from each sample.

*LC-MS data acquisition*

The metabolomic profiles of serum and urine samples were determined separately using untargeted LC-MS. LC-MS was acquired on a Dionex Ultimate® 3000 RS high-performance liquid chromatography (HPLC) system (Thermo Fisher, USA) coupled with a Q-Exactive Orbitrap mass spectrometer (Thermo Fisher, USA) at the Monash Metabolomics and Proteomics Platform (Melbourne, Australia). Chromatographic separation was performed on a ZIC-pHILIC column (5 µM, polymeric, 150 × 4.6 mm, SeQuant, Merck) using a mobile phase composed of 20 mM ammonium carbonate (A) and acetonitrile (B). The gradient program started at 80% acetonitrile, reduced to 50% over 15 minutes, then further reduced to 5% over the next 3 minutes. This low organic phase was maintained for a 3 minutes washing step, followed by re-equilibration at 80% for 8 minutes to prepare the column for the next injection. The flow rate was set to 0.3 mL/minute, and the column temperature to 40°C. The total run time was 32 minutes with an injection sample volume of 10 µL. The mass spectrometer operated in both positive and negative polarity modes, switching at a resolution of 35,000 at 200 m/z, with a detection range of 85 to 1,275 m/z in full-scan mode. The heated electrospray ionization source (HESI) was set to 3.5 kV voltage for positive mode and 4.0 kV for negative mode. Sheath gas was set to 50, aux gas to 20 arbitrary units, capillary temperature to 300°C, and probe heater temperature to 120°C. Each sample set was randomized to account for system drift. A mixture of pure, authentic standards containing approximately 400 metabolites was acquired as separate injections and used to confirm retention times (RT).

*Metabolomics raw data processing*

Raw LCMS data was processed using the metabolome-lipidome-MSDIAL pipeline (https://zenodo.org/records/15701972), including MS-DIAL (version 5.3), the MassBank database (Accessed in September 2024), the pmp R package (version 1.18.0) and the Human Metabolome Database (HMDB, version 5). In MS-DIAL, parameters included MS tolerance = 0.002 Da, min peak height = 100,000, RT time tolerance = 2 minutes, gap filling = TRUE. Using pmp in R, features were filtered for intensities 10-fold higher than blanks, and samples with more than 80% missing features were removed. Features with more than 90% missing values in quality control samples and exceeding a maximum standard deviation of 30% in the quality control samples were filtered out. The filtered dataset was normalized using probabilistic quotient normalization, imputed for missing values using the random forest method, and scaled using the generalized logarithm method. Features in this dataset were further annotated using the HMDB database (mass tolerance = 0.002 Da). All successfully annotated features were manually screened in MS-DIAL to remove features with poor-quality spectra. Duplicate features identified across both negative and positive acquisition modes were selected based on the highest gap-filling percentage and signal-to-noise ratio. This resulted in 2,757 unique metabolites for downstream analysis.

*Metabolomics data analysis*

PCA was computed using the prcomp function of the stats R package (version 4.4.2). Permutational analysis of variance was computed using the vegan R package on Euclidean distances (version 2.6.10). The Limma R package was used to detect differentially abundant metabolites between groups (version 3.62.2). *P* values were adjusted using the False Discovery Rate (FDR) multiple testing correction method, with statistical significance considered at a threshold of 0.05 and a minimum log fold change of 0.5. Correlations were computed using the Hmisc R package (version 5.2.3), with a *P* value < 0.05 considered statistically significant. Plots were plotted using the ggplot R package (version 3.5.2). Heatmaps were plotted using the ComplexHeatmap package (version 2.22.0). Analyses were performed using R (version 4.4.2). Metabolism pathways were identified by mapping individual metabolites to the KEGG database.

**Antibiotic administration**

An antibiotic cocktail of enrofloxacin (10 mg/kg/day, Enrotril, Troy Laboratories, NSW, Australia) and amoxicillin/clavulanic acid (1 mg/kg/day, Augmentin, Aspen Pharmacare, VIC, Australia) was provided to mice in their drinking water.^6^ 3.75 mL of enrofloxacin was diluted in 150 mL of drinking water, while 2.36 g of Augmentin was first dissolved in 10.33 mL of drinking water and then further diluted 1:100 in 150 mL of drinking water. Vehicle-treated mice received 150 mL of drinking water. Water bottles were changed every Monday, Wednesday, and Friday from week 3 to 9.

**Fecal metagenomics**

*Bacterial DNA isolation from fecal pellets*

Fecal pellets were collected at 9 weeks of age from vehicle-treated heterozygote *Slc7a9*^G105R^, homozygote *Slc7a9*^G105R^, and antibiotic-treated homozygote *Slc7a9*^G105R^ mice. DNA was extracted from these samples using the QIAmp PowerFecal Pro DNA Kit (Qiagen) according to the manufacturer’s instructions. One fecal pellet was placed into the PowerBead Pro Tube with 800 μL of Solution CD1, followed by horizontal vortexing at maximum speed using the Vortex Mixer 24 (Omni International, Kennesaw, Georgia, USA). The tubes were then centrifuged at 15,000 x *g* for 1 minute, and the resulting supernatant was transferred to a 2 mL microcentrifuge tube. Next, 200 μL of Solution CD2 was added, briefly vortexed (VM1, Ratek Instruments Pty. Ltd., Boronia, VIC, Australia), and centrifuged as described in the previous step. Up to 700 μL of the supernatant was transferred to another 2 mL microcentrifuge tube, where 600 μL of Solution CD3 was added and briefly vortexed. 650 μL of the lysate was then loaded onto an MB Spin Column, centrifuged, and the flow-through discarded; this step was repeated until all the lysate had passed through the column. After placing the column in a 2 mL collection tube, 500 μL of solution EA was added, centrifuged, and the flow-through was discarded. Following this, 500 μL of solution CD5 was added to the column, centrifuged, and the flow-through discarded. The column was then transferred to a fresh 2 mL collection tube and centrifuged at 16,000 x *g* for 2 minutes. Finally, the column was transferred to a 1.5 mL elution tube, where 75 μL of solution CD6 was added, followed by centrifugation at 15,000 x *g* for 1 minute. DNA concentration was quantified using a NanoDrop^TM^ 2000c spectrophotometer (Thermo-Fisher Scientific) and samples were stored at 4°C until use in metagenomic analysis.

*Sample preparation*

Fecal DNA samples were quantified and assessed for quality using the Qubit system (Invitrogen, USA). Sample integrity was verified with the Agilent 2100 Bioanalyzer system (Agilent Technologies, Germany), which was then utilized to construct Illumina Nextera XT sequencing libraries according to the manufacturer's protocol (Illumina Inc., USA). Finally, the libraries were sequenced on the MGI DNBSEQ G400 platform (MGI Tech Co., Ltd., China) following the manufacturer’s guidelines.

*Processing and data analysis*

Paired-end 2x100 bp raw reads were quality checked using FastQC^7^ and depleted of host (Mus musculus, GRCm38 Ensembl release 102) and PhiX (GCA_002596845.1) reads using Bowtie2 (v2.4.2)^8^ retaining unmapped reads for downstream analysis. Reads were quality-filtered and trimmed for adapter sequences using fastp (v0.23.2).^9^ The MetaPhlAn4 software (v4.0.6)^10^ with the default ChocoPhlAn database (vOct22) was used to estimate the taxonomic abundance of species in each sample. To identify differentially abundant species between heterozygous and homozygous samples, the ANCOM-BC2 method was employed, incorporating structural zero detection^11^ and FDR correction for multiple testing.^12^ The PERMDISP2 method was implemented for heterogeneity analysis using the vegan::betadisper function.^13^ Bray-Curtis beta dispersion was calculated and plotted using mia and miaViz packages.^14^ For functional and pathway enrichment analysis, HUMAnN (v3.8)^15^ was used to identify orthologs against UniRef90 (uniref90_201901b_full) and to quantify community pathway abundance using the MetaCyc database.^16^ A linear model implemented in MaAsLin2^17^ was used to identify differentially abundant pathways quantified by HUMAnN, utilizing log-transformed copies-per-million (CPM) values, without a minimum prevalence or abundance cut-off, and FDR correction for multiple testing. We used R (R Core Team, 2023) and the Bioconductor packages *mia*^18^ and *miaViz*^19^ for plotting and statistical analysis of the metagenomic data.

**Cohousing of *Slc7a9*^G105R^ mice**

Male homozygote *Slc7a9*^G105R^ and C57BL/6JAusb wild-type mice of the same age were weaned onto a cysteine-controlled diet at 3 weeks of age and cohoused from weeks 3 to 9 (6 weeks). Additionally, two groups, male wild-type and male homozygote *Slc7a9*^G105R^ mice, were weaned onto a cysteine-controlled diet at week 3 and served as controls for the cohousing experiment.

**Supplementary figures**

**Supplementary Figure S1**

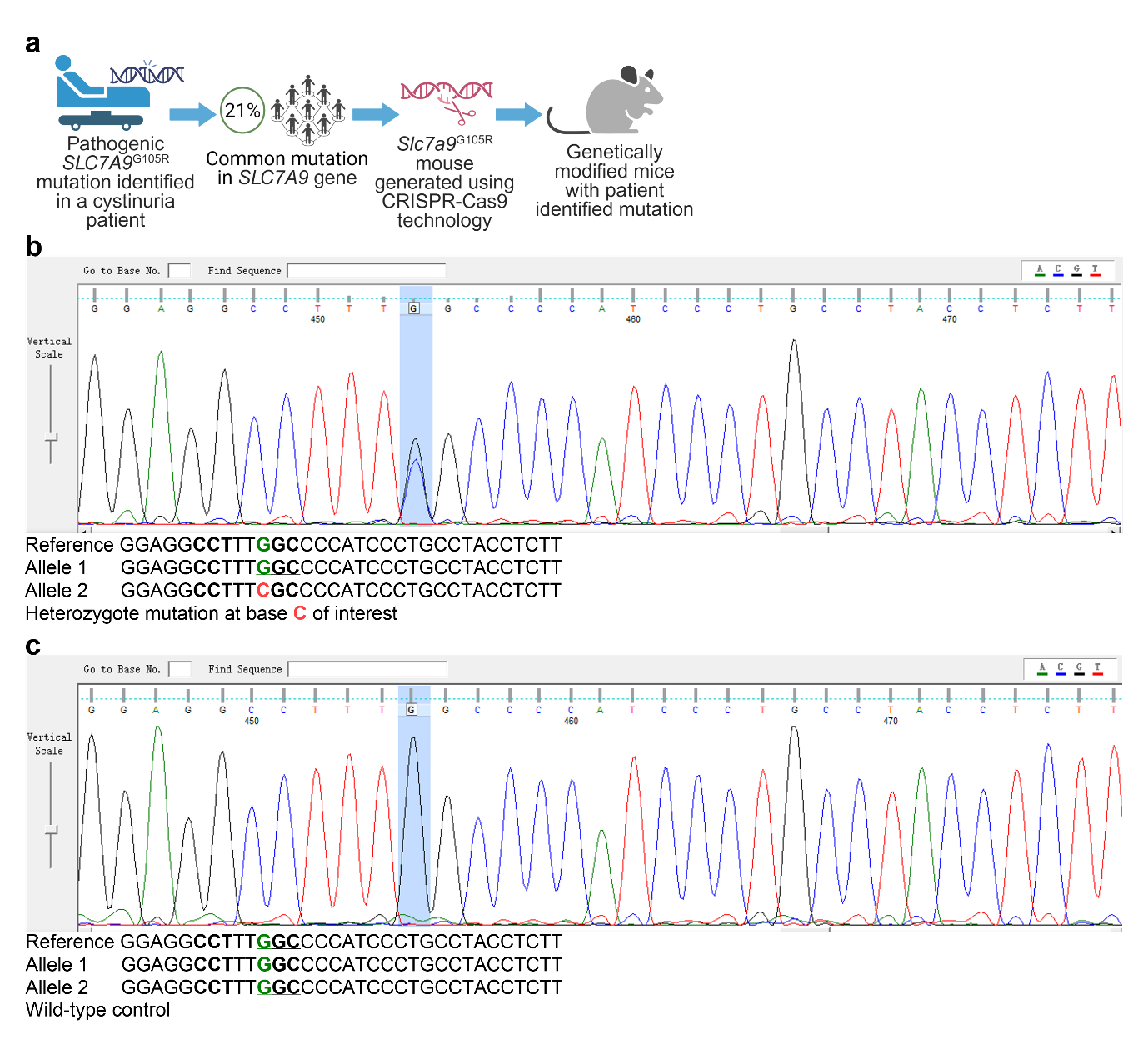

**Figure S1. Generation of a novel *Slc7a9*^G105R^ mutant mouse with CRISPR-Cas9 gene editing.** A novel mouse model of cystinuria was developed using CRISPR/Cas9 technology to replicate a single-point mutation identified in a pediatric cystinuria patient with early-onset cystinuria. (**a**) Schematic representation of *Slc7a9*^G105R^ mouse generation. (**b**-**c**) Sanger sequencing chromatograms confirming the single-point mutation in the *Slc7a9* gene. The schematic was created using Biorender (License # Bhatt, N., 2025; https://BioRender.com/b3b0kvs).

**Supplementary Figure S2**

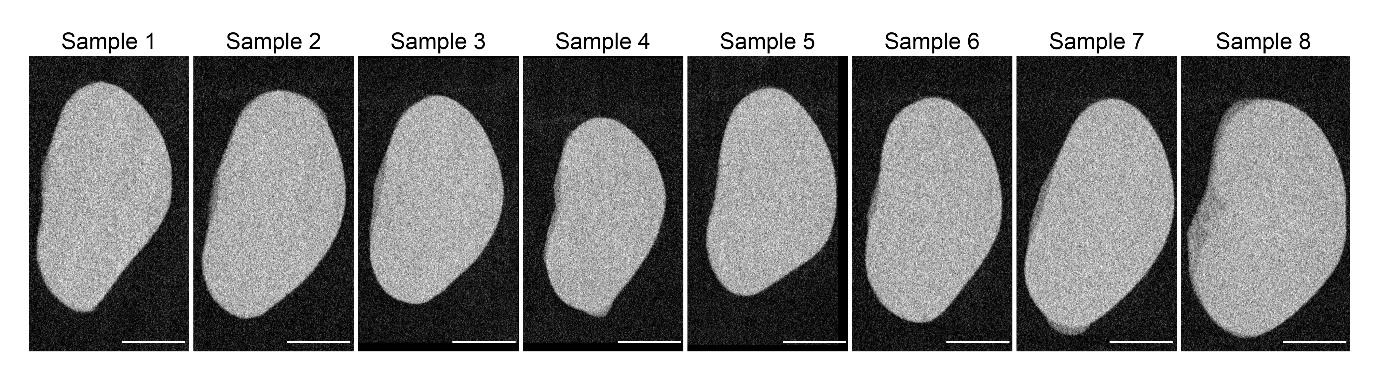

**Figure S2.** **Kidney µ-CT scans show no cystine stones in *Slc7a9*^G105R^ mice.** Representative µ-CT kidney images from male homozygote *Slc7a9*^G105R^ mice at 9 weeks of age. The scale bar represents 2.5 mm.

**Supplementary Figure S3**

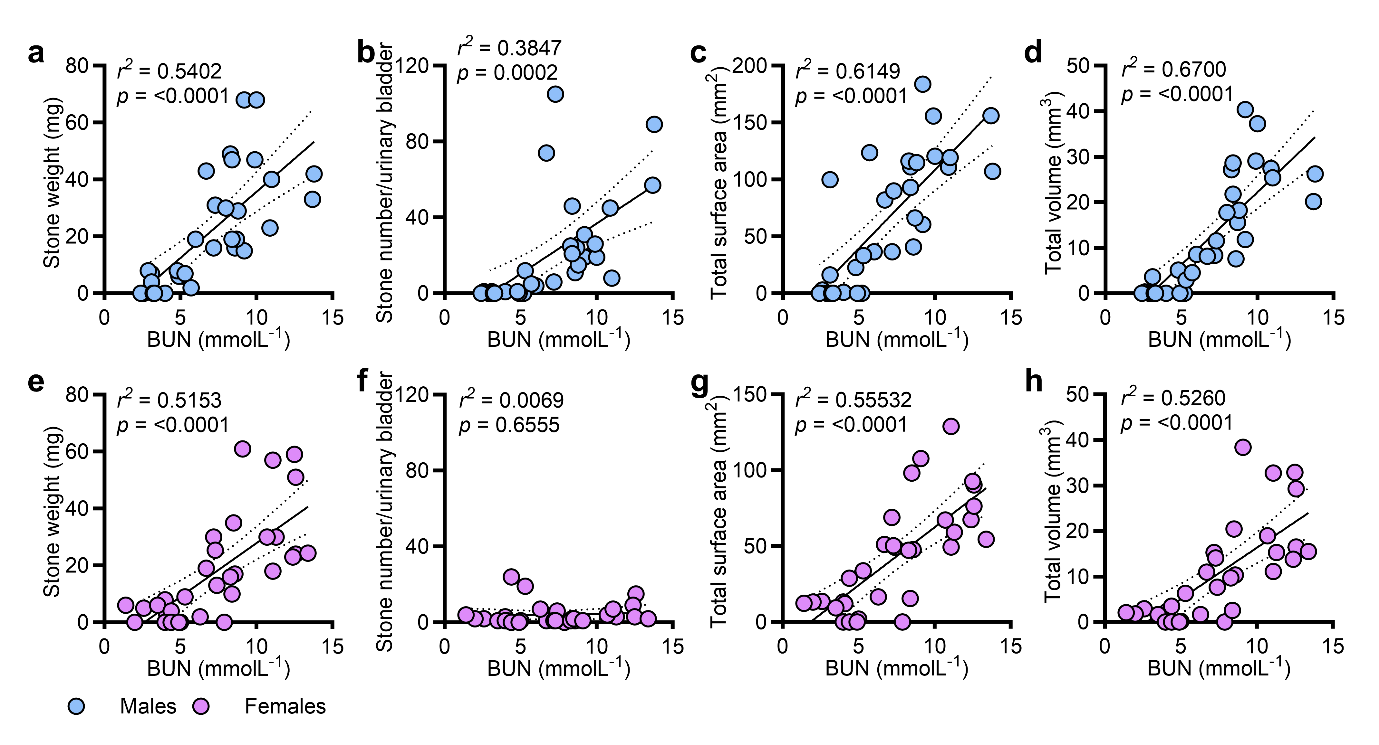

**Figure S3. Blood urea nitrogen correlates with cystine stone burden.**
Homozygote *Slc7a9*^G105R^ male (top panels) and female (bottom panels) mice (aged 3-9 weeks) with varying stone burdens were analyzed. Blood urea nitrogen (BUN) positively correlated with (**a, e**) cystine stone weight, (**b, f**) number, (**c, g**) surface area, and (**d, h**) volume. Correlations were assessed by simple linear regression and reported as *r²* and *P* values.

**Supplementary Figure S4**

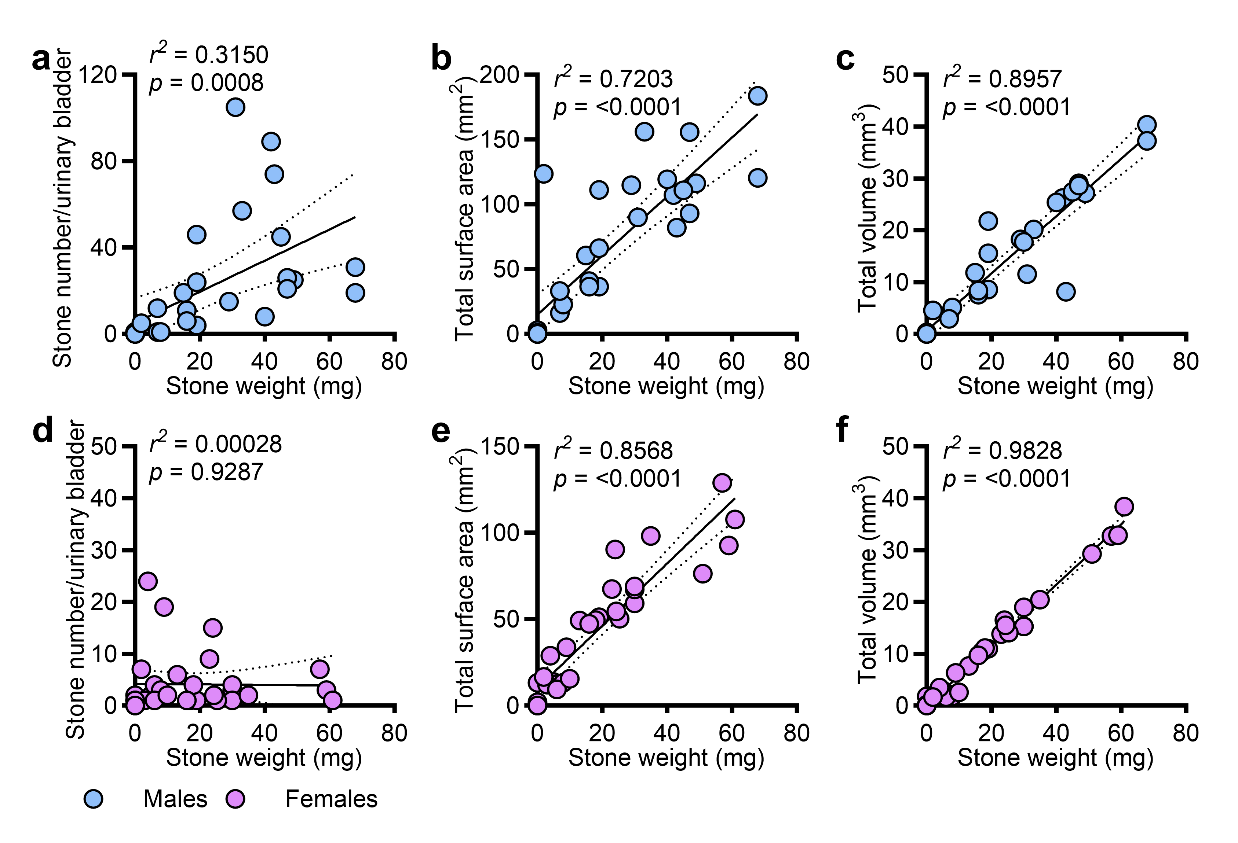

**Figure S4. µ-CT analysis of cystine stones shows a positive correlation with physical stone weight.** Homozygote *Slc7a9*^G105R^ male (top panels) and female (bottom panels) mice (aged 3-9 weeks) with varying stone burdens were analyzed. Cystine stone weight positively correlated with (**a, d**) cystine stone number, (**b, e**) surface area, and (**c, f**) volume. Correlations were assessed by simple linear regression and reported as *r²* and *P* values.

**Supplementary Figure S5**

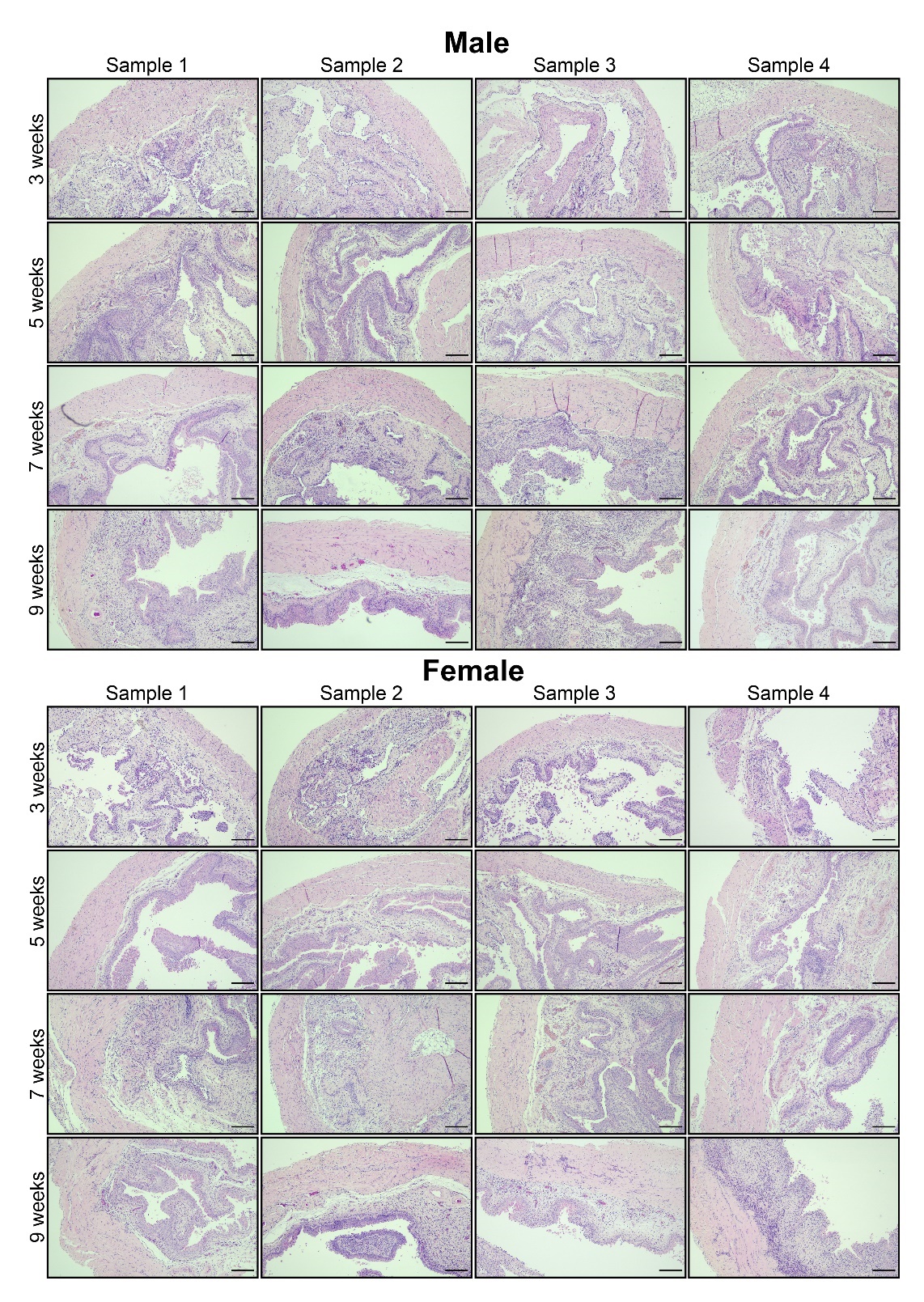

**Figure S5. Increase urinary bladder histopathology in homozygote *Slc7a9*^G105R^ mice over time.** Hematoxylin and eosin (H&E)-stained sections of the urinary bladder in male (upper panels) and female (lower panels) homozygote *Slc7a9*^G105R^ mice at 3, 5, 7, and 9 weeks of age. The scale bar represents 200 µm, with each group comprising representative images from four independent mice.

**Supplementary Figure S6**

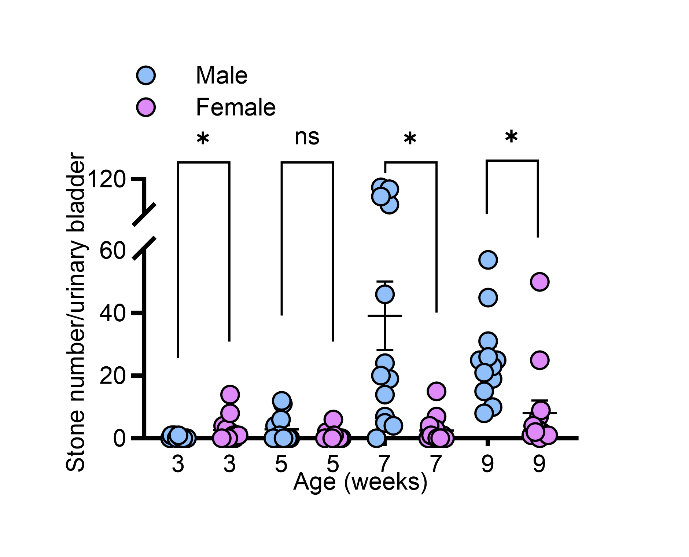

**Figure S6. Male *Slc7a9*^G105R^ mice have increased numbers of urinary bladder cystine stones at 7 and 9 weeks compared to females.** Quantification of urinary bladder cystine stones in male and female homozygote *Slc7a9*^G105R^ mice at 3, 5, 7, and 9 weeks of age. Data were obtained using µ-CT imaging and analyzed with 3D Slicer software. Data are presented as the mean ± the standard error of the mean (SEM). Statistical differences were calculated using a non-parametric unpaired Mann-Whitney t-test. * denotes *P* value < 0.05. ns denotes not significant.

**Supplementary Figure S7**

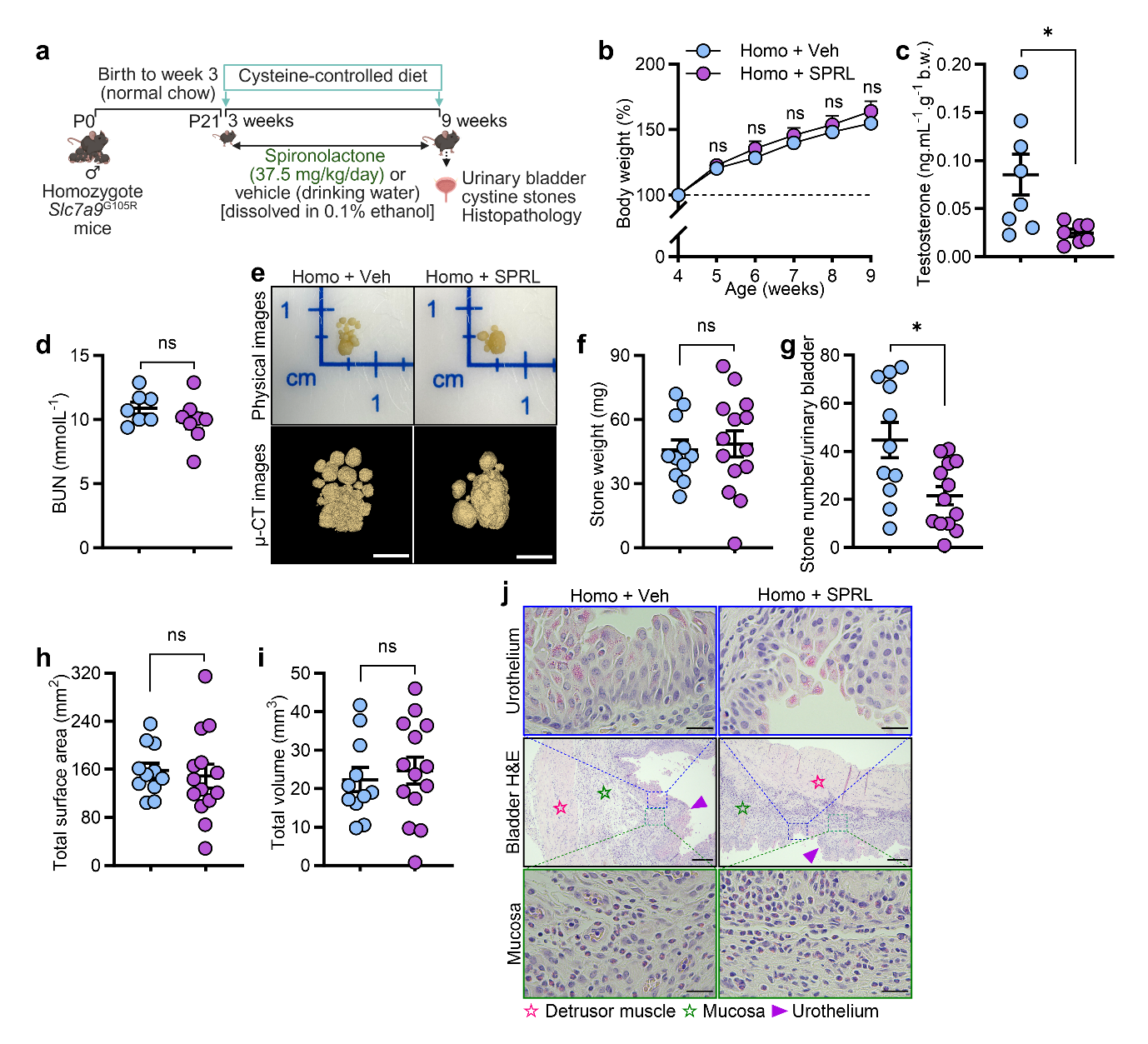

**Figure S7. Spironolactone treatment suppressed testosterone levels and reduced cystine stone numbers in male *Slc7a9*^G105R^ mice.** Cystine stone growth and urinary bladder histopathology were assessed in homozygote *Slc7a9*^G105R^ mice treated with spironolactone and in vehicle-treated controls. (**a**) Schematic representation of the experimental model. (**b**) Body weight change. (**c**) Serum testosterone levels (*n* = 8 per group). (**d**) Serum blood urea nitrogen (BUN) levels (*n* = 7-8 per group). (**e**) Representative physical and µ-CT cystine stone images. (**f**) Cystine stone weight, (**g**) number, (**h**) total surface area, and **(i)** total stone volume (*n* = 11-14 per group). (**j**) Representative hematoxylin and eosin (H&E)-stained bladder sections. Symbols ☆, ☆, and ► indicate the urinary bladder detrusor muscle, mucosa, and urothelium, respectively. The scale bar indicates 2.5 mm for µ-CT stone images, 200 µm for the bladder H&E sections, and 25 µm for the magnified sections of urothelium and mucosa. Data were obtained from two independent experiments and are presented as the mean ± the standard error of the mean (SEM). Statistical differences were calculated using a two-way ANOVA with Tukey’s multiple comparisons test (**b**) and non-parametric unpaired Mann-Whitney t-test (**c-d**, **f**-**i**). * denotes *P* value < 0.05. ns denotes not significant. The schematic was created using Biorender (License # Bhatt, N., 2025; https://BioRender.com /l8om05t).

**Supplementary Figure S8**

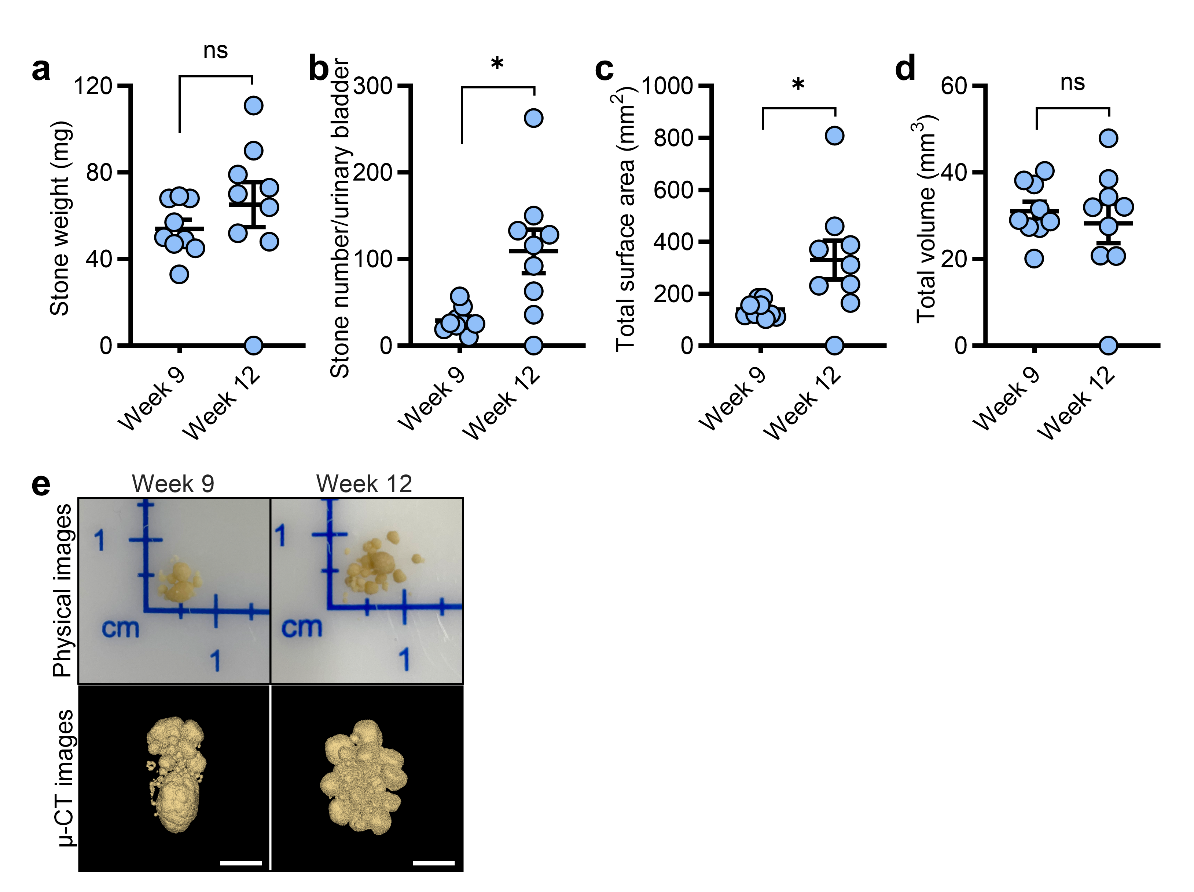

**Figure S8. Cystine stone growth plateaus after 9 weeks in *Slc7a9*^G105R^ mice.**
Cystine stone parameters were compared between 9- and 12-week-old homozygote *Slc7a9*^G105R^ mice. (**a**) Stone weight, (**b**) number, (**c**) surface area, and (**d**) volume (*n* = 9 per group, from two independent experiments). (**e**) Representative physical and µ-CT images of cystine stones at 9 and 12 weeks; scale bar = 2.5 mm. Data are presented as the mean ± the standard error of the mean (SEM). Statistical comparisons were performed using the non-parametric unpaired Mann-Whitney t-test; * denotes *P* value < 0.05. ns denotes not significant.

**Supplementary Figure S9**

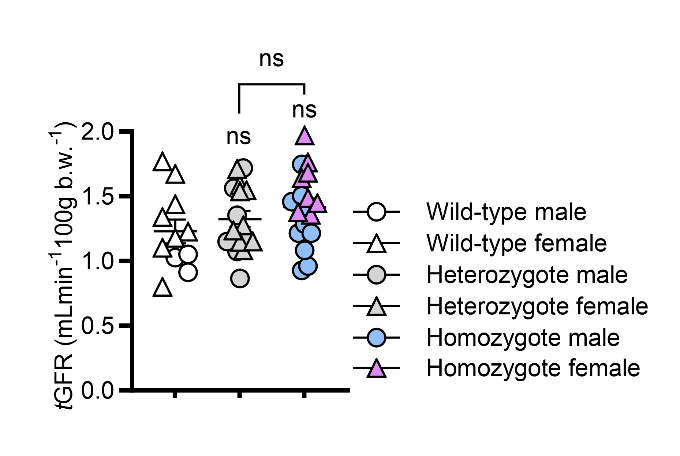

**Figure S9. Transdermal glomerular filtration rate is normal in homozygote *Slc7a9*^G105R^ mice.** At 8 weeks of age, *in vivo* real-time transdermal glomerular filtration rate (*t*GFR) measurements were performed in wild-type, heterozygote, and homozygote *Slc7a9*^G105R^ mice. Data are presented as the mean ± the standard error of the mean (SEM). Statistical differences were calculated using a non-parametric unpaired Mann-Whitney t-test, with ns denoting not significant.

**Supplementary Figure S10**

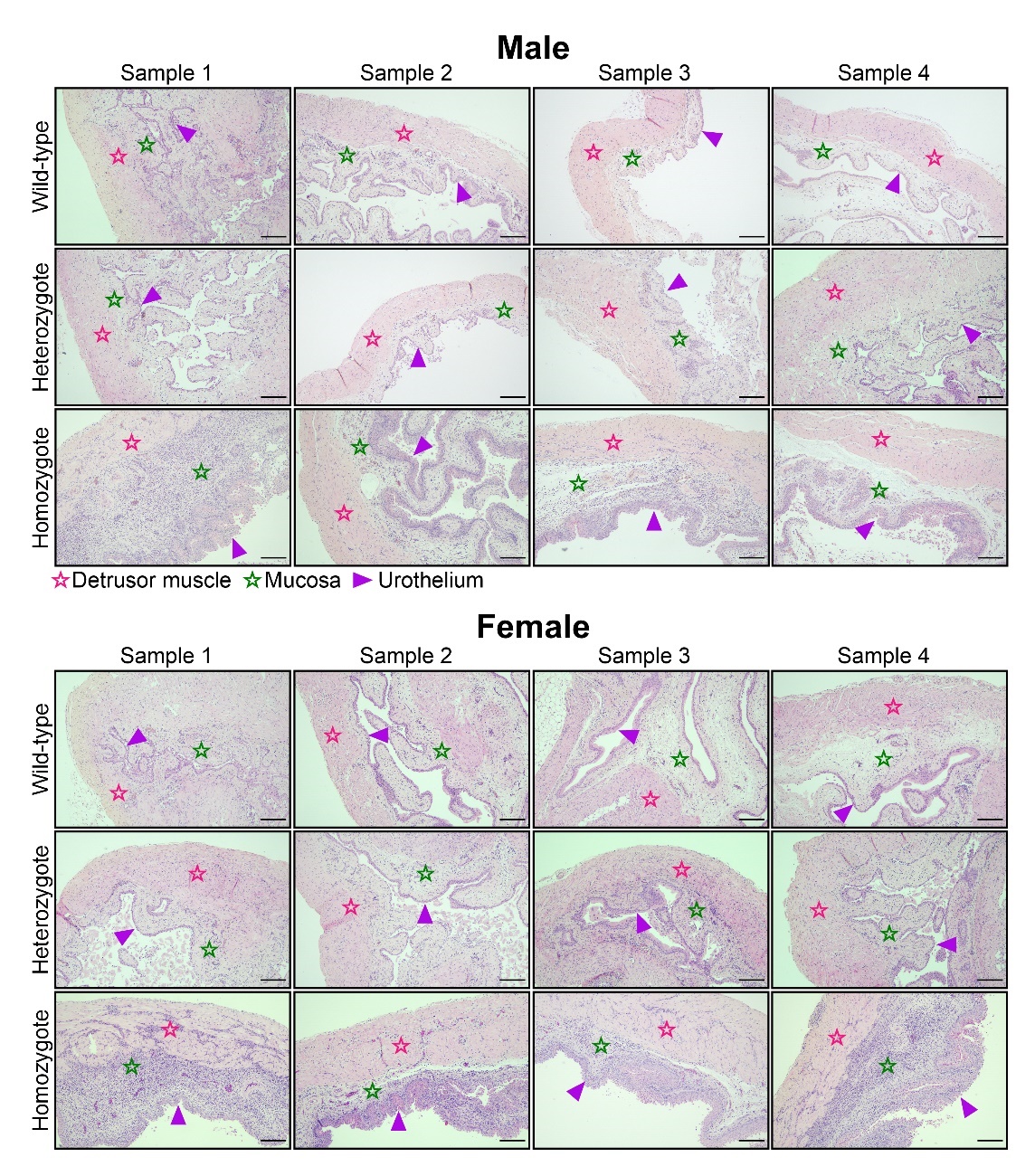

**Figure S10. Profound urinary bladder histopathology in homozygote *Slc7a9*^G105R^ mice compared to wild-type and heterozygote controls.** Hematoxylin and eosin (H&E)-stained sections of the urinary bladder from male and female wild-type, heterozygote, and homozygote *Slc7a9*^G105R^ mice at 9 weeks of age. Symbols ☆, ☆, and ► indicate the urinary bladder detrusor muscle, mucosa, and urothelium, respectively. The scale bar represents 200 µm, with each group comprising representative images from four independent mice.

**Supplementary Figure S11**

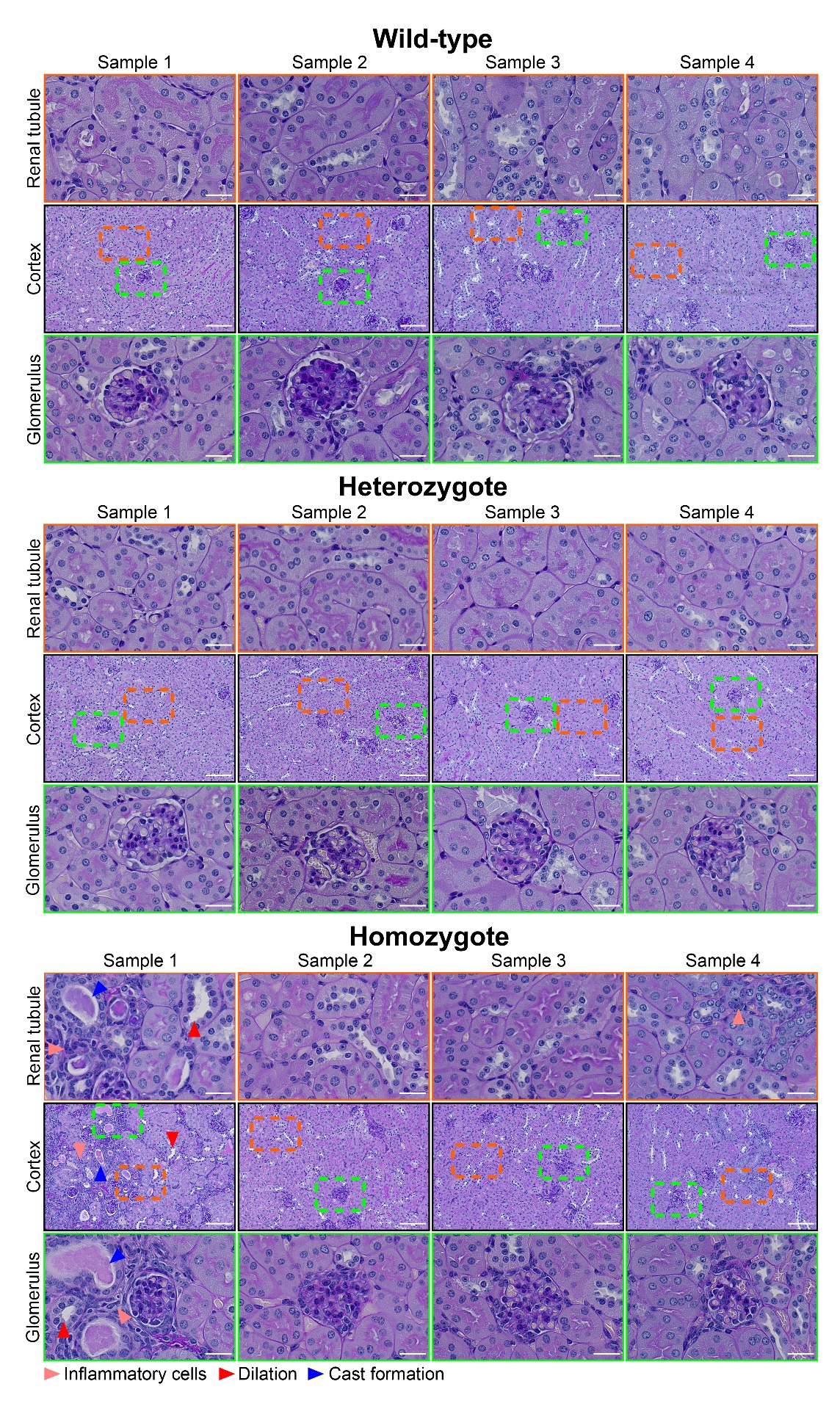

**Figure S11. Minimal kidney histopathology in homozygote *Slc7a9*^G105R^ mice compared to wild-type and heterozygote controls.** Periodic acid-Schiff-Alcian blue (PAS-AB)-stained sections from male wild-type, heterozygote, and homozygote *Slc7a9*^G105R^ mice at 9 weeks of age. Symbols ►, ►, and ► indicate inflammatory cells, renal tubular dilation, and tubular cast formation, respectively. The scale bar represents 100 µm for the cortex and 25 µm for the renal tubule and glomerulus, with each group comprising representative images from four independent mice.

**Supplementary Figure S12**

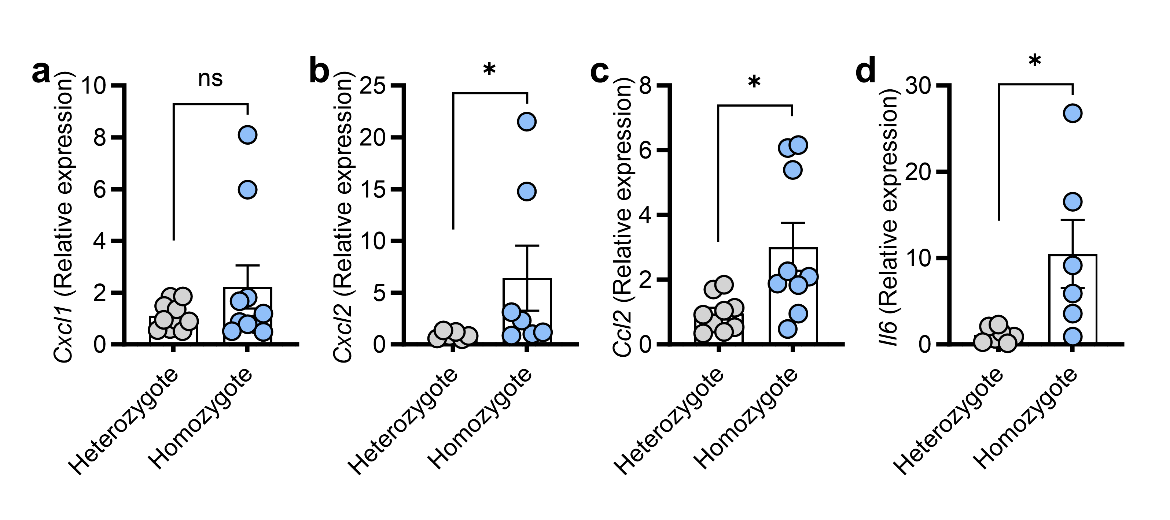

**Figure S12. Increased inflammatory gene expression in the kidneys of homozygote *Slc7a9*^G105R^ mice.** Inflammatory gene expression was assessed in the kidneys of 9-week-old male heterozygote and homozygote *Slc7a9*^G105R^ mice. (**a**) Expression of chemokine (C-X-C motif) ligand 1 (*Cxcl1*), (**b**) *Cxcl2*, (**c**) chemokine (C-C motif) ligand 2 (*Ccl2*), and (**d**) interleukin 6 (*Il6*) were analyzed by quantitative PCR (qPCR) in kidney tissue (*n* = 6-10 per group). Glyceraldehyde-3-phosphate dehydrogenase (*Gapdh*) was used as a housekeeping gene. Gene expression data are presented as the mean ± the standard error of the mean (SEM), with values expressed relative to the mean of the control group. Statistical analysis was conducted using a non-parametric unpaired Mann-Whitney t-test. * denotes *P* value < 0.05. ns indicates not significant.

**Supplementary Figure S13**

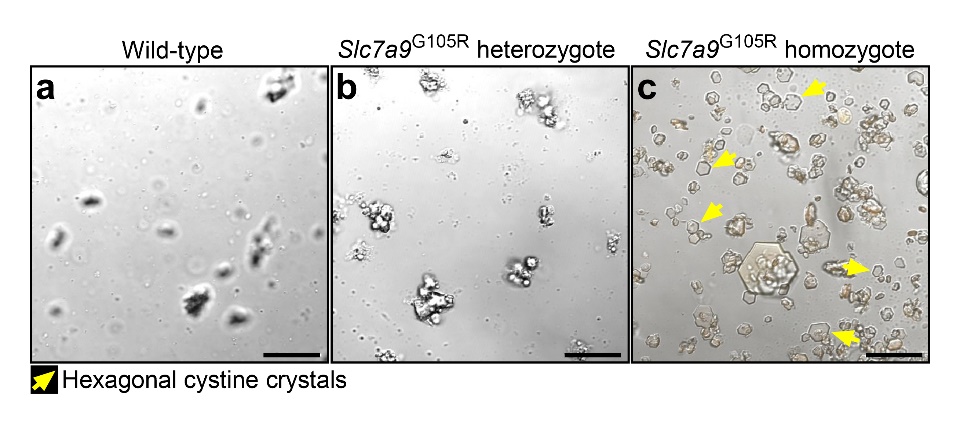

**Figure S13. Microscopy reveals classical urinary cystine crystals in homozygote *Slc7a9*^G105R^ mice.** Images were acquired using Apo LWP 40x, 1.5 NA water immersion objective on a Nikon A1R confocal microscope (Nikon, Japan). Representative microscopic images of urine from (**a**) wild-type, (**b**) heterozygote *Slc7a9*^G105R^, and (**c**) homozygote *Slc7a9*^G105R^ mice at 9 weeks of age. Yellow arrows indicate classical hexagonal urinary cystine crystals. The scale bar depicts 25 µm.

**Supplementary Figure S14**

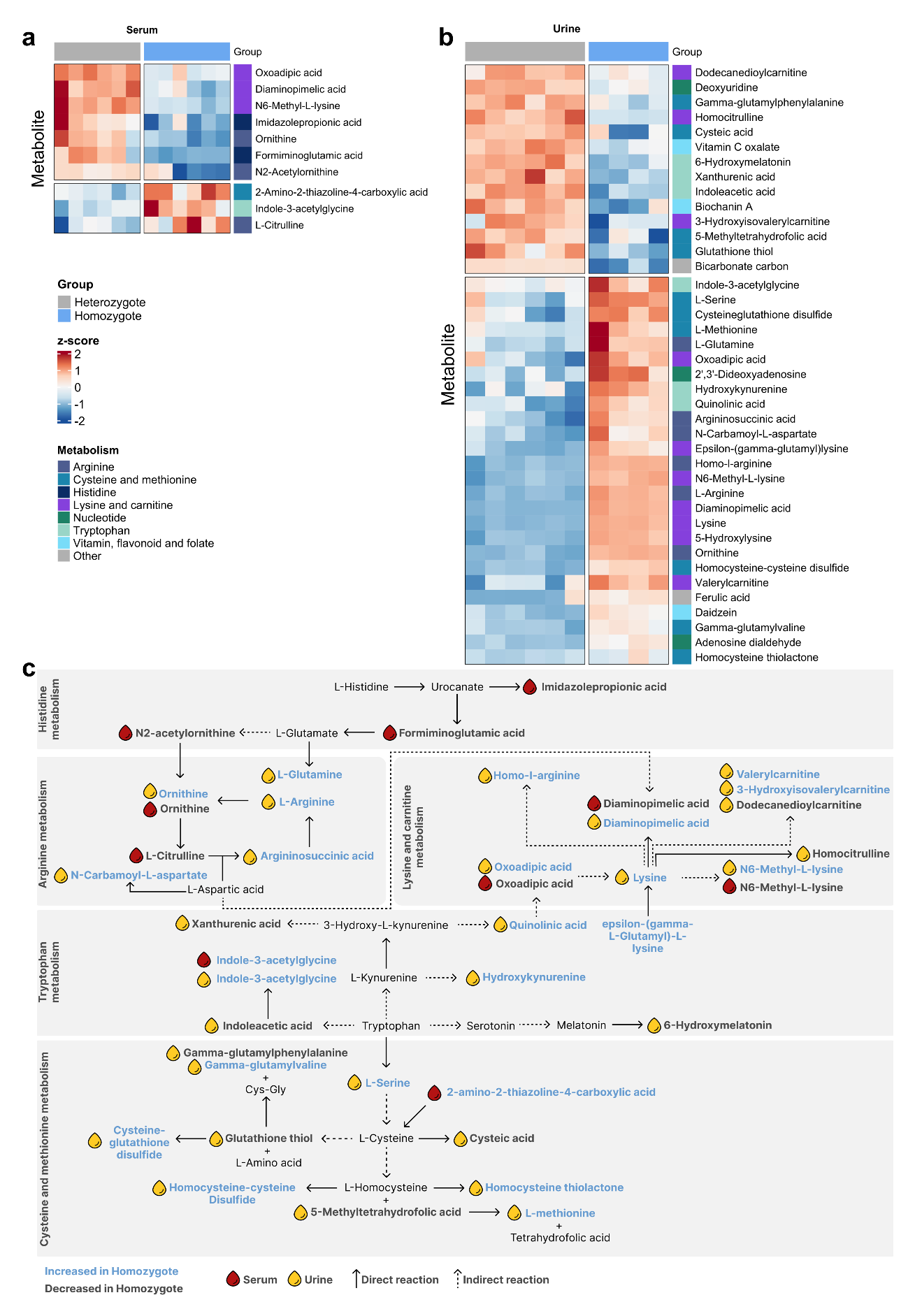

**Figure S14.** **Distinct serum and urine metabolite profiles are present in heterozygote and homozygote *Slc7a9*^G105R^ mice.** (**a-b**) Heatmap of differentially abundant metabolites in homozygous and heterozygous groups mapped to metabolic pathways of interest. Metabolites are colored according to their change in abundance (z-score) in homozygotes compared to heterozygotes. (**c**) Metabolic pathway network showing differentially abundant metabolites mapped to their respective pathways according to the KEGG database, along with interactions across pathways. Arrows outline whether metabolites are connected by a direct (solid line) or indirect (dotted line) reaction. Icons mark metabolites that are significant in the blood (red color) or the urine (yellow color). Bold black text denotes that the metabolite is decreased in homozygotes, and bold blue text denotes that a metabolite is increased in homozygotes compared to heterozygote controls. Metabolites with no icons reflect connecting metabolites that were not significant.

**Supplementary Figure S15**

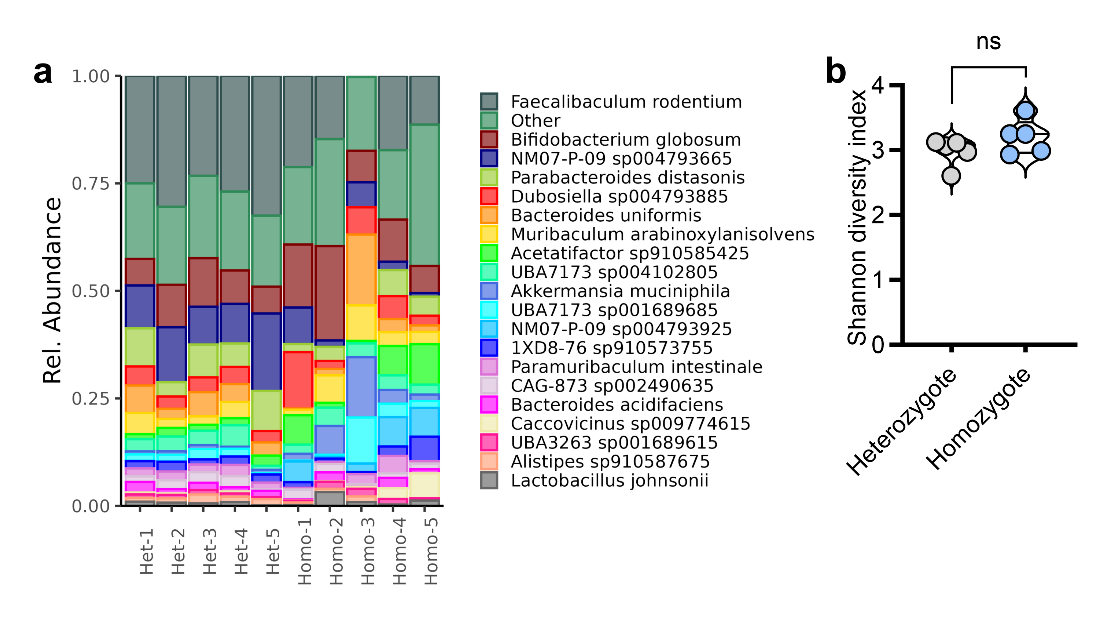

**Figure S15.** **Fecal microbiome diversity is unaltered in homozygote *Slc7a9*^G105R^ mice.** (**a**) Relative abundance of the top 20 microbial species in fecal samples. (**b**). Alpha diversity was assessed using the Shannon index in heterozygote and homozygote *Slc7a9*^G105R^ mice. Statistical analysis was performed using a non-parametric unpaired Mann-Whitney t-test. ns indicates not significant.

**Supplementary Figure S16**

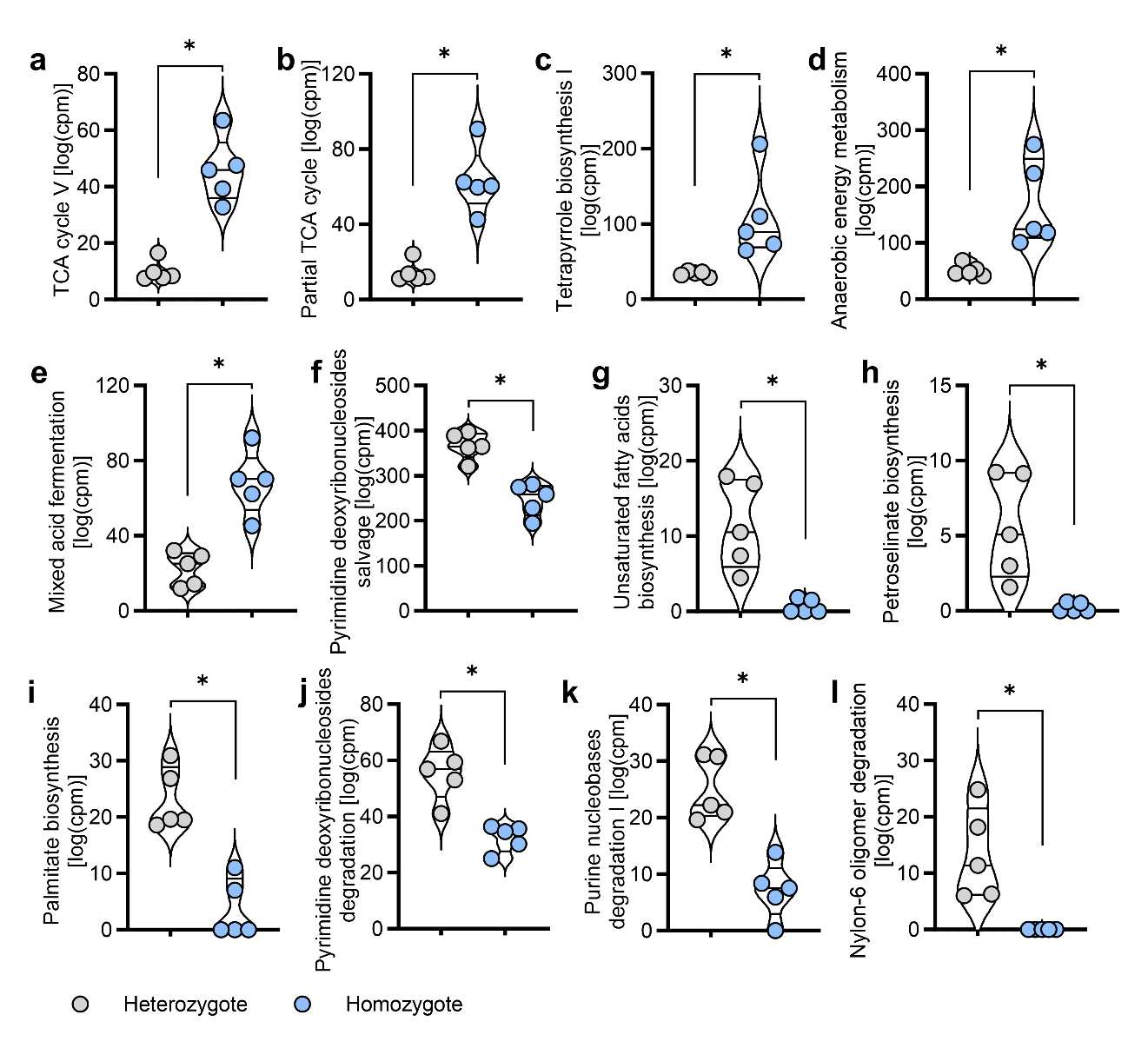

**Figure S16.** **Functional fecal microbiota pathways are significantly altered in homozygote *Slc7a9*^G105R^ mice.** A total of 12 significantly altered functional pathways were identified. (**a**-**e**) The TCA cycle V, partial TCA cycle, tetrapyrrole biosynthesis I, anaerobic energy metabolism, and mixed acid fermentation pathways were enriched in the gut microbiome community of homozygote *Slc7a9*^G105R^ mice compared to heterozygote controls. (**f-l**) Pyrimidine deoxyribonucleosides salvage, unsaturated fatty acids biosynthesis, petroselinate biosynthesis, palmitate biosynthesis, pyrimidine deoxyribonucleosides degradation, purine nucleobases degradation I, and nylon-6 oligomer degradation pathways were at lower abundance in homozygote *Slc7a9*^G105R^ mice compared to heterozygote controls. Data are representative of *n = 5* per group and are presented as the mean ± the standard error of the mean (SEM). Statistical significance was calculated using a non-parametric unpaired Mann-Whitney t-test. * indicates *P* value < 0.05.

**Supplementary Figure S17**

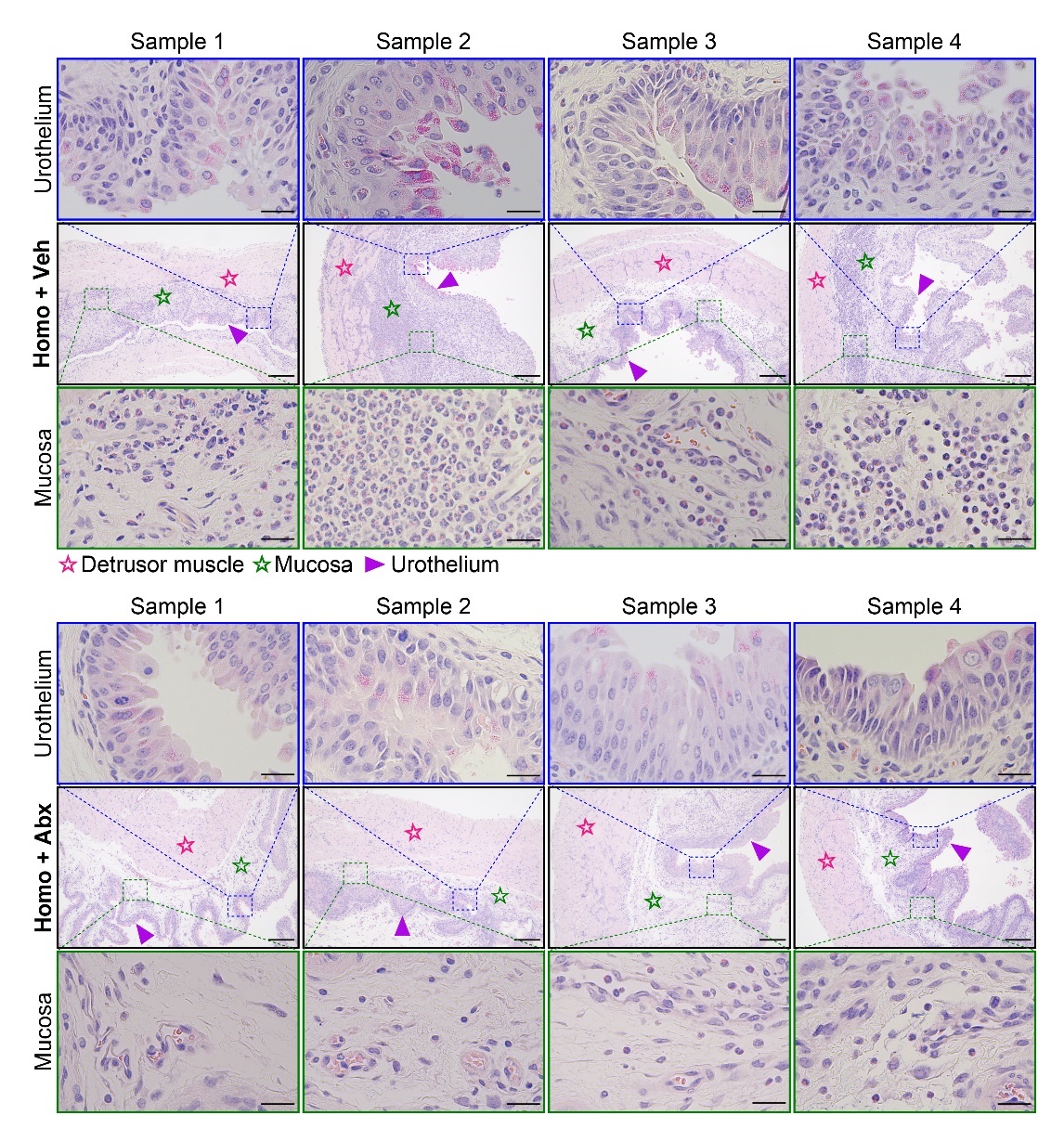

**Figure S17. Antibiotic treatment visually reduces inflammatory cell infiltration into the urinary bladder whilst maintaining urothelium integrity.** Hematoxylin and eosin (H&E)-stained bladder sections from vehicle-treated (upper panels) and antibiotic-treated (lower panels) *Slc7a9*^G105R^ mice. Symbols ☆, ☆, and ► indicate the urinary bladder detrusor muscle, mucosa, and urothelium, respectively. The scale bar represents 200 µm for the bladder section and 25 µm for the magnified urothelium and mucosa, with each group comprising representative images from four independent mice.

**Supplementary Figure S18**

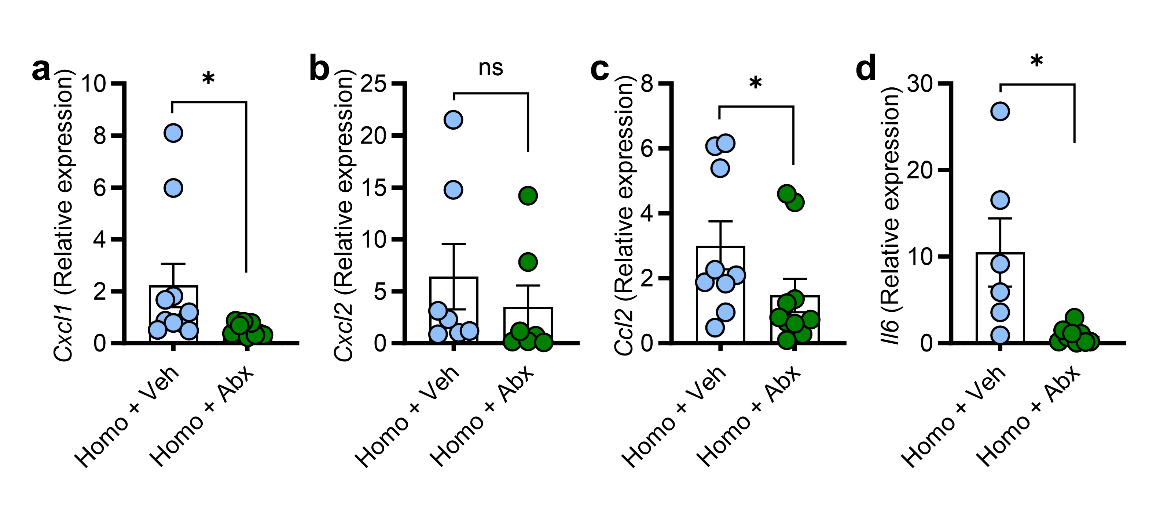

**Figure S18. Antibiotic treatment reduces the expression of inflammatory genes in the kidney.** Inflammatory genes were assessed in kidneys from vehicle-treated (Homo + Veh) and antibiotic-treated *Slc7a9*^G105R^ (Homo + Abx) mice. Expression of (**a**) *Cxcl1*, (**b**) *Cxcl2*, (**c**) *Ccl2*, and (**d**) *Il6* was assessed by quantitative PCR (qPCR) in kidney tissue (*n* = 6-10 per group). *Gapdh* was used as a housekeeping gene. Gene expression data are shown as the mean ± the standard error of the mean (SEM), with values shown relative to the mean of the control group. Statistical significance was determined using a non-parametric unpaired Mann-Whitney t-test. * denotes *P* value < 0.05. ns denotes not significant.

**Supplementary Figure S19**

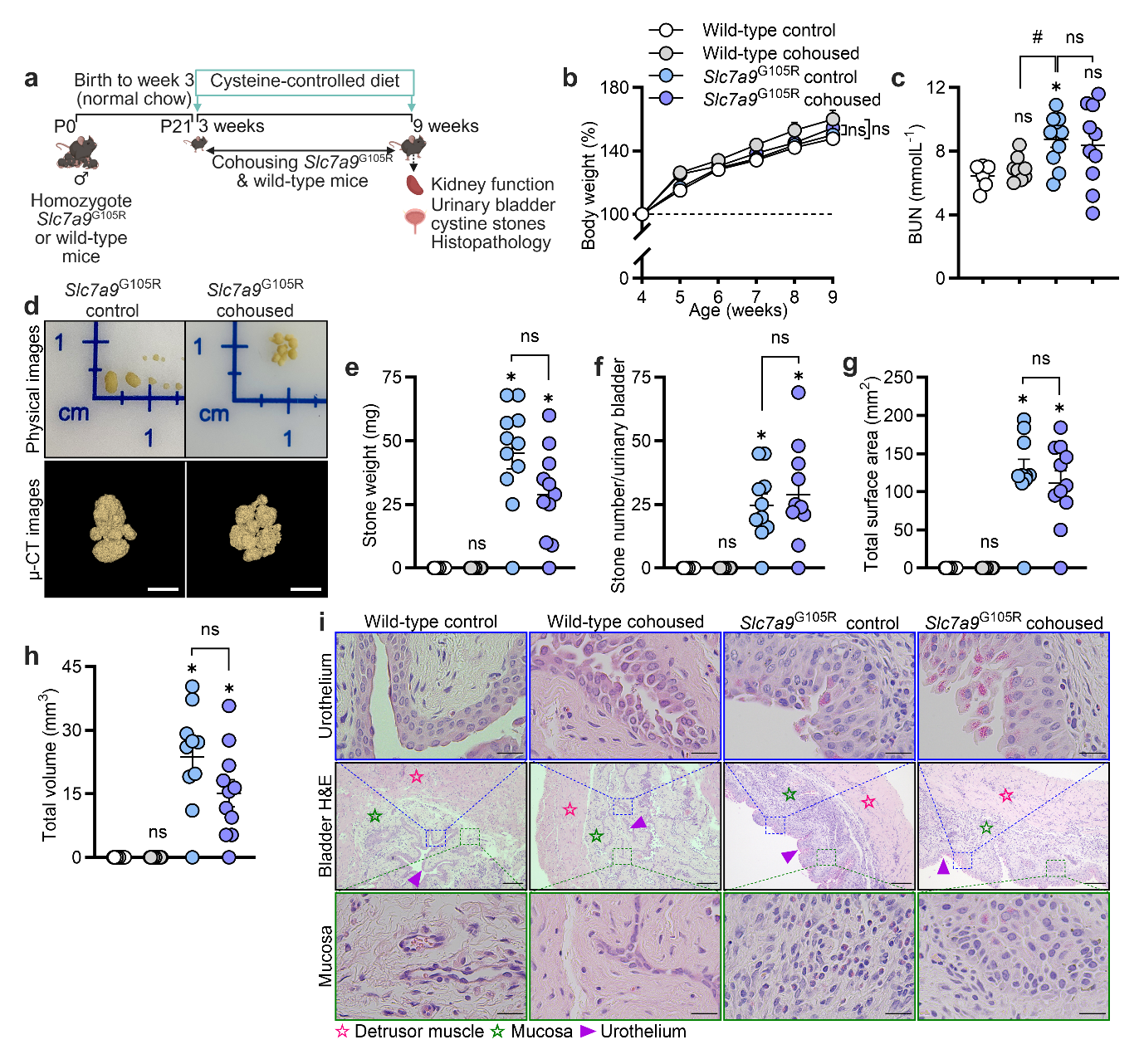

**Figure S19. Cohousing of wild-type mice with homozygote *Slc7a9*^G105R^ mice did not affect the severity of cystinuria.** Male wild-type and homozygote *Slc7a9*^G105R^ mice were cohoused for 6 weeks from weaning (3 weeks) to the endpoint (9 weeks). (**a**) Schematic representation of the experimental model. (**b**) Percentage body weight change over time. (**c**) Serum blood urea nitrogen (BUN) levels (*n* = 6 per group). (**d**) Matched representative physical and µ-CT images of urinary bladder cystine stones. (**e-h**) Quantification of urinary bladder cystine stone weight, number, total surface area, and total stone volume (*n* = 11 per group). (**i**) Representative hematoxylin and eosin (H&E)-stained bladder sections. Symbols ☆, ☆, and ► indicate the urinary bladder detrusor muscle, mucosa, and urothelium, respectively. The scale bar indicates 2.5 mm for µ-CT stone images, 200 µm for the bladder H&E sections, and 25 µm for the magnified sections of urothelium and mucosa. Data were obtained from two independent experiments and are presented as the mean ± the standard error of the mean (SEM). Statistical differences were calculated using a two-way ANOVA with Tukey’s multiple comparisons test (**b**) and a non-parametric unpaired Mann-Whitney t-test (**c**, **e**-**h**). *P* value <0.05. * denotes a significant difference compared to wild-type controls. ^#^ denotes statistical differences between two consecutive groups. ns indicates not significant. The schematic was created using Biorender (License # Bhatt, N., 2025; https://BioRender.com /85dvrga).

**Supplementary Figure S20**

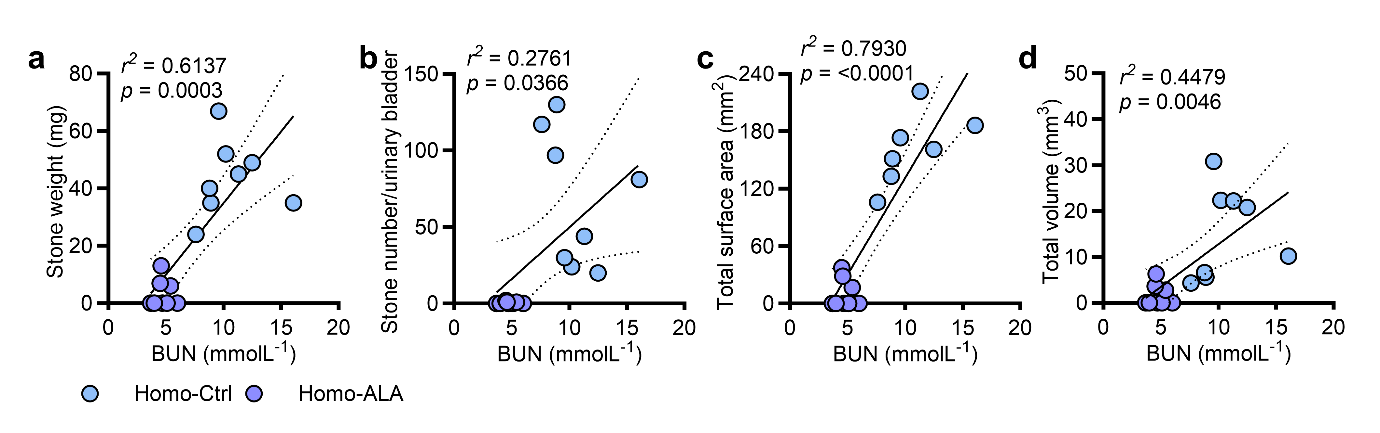

**Figure S20. Prophylactic dietary supplementation with alpha-lipoic acid reduced serum blood urea nitrogen levels, correlating with cystine stone burden.** Correlations between blood urea nitrogen (BUN) levels and cystine stone (**a**) weight, (**b**) number, (**c**) surface area, and (**d**) volume. Simple linear regression analyses were performed to evaluate the relationship between BUN levels and each stone parameter. Statistical correlations are reported as the coefficient of determination (*r²*) and the corresponding P values.

**Supplementary Figure S21**

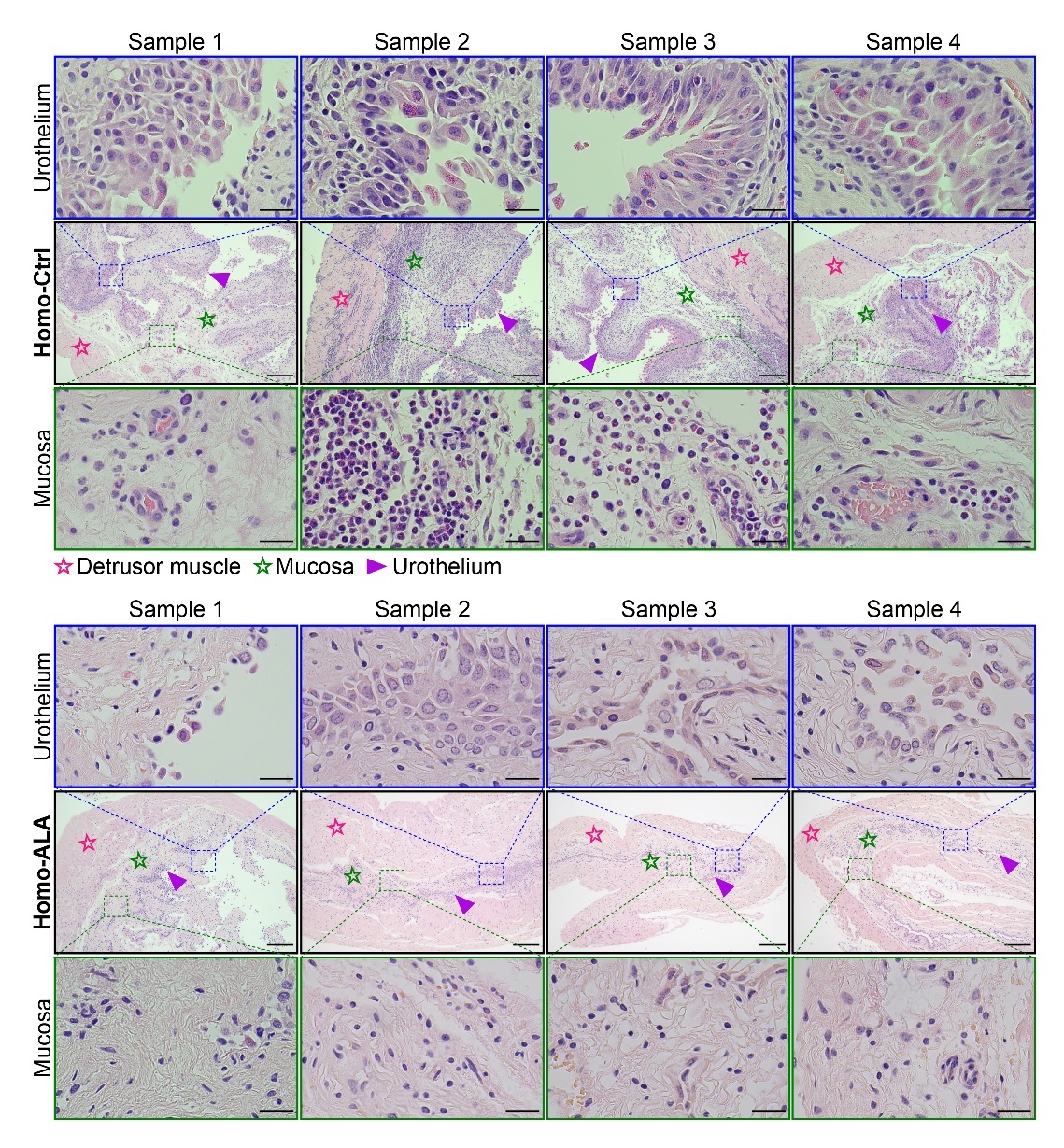

**Figure S21. Prophylactic dietary supplementation with alpha-lipoic acid reduced inflammatory cell infiltration but disrupted the urothelium.** Representative hematoxylin and eosin (H&E)-stained bladder sections from control (upper panels) and alpha-lipoic acid (ALA)-treated (lower panels) *Slc7a9*^G105R^ mice. Symbols ☆, ☆, and ► denote the detrusor muscle, mucosa, and urothelium, respectively. Scale bars represent 200 µm for the bladder sections and 25 µm for the magnified views of the urothelium and mucosa. Each group includes representative images from four independent mice.

**Supplementary Figure S22**

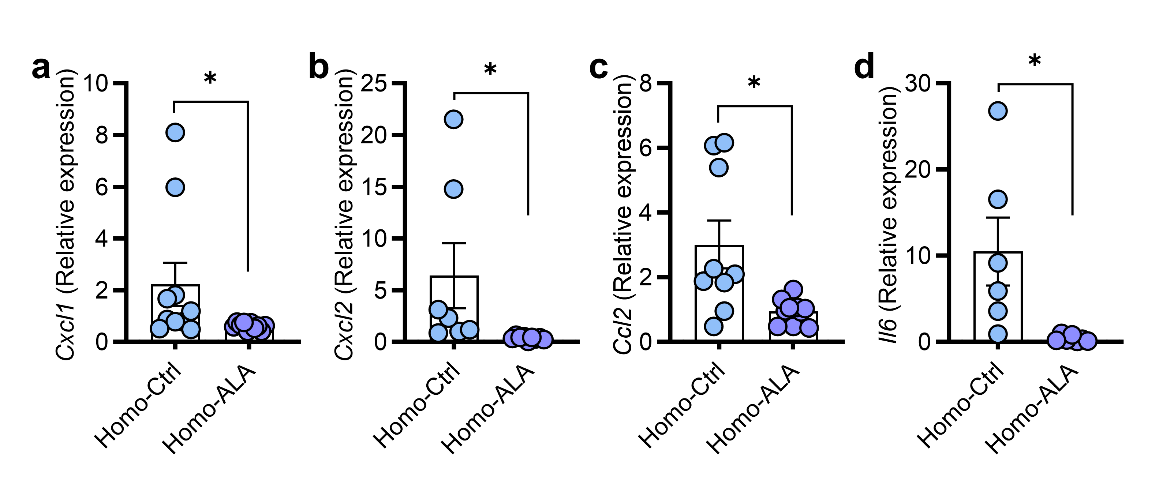

**Figure S22. Prophylactic alpha-lipoic acid supplementation attenuates renal inflammatory gene expression.** Inflammatory gene expression was analyzed in kidney tissues from control (Homo-Ctrl) and alpha-lipoic acid (ALA)-treated homozygote *Slc7a9*^G105R^ (Homo-ALA) mice. Quantitative PCR was used to assess the expression of (**a**) *Cxcl1*, (**b**) *Cxcl2*, (**c**) *Ccl2*, and (**d**) *Il6* (*n* = 6-10 per group). *Gapdh* was used as a housekeeping gene. Gene expression data are presented as the mean ± the standard error of the mean (SEM), with values expressed relative to the mean of the control group. Statistical significance was determined using a non-parametric unpaired Mann-Whitney t-test. * indicates *P* value < 0.05.

**Supplementary Figure S23**

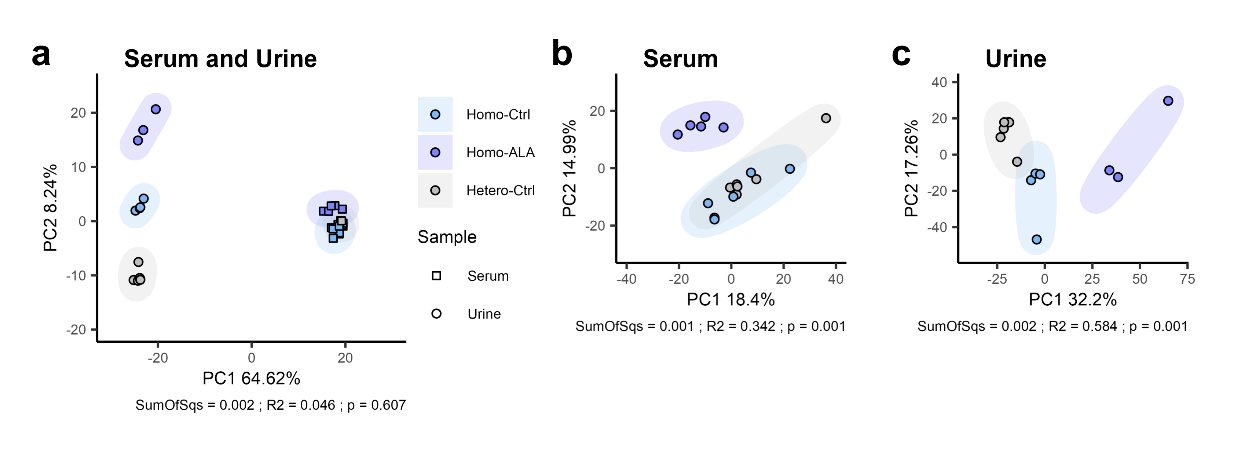

**Figure S23. Serum and urine metabolomic profiles in heterozygote, homozygote, and alpha-lipoic acid-treated homozygote *Slc7a9*^G105R^ mice.** Principal components analysis (PCA) ordination plots showing separation between genotype and treatment groups in (**a**) combined serum and urine samples, (**b**) serum alone, and (**c**) urine alone. Ellipses represent the 95% confidence interval around each group centroid. PERMANOVA results are displayed in the figure captions and include the sum of squares (SumOfSqs, effect size), R2 (proportion of variation explained by group), and associated P value.

**Supplementary Figure S24**

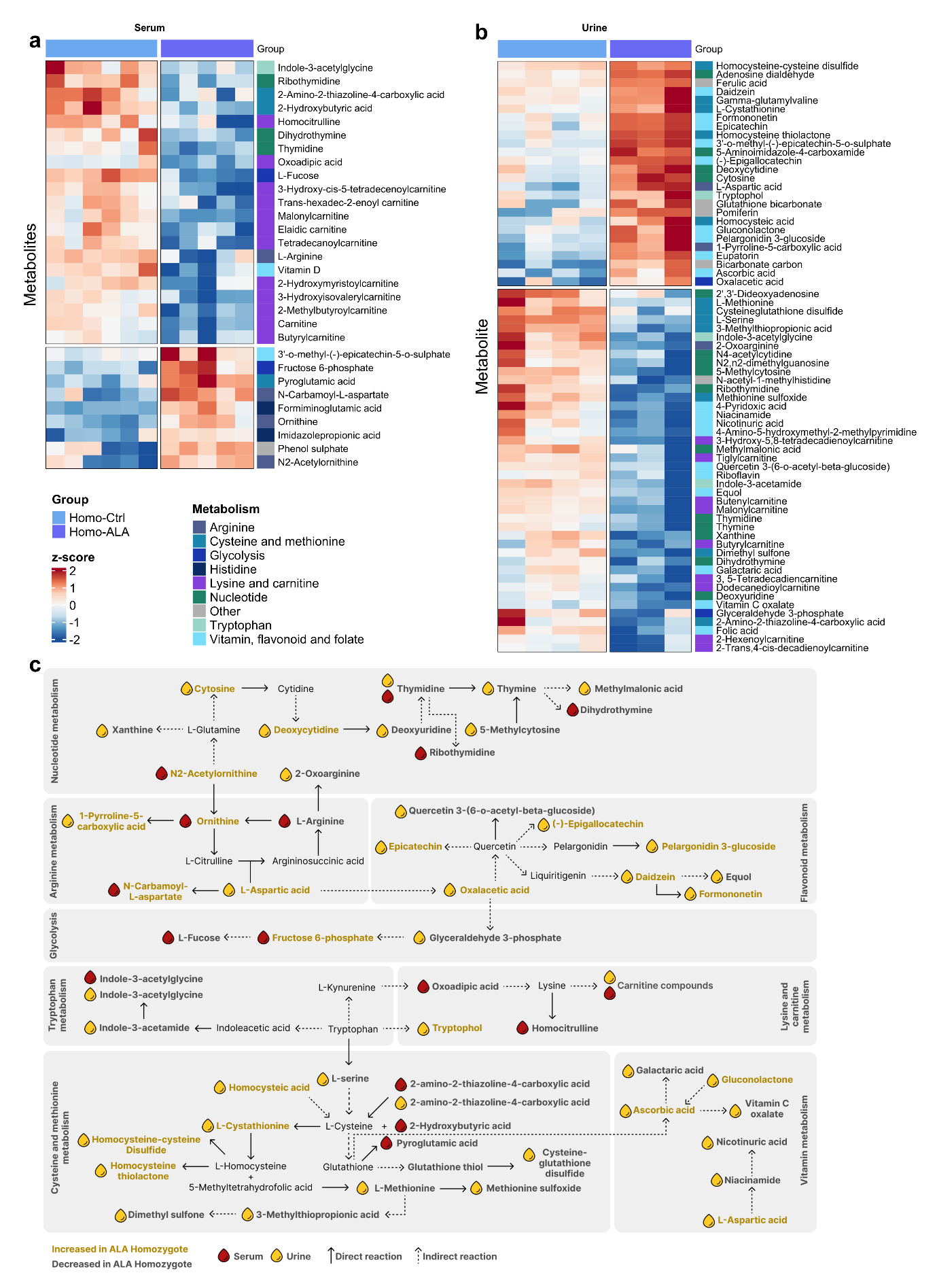

**Figure S24. Alpha-lipoic acid dietary supplementation markedly reshapes systemic and urinary metabolite profiles in homozygote *Slc7a9*^G105R^ mice.** Heatmap of differentially abundant (**a**) serum and (**b**) urine metabolites in homozygote alpha-lipoic acid (ALA)-treated group (Homo-ALA) compared to homozygote controls (Homo-Ctrl) mapped to metabolic pathways of interest. Data are colored according to changes in metabolite abundance (z-score) for each sample. (**c**) Metabolic pathway network showing differentially abundant metabolites mapped to their respective pathways according to the KEGG database, along with interactions across pathways. Arrows outline whether metabolites are connected by a direct (solid line) or indirect (dotted line) reaction. Icons mark metabolites that are significant in the blood (red color) or the urine (yellow color). Bold yellow text denotes that the metabolite is increased in ALA-treated homozygotes, and bold black text denotes that a metabolite is decreased in ALA-treated homozygotes. Metabolites without icons represent connecting metabolites that were not significant.

**Supplementary Figure S25**

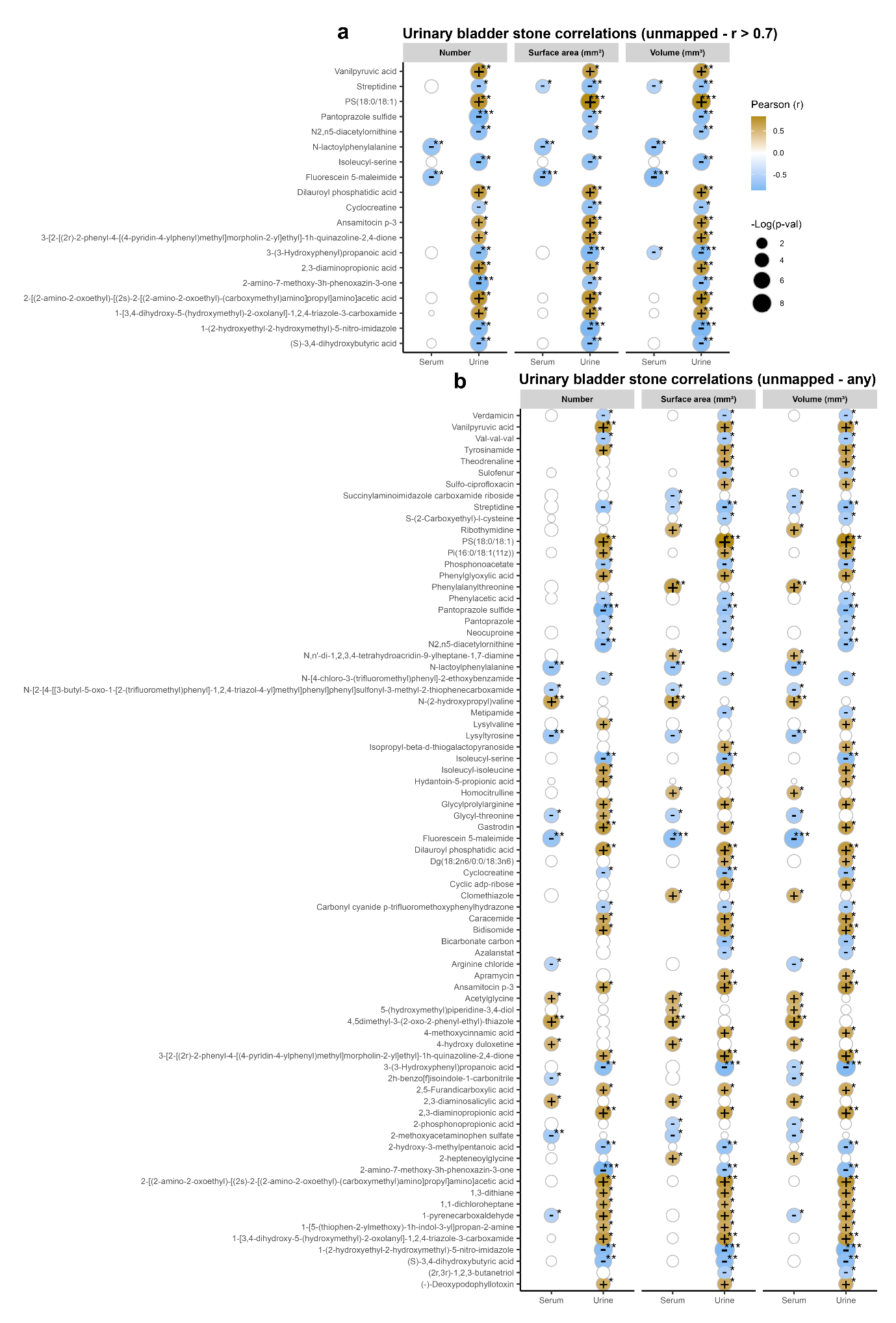

**Figure S25.** **Unmapped** **serum and urine metabolites correlate with the number, surface area, and volume of urinary bladder stones.** The analysis examines the relationship between serum or urine metabolites and the number, surface area, and volume of cystine stones. Empty circles with a grey outline in the plot indicate metabolites for which the Pearson correlation was not significant. Empty spots in the plot indicate that a metabolite was not detected in the serum or urine. (**a**) Metabolites that do not map to metabolic pathways of interest but have a correlation coefficient r > 0.7. (**b**) All metabolites that correlated with stone number, surface area, and volume but did not map to metabolic pathways of interest.

**Supplementary Figure S26**

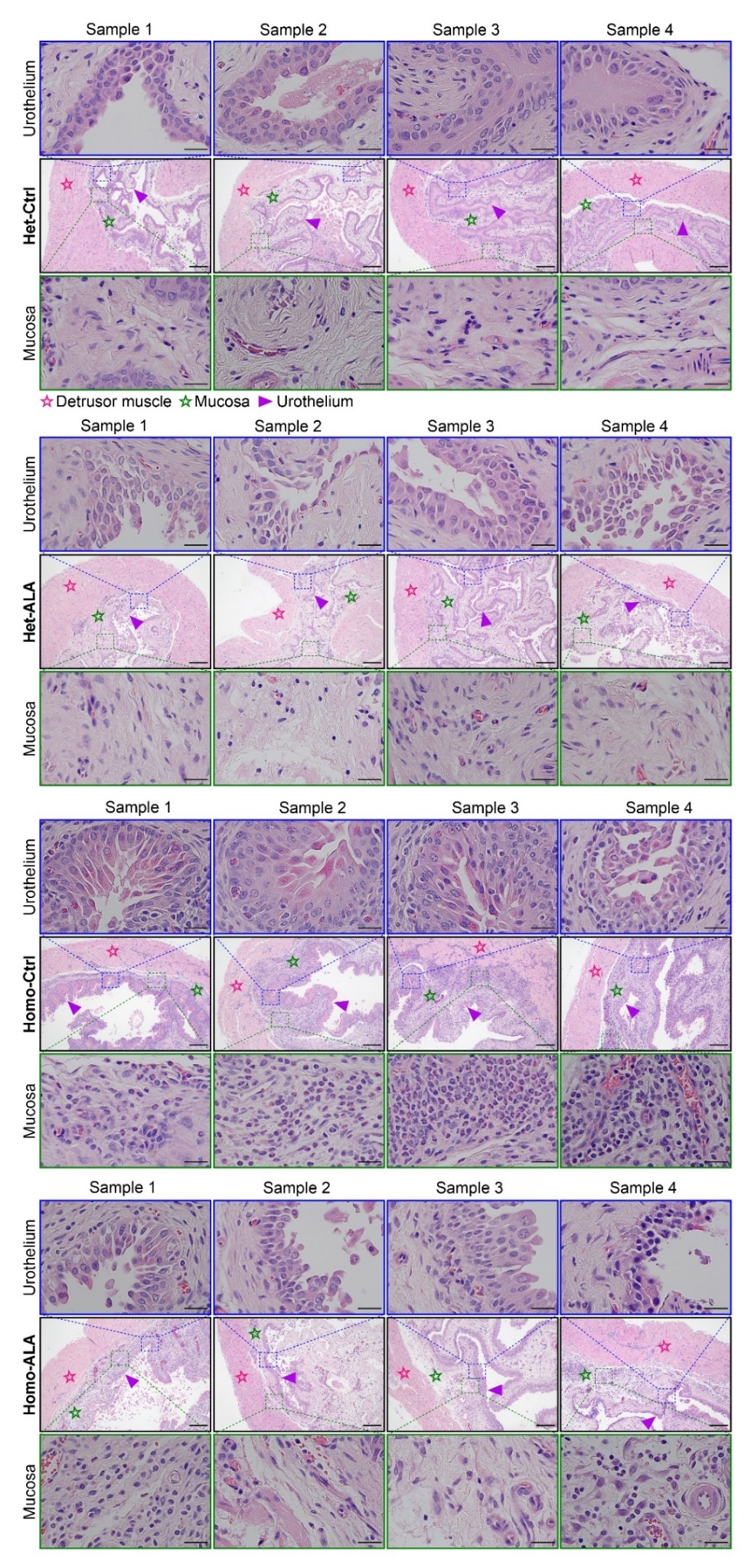

**Figure S26.** **Dietary supplementation with alpha-lipoic acid in established disease reduced cystine stone-induced urinary bladder inflammation but disrupted the urothelium**. Mice were fed an alpha-lipoic acid (ALA)-supplemented diet from 9 to 12 weeks of age (3-week treatment). Representative hematoxylin and eosin (H&E)-stained urinary bladder sections are shown from heterozygote *Slc7a9*^G105R^ controls (Het-Ctrl) and alpha-lipoic acid (ALA)-treated heterozygotes (Het-ALA), as well as homozygote controls (Homo-Ctrl) and ALA-treated homozygotes (Homo-ALA). Symbols ☆, ☆, and ► indicate the detrusor muscle, mucosa, and urothelium, respectively. Scale bars represent 200 µm for bladder sections and 25 µm for magnified views of the urothelium and mucosa. Each group indicates representative images from four independent mice.

**Supplementary tables**

**Supplementary Table S1.** Genomic targets and nucleotide sequences used to generate *Slc7a9*^G105R^ mice

| Nucleotide sequence | |
| --- | --- |
| Reference sequence | TGGACCTTTGGAGAGTGGAAAGTAGTTCTTAGGGTGAAGTTTTAATGGCTAGACTAGTACTGGGAATGCCTCCTTTGCATGGAGAAGCATTCCAGGAATCCCAGGTACTTGCTGGGGTGGGGACTGAACTTGAAGATCTGCTAAAGGTAGAGAGTATGGGGCGGGCAGGGGGTGGAGAAAGGGGAGTGCTCCCCCAAAAAGGCCCTAGGTGTTGGAACTACATGTTGCACCATCATGCCCAGCTGGGTTCCTTGCAGTGGGTGTGTGGTAAATAAGGGTGAATCTGACTGCAGTTAGAGGTCCAGCAGCCAGAACTGGGCTGTTTAGAATGTGAGCCCTCAAAGGAGGAAGCTCTGTGGAATAGGATGGAACCTCCTAAGGAGGGGGCTCCCTTCTGTGTTGCAGGCGCCCTGTGCTTTGCAGAGCTTGGCACAATGATTACCAAGTCAGGGGGTGAGTACCCCTATCTGATGGAGG**CCT**TT**GGC**CCCATCCCTGCCTACCTCTTCTCCTGGACCAGCCTGATCGTCATGAAGCCCTCATCCTTCGCCATCATCTGCCTCAGCTTTTCAGAGTATGTGTGTGCAGCCT |
| Forward primer | 5’-TGGACCTTTGGAGAGTGGAA-3’ |
| Reverse primer | 5’-AGGCTGCACACACATACTCT-3’ |
| Guide sequence | |
| *Slc7a9*^G105R^ crRNA | GTAGGCAGGGATGGG**GCC**AA AGG |
| Reverse complement | **CCT** TT**GGC**CCCATCCCTGCCTAC |
| *Slc7a9*^G105R^ ssOligo | ATGATGGCGAAGGATGAGGGCTTCATGACGATCAGGCTGGTCCAGGAGAAGAGGTAGGCAGGGATGGGGC**G**AA**AGG**CCTCCATCAGATAGGGGTACTCACCCCCTGACTTGGTAATCATTGTGCCAAGCTCTGCAAAGCAC |
| Reverse complement | GTGCTTTGCAGAGCTTGGCACAATGATTACCAAGTCAGGGGGTGAGTACCCCTATCTGATGGAGG**CCT**TT**CGC**CCCATCCCTGCCTACCTCTTCTCCTGGACCAGCCTGATCGTCATGAAGCCCTCATCCTTCGCCATCAT |

Reference base **G,** Mutation base **C**

**Supplementary Table S2.** Nutritional components of standard, cysteine-controlled (SF20-009), and alpha-lipoic-acid (ALA) supplement (SF21-078) diets

| Components | Standard diet (Breeding) | Cysteine-controlled diet | ALA supplement diet |
| --- | --- | --- | --- |
| Protein | 19.0% | 12.7% | 12.9% |
| Total Fat | 4.6% | 4.0% | 4.0% |
| Crude fiber | 5.2% | 4.7% | 4.7% |
| Total carbohydrate | 59.9% | 65.5% | 65.0% |
| Digestible energy | 14.2 MJ/Kg | 15.3 MJ/Kg | 15.3 MJ/Kg |
| % total calculated energy from protein | 23.0% | 14.0% | 14.0% |
| % total calculated energy from lipids | 12.0% | 9.5% | 9.5% |
| Valine | 0.87% | 0.80% | 0.77% |
| Leucine | 1.40% | 1.20% | 1.20% |
| Isoleucine | 0.80% | 0.62% | 0.62% |
| Threonine | 0.70% | 0.50% | 0.48% |
| Methionine | 0.30% | 0.37% | 0.37% |
| Cystine | 0.30% | 0.50% | 0.50% |
| Lysine | 0.90% | 1.1% | 1.1% |
| Phenylalanine | 0.90% | 0.62% | 0.62% |
| Tyrosine | 0.50% | 0.82% | 0.82% |
| Tryptophan | 0.20% | 0.12% | 0.12% |
| Histidine | 0.53% | 0.25% | 0.25% |
| L-cysteine HCL | - | 4.0 g/Kg | 4.0 g/Kg |
| Alpha-lipoic acid | - | - | 5.0 g/Kg |

**Supplementary Table S3.** List of up-regulated genes in homozygote *Slc7a9*^G105R^ mice compared to heterozygote controls (2-fold change, pooled sample; *n* = 10 per group)

| Gene Symbol | Description | RT^2^ Catalog | Fold change |
| --- | --- | --- | --- |
| *Il9* | Interleukin 9 | [PPM03110A](https://geneglobe.qiagen.com/search?cat=&q=PPM03110A) | 12.36 |
| *Tlr9* | Toll-like receptor 9 | [PPM04221A](https://geneglobe.qiagen.com/search?cat=&q=PPM04221A) | 12.26 |
| *Tnf* | Tumor necrosis factor | [PPM03113G](https://geneglobe.qiagen.com/search?cat=&q=PPM03113G) | 9.85 |
| *Tlr6* | Toll-like receptor 6 | [PPM04210B](https://geneglobe.qiagen.com/search?cat=&q=PPM04210B) | 9.40 |
| *Tlr5* | Toll-like receptor 5 | [PPM04206E](https://geneglobe.qiagen.com/search?cat=&q=PPM04206E) | 8.32 |
| *Il1rn* | Interleukin 1 receptor antagonist | [PPM03547B](https://geneglobe.qiagen.com/search?cat=&q=PPM03547B) | 6.17 |
| *C4b* | Complement component 4B | [PPM05780A](https://geneglobe.qiagen.com/search?cat=&q=PPM05780A) | 4.54 |
| *Cxcl1* | Chemokine (C-X-C motif) ligand 1 | [PPM03058C](https://geneglobe.qiagen.com/search?cat=&q=PPM03058C) | 4.02 |
| *Ccl25* | Chemokine (C-C motif) ligand 25 | [PPM02972F](https://geneglobe.qiagen.com/search?cat=&q=PPM02972F) | 2.65 |
| *Il7* | Interleukin 7 | [PPM03016C](https://geneglobe.qiagen.com/search?cat=&q=PPM03016C) | 2.37 |
| *Ccl1* | Chemokine (C-C motif) ligand 1 | [PPM03138C](https://geneglobe.qiagen.com/search?cat=&q=PPM03138C) | 2.25 |
| *Ccl11* | Chemokine (C-C motif) ligand 11 | [PPM02967G](https://geneglobe.qiagen.com/search?cat=&q=PPM02967G) | 2.25 |
| *Ccl22* | Chemokine (C-C motif) ligand 22 | [PPM02950B](https://geneglobe.qiagen.com/search?cat=&q=PPM02950B) | 2.25 |
| *Ccl24* | Chemokine (C-C motif) ligand 24 | [PPM03159F](https://geneglobe.qiagen.com/search?cat=&q=PPM03159F) | 2.25 |
| *Ccl7* | Chemokine (C-C motif) ligand 7 | [PPM02955B](https://geneglobe.qiagen.com/search?cat=&q=PPM02955B) | 2.25 |
| *Ccl8* | Chemokine (C-C motif) ligand 8 | [PPM03165A](https://geneglobe.qiagen.com/search?cat=&q=PPM03165A) | 2.25 |
| *Ccr4* | Chemokine (C-C motif) receptor 4 | [PPM03147A](https://geneglobe.qiagen.com/search?cat=&q=PPM03147A) | 2.25 |
| *Ccr7* | Chemokine (C-C motif) receptor 7 | [PPM03156G](https://geneglobe.qiagen.com/search?cat=&q=PPM03156G) | 2.25 |
| *Cd40lg* | CD40 ligand | [PPM03226C](https://geneglobe.qiagen.com/search?cat=&q=PPM03226C) | 2.25 |
| *Crp* | C-reactive protein, pentraxin-related | [PPM05334C](https://geneglobe.qiagen.com/search?cat=&q=PPM05334C) | 2.25 |
| *Cxcl11* | Chemokine (C-X-C motif) ligand 11 | [PPM03192C](https://geneglobe.qiagen.com/search?cat=&q=PPM03192C) | 2.25 |
| *Cxcl3* | Chemokine (C-X-C motif) ligand 3 | [PPM34590C](https://geneglobe.qiagen.com/search?cat=&q=PPM34590C) | 2.25 |
| *Cxcl5* | Chemokine (C-X-C motif) ligand 5 | [PPM02966F](https://geneglobe.qiagen.com/search?cat=&q=PPM02966F) | 2.25 |
| *Cxcr1* | Chemokine (C-X-C motif) receptor 1 | [PPM05308E](https://geneglobe.qiagen.com/search?cat=&q=PPM05308E) | 2.25 |
| *Cxcr2* | Chemokine (C-X-C motif) receptor 2 | [PPM03029A](https://geneglobe.qiagen.com/search?cat=&q=PPM03029A) | 2.25 |
| *Fasl* | Fas ligand (TNF superfamily, member 6) | [PPM02926E](https://geneglobe.qiagen.com/search?cat=&q=PPM02926E) | 2.25 |
| *Ifng* | Interferon gamma | [PPM03121A](https://geneglobe.qiagen.com/search?cat=&q=PPM03121A) | 2.25 |
| *Il10* | Interleukin 10 | [PPM03017C](https://geneglobe.qiagen.com/search?cat=&q=PPM03017C) | 2.25 |
| *Il17a* | Interleukin 17A | [PPM03023A](https://geneglobe.qiagen.com/search?cat=&q=PPM03023A) | 2.25 |
| *Il22* | Interleukin 22 | [PPM05481A](https://geneglobe.qiagen.com/search?cat=&q=PPM05481A) | 2.25 |
| *Il5* | Interleukin 5 | [PPM03014F](https://geneglobe.qiagen.com/search?cat=&q=PPM03014F) | 2.25 |
| *Il6* | Interleukin 6 | [PPM03015A](https://geneglobe.qiagen.com/search?cat=&q=PPM03015A) | 2.25 |
| *Lta* | Lymphotoxin A | [PPM03114A](https://geneglobe.qiagen.com/search?cat=&q=PPM03114A) | 2.25 |
| *Ltb* | Lymphotoxin B | [PPM03119A](https://geneglobe.qiagen.com/search?cat=&q=PPM03119A) | 2.25 |
| *Cxcl2* | Chemokine (C-X-C motif) ligand 2 | [PPM02969F](https://geneglobe.qiagen.com/search?cat=&q=PPM02969F) | 2.22 |
| *Kng1* | Kininogen 1 | [PPM60015A](https://geneglobe.qiagen.com/search?cat=&q=PPM60015A) | 2.00 |

**Supplementary Table S4.** List of down-regulated genes in homozygote *Slc7a9*^G105R^ mice compared to heterozygote controls (2-fold change, pooled sample; *n* = 10 per group)

| Gene Symbol | Description | RT^2^ Catalog | Fold change |
| --- | --- | --- | --- |
| *Tlr2* | Toll-like receptor 2 | [PPM04220B](https://geneglobe.qiagen.com/search?cat=&q=PPM04220B) | -8.15 |
| *Il23a* | Interleukin 23, alpha subunit p19 | [PPM03763F](https://geneglobe.qiagen.com/search?cat=&q=PPM03763F) | -3.28 |
| *Il1a* | Interleukin 1 alpha | [PPM03010F](https://geneglobe.qiagen.com/search?cat=&q=PPM03010F) | -3.14 |
| *Ccl2* | Chemokine (C-C motif) ligand 2 | [PPM03151G](https://geneglobe.qiagen.com/search?cat=&q=PPM03151G) | -2.94 |
| *Ccl19* | Chemokine (C-C motif) ligand 19 | [PPM03157C](https://geneglobe.qiagen.com/search?cat=&q=PPM03157C) | -2.63 |
| *Il23r* | Interleukin 23 receptor | [PPM33761A](https://geneglobe.qiagen.com/search?cat=&q=PPM33761A) | -2.59 |
| *Ccr3* | Chemokine (C-C motif) receptor 3 | [PPM03173A](https://geneglobe.qiagen.com/search?cat=&q=PPM03173A) | -2.18 |
| *Tlr7* | Toll-like receptor 7 | [PPM04208A](https://geneglobe.qiagen.com/search?cat=&q=PPM04208A) | -2.17 |
| *Ccl12* | Chemokine (C-C motif) ligand 12 | [PPM02977E](https://geneglobe.qiagen.com/search?cat=&q=PPM02977E) | -2.08 |

**Supplementary Table S5.** List of up-regulated genes in antibiotic-treated homozygote *Slc7a9*^G105R^ mice (Homo + Abx) compared to homozygote *Slc7a9*^G105R^ vehicle-treated controls (Homo + Veh) (2-fold change, pooled sample; *n* = 10 per group)

| Gene Symbol | Description | RT^2^ Catalog | Fold change |
| --- | --- | --- | --- |
| *Tlr2* | Toll-like receptor 2 | [PPM04220B](https://geneglobe.qiagen.com/search?cat=&q=PPM04220B) | 9.15 |
| *Ccl2* | Chemokine (C-C motif) ligand 2 | [PPM03151G](https://geneglobe.qiagen.com/search?cat=&q=PPM03151G) | 3.75 |
| *Ccl11* | Chemokine (C-C motif) ligand 11 | [PPM02967G](https://geneglobe.qiagen.com/search?cat=&q=PPM02967G) | 3.61 |
| *Ltb* | Lymphotoxin B | [PPM03119A](https://geneglobe.qiagen.com/search?cat=&q=PPM03119A) | 2.77 |
| *Ccl5* | Chemokine (C-C motif) ligand 5 | [PPM02960F](https://geneglobe.qiagen.com/search?cat=&q=PPM02960F) | 2.69 |
| *Ccr3* | Chemokine (C-C motif) receptor 3 | [PPM03173A](https://geneglobe.qiagen.com/search?cat=&q=PPM03173A) | 2.41 |
| *Ripk2* | Receptor (TNFRSF)-interacting serine-threonine kinase 2 | [PPM06268A](https://geneglobe.qiagen.com/search?cat=&q=PPM06268A) | 2.38 |
| *Il23a* | Interleukin 23, alpha subunit p19 | [PPM03763F](https://geneglobe.qiagen.com/search?cat=&q=PPM03763F) | 2.29 |
| *Kng1* | Kininogen 1 | [PPM60015A](https://geneglobe.qiagen.com/search?cat=&q=PPM60015A) | 2.26 |
| *Tlr7* | Toll-like receptor 7 | [PPM04208A](https://geneglobe.qiagen.com/search?cat=&q=PPM04208A) | 2.10 |
| *Tlr4* | Toll-like receptor 4 | [PPM04207F](https://geneglobe.qiagen.com/search?cat=&q=PPM04207F) | 2.09 |

**Supplementary Table S6.** List of down-regulated genes in antibiotic-treated homozygote *Slc7a9*^G105R^ mice (Homo + Abx) compared to homozygote *Slc7a9*^G105R^ vehicle-treated controls (Homo + Veh) (2-fold change, pooled sample; *n* = 10 per group)

| Gene Symbol | Description | RT^2^ Catalog | Fold change |
| --- | --- | --- | --- |
| *Il9* | Interleukin 9 | [PPM03110A](https://geneglobe.qiagen.com/search?cat=&q=PPM03110A) | -17.02 |
| *Ccl17* | Chemokine (C-C motif) ligand 17 | [PPM02963B](https://geneglobe.qiagen.com/search?cat=&q=PPM02963B) | -15.21 |
| *Il7* | Interleukin 7 | [PPM03016C](https://geneglobe.qiagen.com/search?cat=&q=PPM03016C) | -6.04 |
| *Cxcl1* | Chemokine (C-X-C motif) ligand 1 | [PPM03058C](https://geneglobe.qiagen.com/search?cat=&q=PPM03058C) | -5.53 |
| *Ccl12* | Chemokine (C-C motif) ligand 12 | [PPM02977E](https://geneglobe.qiagen.com/search?cat=&q=PPM02977E) | -3.10 |
| *Ccl20* | Chemokine (C-C motif) ligand 20 | [PPM03142B](https://geneglobe.qiagen.com/search?cat=&q=PPM03142B) | -3.10 |
| *Ccl24* | Chemokine (C-C motif) ligand 24 | [PPM03159F](https://geneglobe.qiagen.com/search?cat=&q=PPM03159F) | -3.10 |
| *Ccl7* | Chemokine (C-C motif) ligand 7 | [PPM02955B](https://geneglobe.qiagen.com/search?cat=&q=PPM02955B) | -3.10 |
| *Ccl8* | Chemokine (C-C motif) ligand 8 | [PPM03165A](https://geneglobe.qiagen.com/search?cat=&q=PPM03165A) | -3.10 |
| *Ccr4* | Chemokine (C-C motif) receptor 4 | [PPM03147A](https://geneglobe.qiagen.com/search?cat=&q=PPM03147A) | -3.10 |
| *Cd40lg* | CD40 ligand | [PPM03226C](https://geneglobe.qiagen.com/search?cat=&q=PPM03226C) | -3.10 |
| *Cxcl3* | Chemokine (C-X-C motif) ligand 3 | [PPM34590C](https://geneglobe.qiagen.com/search?cat=&q=PPM34590C) | -3.10 |
| *Cxcl5* | Chemokine (C-X-C motif) ligand 5 | [PPM02966F](https://geneglobe.qiagen.com/search?cat=&q=PPM02966F) | -3.10 |
| *Cxcr1* | Chemokine (C-X-C motif) receptor 1 | [PPM05308E](https://geneglobe.qiagen.com/search?cat=&q=PPM05308E) | -3.10 |
| *Cxcr2* | Chemokine (C-X-C motif) receptor 2 | [PPM03029A](https://geneglobe.qiagen.com/search?cat=&q=PPM03029A) | -3.10 |
| *Fasl* | Fas ligand (TNF superfamily, member 6) | [PPM02926E](https://geneglobe.qiagen.com/search?cat=&q=PPM02926E) | -3.10 |
| *Ifng* | Interferon gamma | [PPM03121A](https://geneglobe.qiagen.com/search?cat=&q=PPM03121A) | -3.10 |
| *Il10* | Interleukin 10 | [PPM03017C](https://geneglobe.qiagen.com/search?cat=&q=PPM03017C) | -3.10 |
| *Il23r* | Interleukin 23 receptor | [PPM33761A](https://geneglobe.qiagen.com/search?cat=&q=PPM33761A) | -3.10 |
| *Il6* | Interleukin 6 | [PPM03015A](https://geneglobe.qiagen.com/search?cat=&q=PPM03015A) | -3.10 |
| *Lta* | Lymphotoxin A | [PPM03114A](https://geneglobe.qiagen.com/search?cat=&q=PPM03114A) | -3.10 |
| *Tnfsf14* | Tumor necrosis factor (ligand) superfamily, member 14 | [PPM03553F](https://geneglobe.qiagen.com/search?cat=&q=PPM03553F) | -3.10 |
| *Tlr3* | Toll-like receptor 3 | [PPM04216B](https://geneglobe.qiagen.com/search?cat=&q=PPM04216B) | -2.54 |
| *Gusb* | Glucuronidase, beta | [PPM05490C](https://geneglobe.qiagen.com/search?cat=&q=PPM05490C) | -2.39 |
| *Il17a* | Interleukin 17A | [PPM03023A](https://geneglobe.qiagen.com/search?cat=&q=PPM03023A) | -2.22 |
| *Tlr6* | Toll-like receptor 6 | [PPM04210B](https://geneglobe.qiagen.com/search?cat=&q=PPM04210B) | -2.10 |
| *Ccl22* | Chemokine (C-C motif) ligand 22 | [PPM02950B](https://geneglobe.qiagen.com/search?cat=&q=PPM02950B) | -2.06 |
| *Csf1* | Colony stimulating factor 1 (macrophage) | [PPM03116C](https://geneglobe.qiagen.com/search?cat=&q=PPM03116C) | -2.03 |

**Supplementary Table S7.** List of up-regulated genes in alpha-lipoic acid (ALA) supplemented diet homozygote *Slc7a9*^G105R^ mice (Homo-ALA) compared to homozygote *Slc7a9*^G105R^ controls (Homo-Ctrl) (2-fold change, pooled sample; *n* = 10 per group)

| Gene Symbol | Description | RT^2^ Catalog | Fold change |
| --- | --- | --- | --- |
| *Tlr2* | Toll-like receptor 2 | [PPM04220B](https://geneglobe.qiagen.com/search?cat=&q=PPM04220B) | 5.94 |
| *Ccl5* | Chemokine (C-C motif) ligand 5 | [PPM02960F](https://geneglobe.qiagen.com/search?cat=&q=PPM02960F) | 5.35 |
| *Ccl12* | Chemokine (C-C motif) ligand 12 | [PPM02977E](https://geneglobe.qiagen.com/search?cat=&q=PPM02977E) | 4.33 |
| *C3* | Complement component 3 | [PPM04471E](https://geneglobe.qiagen.com/search?cat=&q=PPM04471E) | 3.57 |
| *Ccl2* | Chemokine (C-C motif) ligand 2 | [PPM03151G](https://geneglobe.qiagen.com/search?cat=&q=PPM03151G) | 2.53 |
| *Ripk2* | Receptor (TNFRSF)-interacting serine-threonine kinase 2 | [PPM06268A](https://geneglobe.qiagen.com/search?cat=&q=PPM06268A) | 2.29 |
| *Ccl8* | Chemokine (C-C motif) ligand 8 | [PPM03165A](https://geneglobe.qiagen.com/search?cat=&q=PPM03165A) | 2.15 |

**Table S8.** List of down-regulated genes in alpha-lipoic acid (ALA) supplemented diet homozygote *Slc7a9*^G105R^ mice (Homo-ALA) compared to homozygote *Slc7a9*^G105R^ controls (Homo-Ctrl) (2-fold change, pooled sample; *n* = 10 per group)

| Gene Symbol | Description | RT^2^ Catalog | Fold change |
| --- | --- | --- | --- |
| *Cxcl1* | Chemokine (C-X-C motif) ligand 1 | [PPM03058C](https://geneglobe.qiagen.com/search?cat=&q=PPM03058C) | -5.12 |
| *Il7* | Interleukin 7 | [PPM03016C](https://geneglobe.qiagen.com/search?cat=&q=PPM03016C) | -4.26 |
| *Il9* | Interleukin 9 | [PPM03110A](https://geneglobe.qiagen.com/search?cat=&q=PPM03110A) | -3.60 |
| *Cxcr4* | Chemokine (C-X-C motif) receptor 4 | [PPM03149E](https://geneglobe.qiagen.com/search?cat=&q=PPM03149E) | -3.51 |
| *Tnf* | Tumor necrosis factor | [PPM03113G](https://geneglobe.qiagen.com/search?cat=&q=PPM03113G) | -3.27 |
| *Ccl11* | Chemokine (C-C motif) ligand 11 | [PPM02967G](https://geneglobe.qiagen.com/search?cat=&q=PPM02967G) | -3.17 |
| *Ccl20* | Chemokine (C-C motif) ligand 20 | [PPM03142B](https://geneglobe.qiagen.com/search?cat=&q=PPM03142B) | -3.17 |
| *Ccl24* | Chemokine (C-C motif) ligand 24 | [PPM03159F](https://geneglobe.qiagen.com/search?cat=&q=PPM03159F) | -3.17 |
| *Ccr4* | Chemokine (C-C motif) receptor 4 | [PPM03147A](https://geneglobe.qiagen.com/search?cat=&q=PPM03147A) | -3.17 |
| *Ccl1* | Chemokine (C-C motif) ligand 1 | [PPM03138C](https://geneglobe.qiagen.com/search?cat=&q=PPM03138C) | -3.17 |
| *Cd40lg* | CD40 ligand | [PPM03226C](https://geneglobe.qiagen.com/search?cat=&q=PPM03226C) | -3.17 |
| *Crp* | C-reactive protein, pentraxin-related | [PPM05334C](https://geneglobe.qiagen.com/search?cat=&q=PPM05334C) | -3.17 |
| *Cxcl11* | Chemokine (C-X-C motif) ligand 11 | [PPM03192C](https://geneglobe.qiagen.com/search?cat=&q=PPM03192C) | -3.17 |
| *Cxcl3* | Chemokine (C-X-C motif) ligand 3 | [PPM34590C](https://geneglobe.qiagen.com/search?cat=&q=PPM34590C) | -3.17 |
| *Cxcl5* | Chemokine (C-X-C motif) ligand 5 | [PPM02966F](https://geneglobe.qiagen.com/search?cat=&q=PPM02966F) | -3.17 |
| *Cxcl9* | Chemokine (C-X-C motif) ligand 9 | [PPM02973B](https://geneglobe.qiagen.com/search?cat=&q=PPM02973B) | -3.17 |
| *Cxcr1* | Chemokine (C-X-C motif) receptor 1 | [PPM05308E](https://geneglobe.qiagen.com/search?cat=&q=PPM05308E) | -3.17 |
| *Cxcr2* | Chemokine (C-X-C motif) receptor 2 | [PPM03029A](https://geneglobe.qiagen.com/search?cat=&q=PPM03029A) | -3.17 |
| *Fasl* | Fas ligand (TNF superfamily, member 6) | [PPM02926E](https://geneglobe.qiagen.com/search?cat=&q=PPM02926E) | -3.17 |
| *Ifng* | Interferon gamma | [PPM03121A](https://geneglobe.qiagen.com/search?cat=&q=PPM03121A) | -3.17 |
| *Il17a* | Interleukin 17A | [PPM03023A](https://geneglobe.qiagen.com/search?cat=&q=PPM03023A) | -3.17 |
| *Il22* | Interleukin 22 | [PPM05481A](https://geneglobe.qiagen.com/search?cat=&q=PPM05481A) | -3.17 |
| *Il23a* | Interleukin 23, alpha subunit p19 | [PPM03763F](https://geneglobe.qiagen.com/search?cat=&q=PPM03763F) | -3.17 |
| *Il23r* | Interleukin 23 receptor | [PPM33761A](https://geneglobe.qiagen.com/search?cat=&q=PPM33761A) | -3.17 |
| *Il6* | Interleukin 6 | [PPM03015A](https://geneglobe.qiagen.com/search?cat=&q=PPM03015A) | -3.17 |
| *Kng1* | Kininogen 1 | [PPM60015A](https://geneglobe.qiagen.com/search?cat=&q=PPM60015A) | -3.17 |
| *Lta* | Lymphotoxin A | [PPM03114A](https://geneglobe.qiagen.com/search?cat=&q=PPM03114A) | -3.17 |
| *Tnfsf14* | Tumor necrosis factor (ligand) superfamily, member 14 | [PPM03553F](https://geneglobe.qiagen.com/search?cat=&q=PPM03553F) | -3.17 |
| *Ccr2* | Chemokine (C-C motif) receptor 2 | [PPM03176A](https://geneglobe.qiagen.com/search?cat=&q=PPM03176A) | -3.14 |
| *Gusb* | Glucuronidase, beta | [PPM05490C](https://geneglobe.qiagen.com/search?cat=&q=PPM05490C) | -3.10 |
| *Cxcl2* | Chemokine (C-X-C motif) ligand 2 | [PPM02969F](https://geneglobe.qiagen.com/search?cat=&q=PPM02969F) | -2.95 |
| *Cd40* | CD40 antigen | [PPM03426C](https://geneglobe.qiagen.com/search?cat=&q=PPM03426C) | -2.62 |
| *Tlr6* | Toll-like receptor 6 | [PPM04210B](https://geneglobe.qiagen.com/search?cat=&q=PPM04210B) | -2.55 |
| *Ccl25* | Chemokine (C-C motif) ligand 25 | [PPM02972F](https://geneglobe.qiagen.com/search?cat=&q=PPM02972F) | -2.49 |
| *Ccl17* | Chemokine (C-C motif) ligand 17 | [PPM02963B](https://geneglobe.qiagen.com/search?cat=&q=PPM02963B) | -2.48 |
| *Tlr3* | Toll-like receptor 3 | [PPM04216B](https://geneglobe.qiagen.com/search?cat=&q=PPM04216B) | -2.35 |
| *Csf1* | Colony stimulating factor 1 (macrophage) | [PPM03116C](https://geneglobe.qiagen.com/search?cat=&q=PPM03116C) | -2.31 |

**Supplementary References**

1. Wu N, Liu B, Du H, et al. The Progress of CRISPR/Cas9-Mediated Gene Editing in Generating Mouse/Zebrafish Models of Human Skeletal Diseases. *Computational and Structural Biotechnology Journal*. 2019;17:954-962.
2. Scarfe L, Schock-Kusch D, Ressel L, et al. Transdermal Measurement of Glomerular Filtration Rate in Mice. *J Vis Exp*. 2018;(140).
3. Janssen PM, Murray JD, Schill KE, et al. Prednisolone attenuates improvement of cardiac and skeletal contractile function and histopathology by lisinopril and spironolactone in the mdx mouse model of Duchenne muscular dystrophy. *PLoS One*. 2014;9(2):e88360.
4. Rafael-Fortney JA, Chimanji NS, Schill KE, et al. Early treatment with lisinopril and spironolactone preserves cardiac and skeletal muscle in Duchenne muscular dystrophy mice. *Circulation*. 2011;124(5):582-8.
5. Alpert AJ. Hydrophilic-interaction chromatography for the separation of peptides, nucleic acids and other polar compounds. *Journal of Chromatography A*. 1990;499:177-196.
6. Wypych TP, Pattaroni C, Perdijk O, et al. Microbial metabolism of l-tyrosine protects against allergic airway inflammation. *Nature Immunology*. 2021;22(3):279-286.
7. A quality control tool for high throughput sequence data. Version 0.1. *Babraham Bioinformatics*; 2010.
8. Langmead B, Salzberg SL. Fast gapped-read alignment with Bowtie 2. *Nature Methods*. 2012;9(4):357-359.
9. Chen S. Ultrafast one-pass FASTQ data preprocessing, quality control, and deduplication using fastp. *iMeta*. 2023;2(2):e107.
10. Blanco-Míguez A, Beghini F, Cumbo F, et al. Extending and improving metagenomic taxonomic profiling with uncharacterized species using MetaPhlAn 4. *Nature Biotechnology*. 2023;41(11):1633-1644.
11. Lin H, Peddada SD. Multigroup analysis of compositions of microbiomes with covariate adjustments and repeated measures. *Nature Methods*. 2024;21(1):83-91.
12. Benjamini Y, Hochberg Y. Controlling the False Discovery Rate: A Practical and Powerful Approach to Multiple Testing. *Journal of the Royal Statistical Society Series B (Methodological)*. 1995;57(1):289-300.
13. Ordination methods, diversity analysis and other functions for community and vegetation ecologists. Version R package version 2.6-6.1. vegan: *Community Ecology Package*; 2024.
14. Anderson MJ. Distance-Based Tests for Homogeneity of Multivariate Dispersions. *Biometrics*. 2005;62(1):245-253.
15. Beghini F, McIver LJ, Blanco-Míguez A, et al. Integrating taxonomic, functional, and strain-level profiling of diverse microbial communities with bioBakery 3. *eLife*. 2021;10:e65088.
16. Caspi R, Billington R, Keseler IM, et al. The MetaCyc database of metabolic pathways and enzymes - a 2019 update. *Nucleic Acids Research*. 2019;48(D1):D445-D453.
17. Mallick H, Rahnavard A, McIver LJ, et al. Multivariable association discovery in population-scale meta-omics studies. *PLoS Comput Biol*. 2021;17(11):e1009442.
18. mia: Microbiome analysis. Version 1.10.0. *Bioconductor*; 2023.
19. Microbiome Analysis Plotting and Visualization. Version 1.10.0. *Bioconductor*; 2023.
